# zAcademy: an open-source programmable conditioning arena reveals sex- and genotype-dependent avoidance learning in adult zebrafish

**DOI:** 10.64898/2026.09.11.750830

**Authors:** Shipei Huang, Baixi He, Yijie Geng

## Abstract

Zebrafish are widely used to study learning and memory. Here, we describe zAcademy, a programmable conditioning arena released in full as open hardware and open software. zAcademy combines a 3D-printed dual-chamber arena, bottom-mounted LED matrices that generate visual patterns under software control, graphite rod electrodes gated by an Arduino-controlled relay, an overhead infrared camera, and a position-based analysis workflow. In one 180-min session, adult zebrafish were classically conditioned to associate a checkerboard light pattern beneath the arena (conditioned stimulus, CS) with a mild shock (unconditioned stimulus, US). Conditioned avoidance was assessed in six unreinforced CS-alone probes: one immediately after training to test learning, the remaining five at 30-min intervals to test memory. Spatial preference was quantified as a performance index (PI; occupancy of the dark zone). Wild-type fish shifted toward the dark zone immediately after training, whereas *shank3* KO fish did not. The wild-type response was carried primarily by females rather than males. In addition, among initial responders, females sustained elevated PI at more later probes. zAcademy provides an inexpensive, fully documented, and readily modified platform for comparing experience-dependent behavioral change across genotypes and sexes.

## INTRODUCTION

Learning and memory enable animals to adapt behavior on the basis of prior experience, supporting survival, environmental adaptation, and flexible nervous system function^1,2^. Impairments in these processes are central to many neurological and neurodevelopmental disorders, which makes experimental models of experience-dependent behavioral change important for both mechanistic neuroscience and translational research^3^. Because learning and memory are ordinarily assessed through measurable behavioral change following controlled training, robust paradigms require biologically meaningful stimuli, reproducible stimulus delivery, and quantitative measures of behavioral response over time.

Zebrafish (*Danio rerio*) have emerged as a powerful vertebrate model for behavioral neuroscience because they combine genetic tractability, experimental scalability, and quantifiable behavioral responses. They are small, inexpensive to maintain, highly fecund, rapidly developing, externally fertilized, and optically transparent during early development, making them well suited to scalable and tightly controlled laboratory studies^4,5^. Their relevance to human biology is supported by genome-scale conservation, with approximately 70% of human genes having at least one zebrafish ortholog^6^. Zebrafish also exhibit a broad repertoire of measurable behaviors, including locomotion, exploration, anxiety-like responses, social behavior, visually guided behavior, and learning-related responses^4,7–10^.

A wide range of assays now covers most forms of zebrafish learning. Non-associative learning is measured as habituation of the acoustic startle or dark-flash response in larvae^11^; because these assays run in multiwell plates at high throughput, they have supported forward genetic screens that recovered more than a dozen habituation mutants, including *pappaa* and *cacna2d3*^12,13^. Associative learning in adults is assessed by classical fear conditioning, in which a visual or olfactory conditioned stimulus is paired with shock or alarm substance^14^; by conditioned place avoidance and conditioned place preference^15,16^; by one-trial inhibitory avoidance^17^; and by active avoidance in a shuttle box, a paradigm used in an early molecular study of memory consolidation in zebrafish^18^. Spatial and working memory are assessed in the T-maze^19^ and the free-movement pattern Y-maze^20^, recognition memory in novel object tests^21^, and social transmission in observational learning paradigms^22^. Within this landscape, automated platforms that combine programmed visual stimulation, electrical stimulation, and continuous video tracking occupy a distinct niche, allowing acquisition and subsequent expression of a conditioned response to be measured repeatedly in the same animal without rehandling^23,24^.

These assays differ substantially in hardware, procedure, and the questions they have been used to ask. While larval habituation assays have been extensively applied to mutants, automated associative conditioning in juveniles and adults has been conducted only in wild-type animals^23,24^. Sex is likewise infrequently analyzed. Adult studies commonly pool males and females or limit to one sex, while larval and juvenile paradigms preclude the comparison altogether, because fish at these stages are not yet sexually differentiated^25^. Where sex has been examined in adults, the findings vary: no sex difference was found in a Y-maze or Pavlovian fear conditioning task despite higher anxiety-like behavior in females^26^, females outperformed males in a five-day T-maze task^27^, and an appetitive place preference assay observed a sex-by-personality interaction without a main effect of sex^28^. These studies differ in reinforcer valence, stimulus modality, and retention interval, which may account for some of the discrepancy. Reproducibility of the apparatus itself is a further constraint: the automated system of Valente et al. was not released as modular open hardware^24^, and the conditioned place avoidance protocol of Palumbo et al., optimized for juveniles, uses a fixed-output operational amplifier circuit in which changing shock intensity requires manually substituting resistor values^23^.

To address these constraints we developed zAcademy, a programmable dual-chamber conditioning platform for adult zebrafish that integrates bottom-projected checkerboard visual stimulation, relay-switched electrical stimulation with software-selectable voltage, infrared video recording, and position-based behavioral analysis. The complete design, including arena STL files, Arduino firmware, automated recording scripts, tracking workflow, and analysis code, is released openly. Using zAcademy we implemented a continuous three-hour protocol combining baseline assessment, visual–shock conditioning, and six repeated unreinforced probe windows within a single experiment, quantifying spatial preference as a performance index (PI) based on occupancy of the dark versus checkerboard-illuminated region. We applied the platform to wild-type zebrafish and to a *shank3* KO line, and analyzed acquisition and retention separately in males and females.

## RESULTS

### Design and construction of zAcademy

zAcademy integrates stimulus delivery, electrical stimulation, and behavioral recording within a single enclosed setup (Figure 1). The behavioral arena was designed in SelfCAD and fabricated on a dual-nozzle 3D printer, allowing multi-material printing within one object. The upper walls were printed in black PLA to reduce internal reflection and maintain a dark visual environment during conditioning, while the chamber base was printed in white PLA to transmit light from the LED matrix beneath. Water-soluble PVA supported the bottom opening during printing and was dissolved away afterward, avoiding mechanical damage during support removal. The arena contains two identical chambers, each 192 mm long, 62 mm wide, and 120 mm deep, allowing two fish (one in each chamber) to be run in parallel.

**Figure 1.**
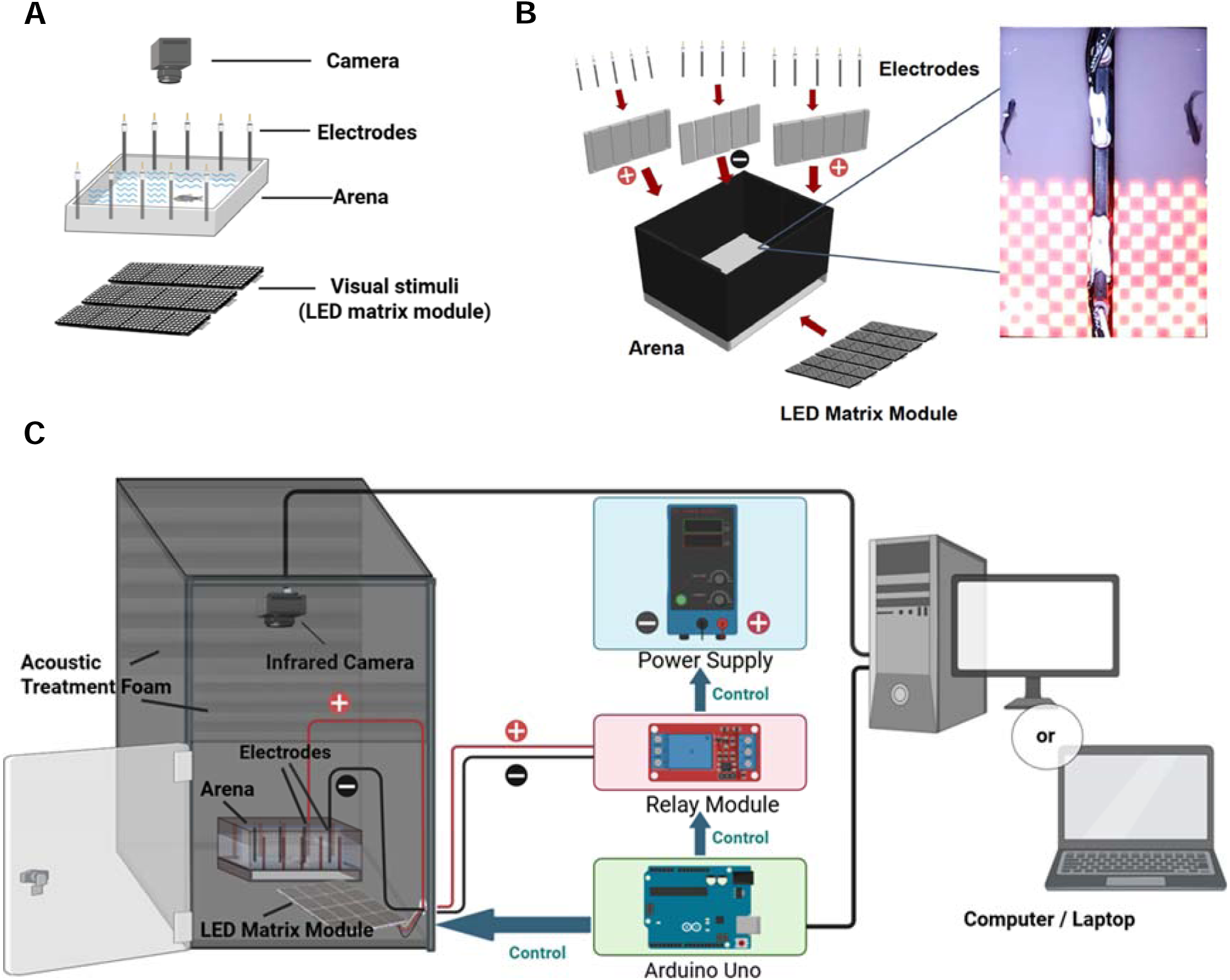
Setup of the zAcademy system. (**A**) Experimental setup. The visual stimulus is presented on an LED matrix module beneath a 3D-printed arena. Electrical stimulation across the arena is delivered by graphite rod electrodes. (**B**) Arena assembly and component integration. Graphite rod electrodes were positioned along the chamber walls to deliver electrical stimulation, while the LED matrix module was mounted beneath the arena floor to provide programmable visual cues. The inset image shows the checkerboard stimulus projected through the translucent base of the arena. (**C**) Final assembly and system-level architecture. The complete setup integrates a dual-chamber behavioral arena, bottom-mounted LED matrix module, graphite rod electrodes, overhead infrared camera, Arduino Uno, relay module, external power supply, and computer interface. The Arduino coordinates visual stimulus presentation through the LED matrix and controls the relay module for programmed electrical stimulation. The external power supply provides the stimulation voltage, while the infrared camera records top-down behavioral video.

Visual stimuli were delivered from below through the translucent chamber floor by MAX7219-based 32 × 8 dot matrix LED modules, each built from four integrated 8 × 8 panels (Figure 1A). The matrix was addressed as two independently controlled regions, designated Area 1 and Area 2, corresponding to the two halves of each chamber. Because the display is addressed at the pixel level, the spatial pattern, its position, and its timing are all defined in firmware rather than in hardware, and the LED assembly remains physically separated from the water.

Electrical stimulation was delivered through graphite rod electrodes (6 mm diameter, 90 mm length) integrated into the arena walls. The two chambers share a central row of negative electrodes along the divider, while each chamber carries a separate row of positive electrodes along its outer wall, so that current passes across the width of each chamber (Figure 1B). Stimulation was supplied by an external DC bench power supply switched by a single-channel relay under Arduino control. Because the bench supply sets the stimulation voltage and the relay only gates it, shock amplitude is adjusted by changing a setting on the supply rather than by modifying the circuit, while the relay keeps the low-voltage control electronics electrically isolated from the stimulation pathway (Figure 1C). For the experiments reported here the supply was set to 9.5 V with a 0.200 A compliance limit, and delivery was verified before every session by measuring 9.4–9.5 V across each electrode pair with a multimeter.

An Arduino Uno served as the central controller, coordinating LED pattern presentation, relay activation, and protocol timing, with a TM1637 four-digit display mounted outside the enclosure reporting elapsed protocol time. Behavior was recorded from above by a 38 × 38 mm infrared-sensitive camera with an automatic IR-CUT filter and integrated 850 nm LEDs, which engaged automatically during dark periods and disengaged when the LED matrix was illuminated. Because zebrafish visual sensitivity extends into the deep red and the optomotor response threshold in adults has been estimated at approximately 840 nm^29^, the 850 nm illumination was not assumed to be invisible to the fish; it was instead held constant as part of the recording environment across all phases and all groups. The arena was housed in a light-isolated enclosure (62.3 × 34.2 × 34.2 cm) lined with acoustic foam to reduce external light and sound, with cabling routed through a single small pass-through. A connected computer handled firmware upload, live monitoring, and video capture (Figure 1C).

All files required to reproduce the platform are released with this paper: the arena and water-level ruler STL files, the Arduino calibration and experimental sketches, the OBS-based automated recording scripts, the Tracker position-extraction workflow, and the R analysis code. Assembly, operation, position extraction, and troubleshooting are documented in Supplementary Materials 1–4 respectively.

### A single-session protocol for measuring learning and memory

The behavioral protocol lasted 180 min and consisted of three sequential phases: a 24-min baseline period, a 3-min training period, and a 153-min post-training analysis period (Figure 2). The protocol was programmed in the Arduino IDE and executed automatically after initiation, so that visual stimulus presentation, electrical stimulation, and phase transitions were controlled identically across trials and across fish.

**Figure 2.**
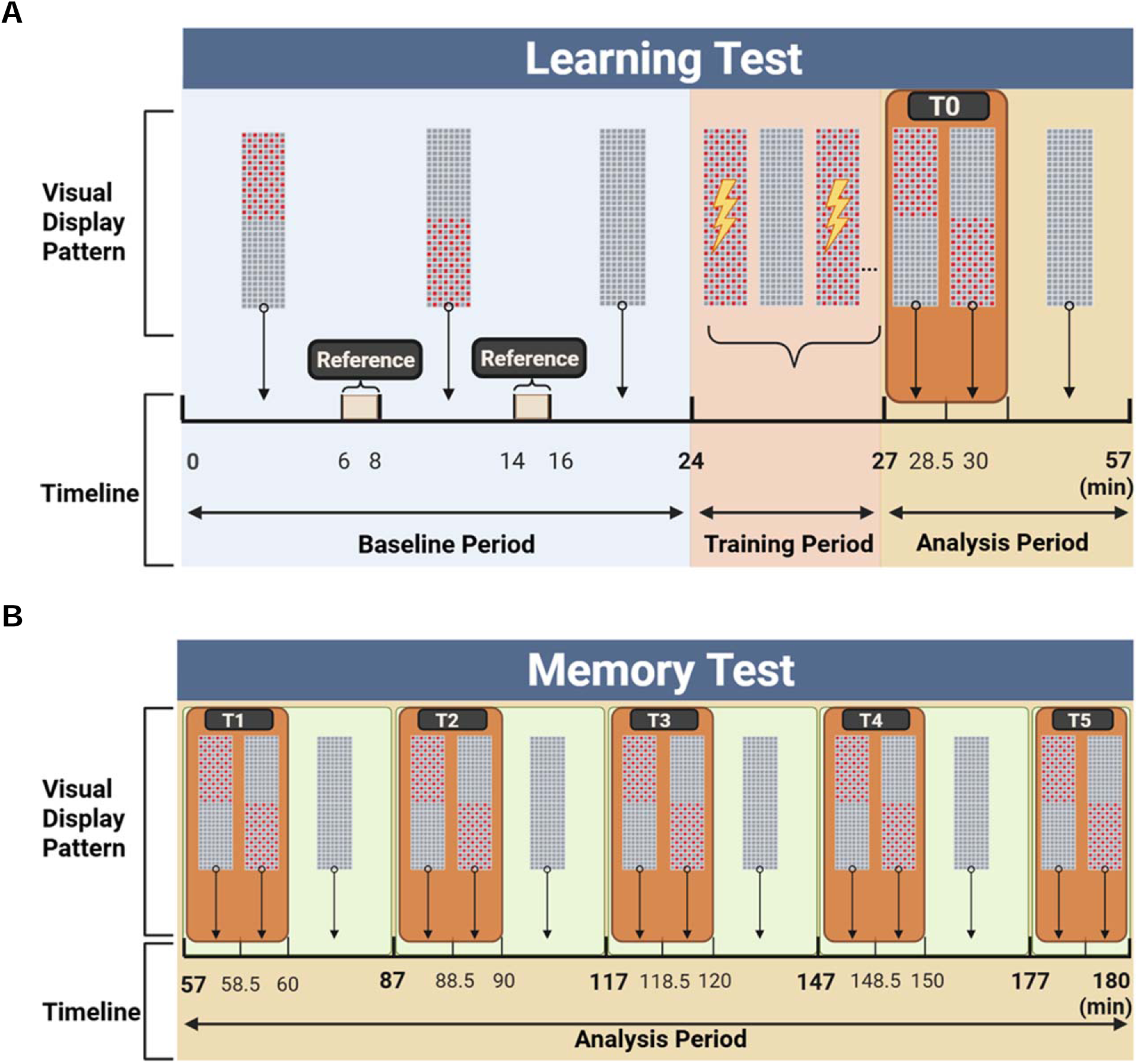
Three-hour zebrafish conditioning and post-training analysis protocol. The behavioral protocol consisted of a 24-min baseline period, a 3-min training period, and a 153-min analysis period. During baseline, one half of the arena displayed the checkerboard pattern for 8 min, followed by 8 min in which the opposite half displayed the pattern, and then 8 min of complete darkness. During training, the full checkerboard pattern was paired with electrical stimulation for 2 s, followed by 13 s of darkness, repeated for 12 cycles. During analysis, six 3-min probe tests were performed at 30-min intervals from T0 to T5. Each test consisted of 1.5 min of checkerboard illumination on one half of the arena followed by 1.5 min of checkerboard illumination on the opposite half, with no electrical stimulation delivered. (**A**) Learning test. (**B**) Memory test.

During the baseline period, zebrafish experienced three consecutive 8-min conditions. In the first 8 min, the checkerboard pattern was displayed on one half of the arena while the opposite half remained dark; in the second, the pattern switched to the opposite half; in the third, the entire arena remained dark (Figure 2A, baseline period). Presenting the pattern on both halves in turn allows any intrinsic side bias to be averaged out of the pre-training reference, and the terminal dark interval provides a rest period that reduces carryover of the immediately preceding visual pattern.

The training period lasted 3 min and consisted of 12 repeated cycles. In each cycle the full chamber displayed the checkerboard pattern while 9.5 V electrical stimulation was delivered for 2 s, followed by a 13-s dark period with no stimulation. Each cycle therefore lasted 15 s, for a total training duration of 180 s (Figure 2A, training period). Because the checkerboard filled the entire chamber and shocks were delivered independent of fish position, training paired the visual stimulus with the aversive stimulus without making stimulation contingent on location.

The post-training analysis period assessed whether zebrafish altered their spatial preference after conditioning. Six probe tests were performed at 30-min intervals. The first, delivered immediately after training (T0), tested learning (Figure 2A, analysis period); the remaining five, at 30, 60, 90, 120, and 150 min after training (T1–T5), tested memory (Figure 2B). Each test lasted 3 min: the checkerboard pattern was displayed on one half of the arena for 1.5 min and then switched to the opposite half for 1.5 min, again controlling for side bias within each measurement. No electrical stimulation was delivered at any point during the analysis period, so all probe measurements are CS-alone and unreinforced. Between probes the arena remained dark for 27 min. Running the learning probe and five memory probes within one uninterrupted session means every measurement comes from the same animal, in the same chamber, without rehandling.

### Wild-type but not *shank3* KO zebrafish acquire conditioned avoidance

Immediate behavioral response to conditioning was evaluated by comparing baseline PI with PI at T0. Wild-type zebrafish showed a significant increase in PI from baseline to T0 by one-tailed paired comparison, indicating a shift toward greater occupancy of the dark region immediately after conditioning (Figure 3A). To test whether the assay could resolve a genotype relevant to neurodevelopmental disorder, the same protocol was applied to a *shank3* KO line. *SHANK3* encodes a postsynaptic scaffolding protein whose haploinsufficiency causes Phelan–McDermid syndrome and is a well-established autism risk factor^30,31^, and existing zebrafish *shank3* models show social, locomotor, and sensory phenotypes but have not been assessed for associative learning^32,33^. In contrast to wild-type fish, *shank3* KO zebrafish showed no change under the identical protocol (Figure 3B).

**Figure 3.**
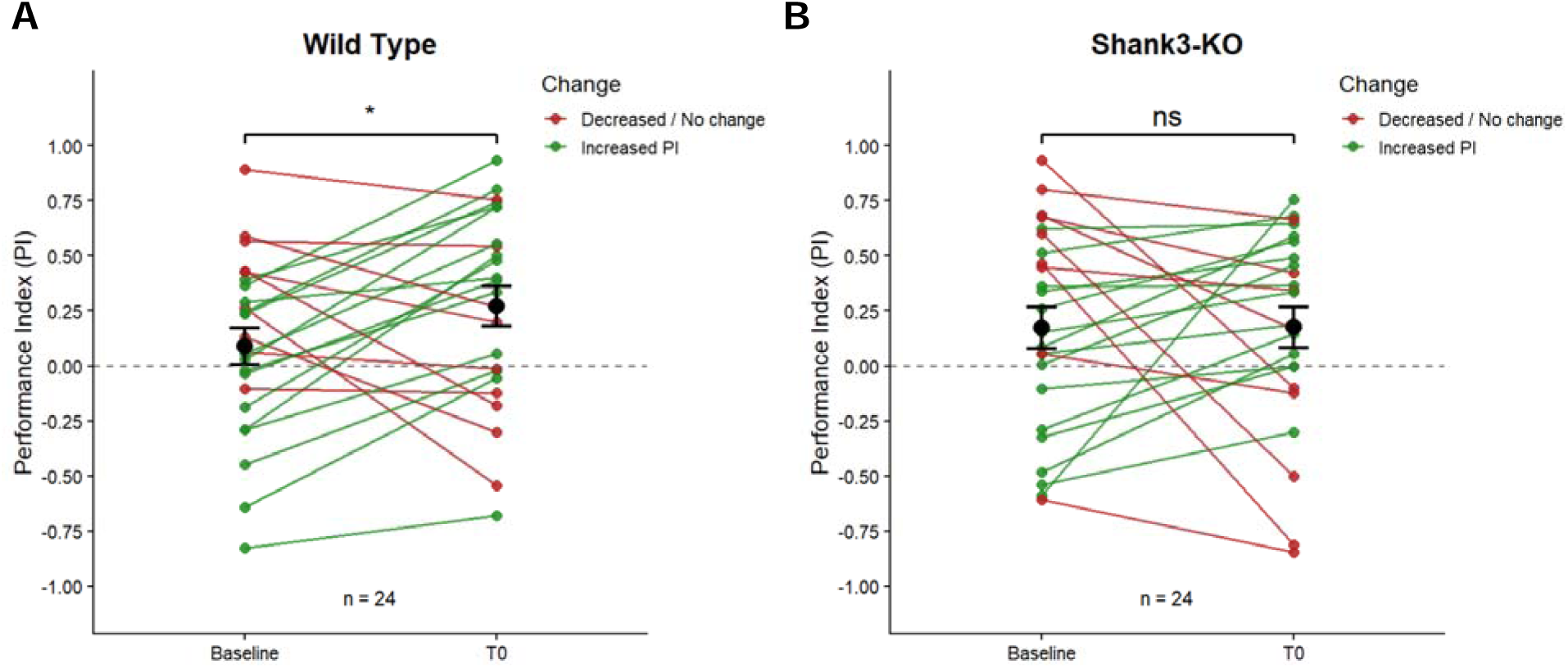
Immediate change in performance index from baseline to T0 in wild-type and *shank3* KO zebrafish. Performance index (PI) was compared between baseline and the first post-training probe, T0. Positive PI values indicate greater occupancy of the dark region, whereas negative values indicate greater occupancy of the checkerboard-illuminated region. Individual paired changes are shown for each fish, with green lines indicating increased PI and red lines indicating decreased or unchanged PI. Black points and error bars represent mean ± SEM. Wild-type zebrafish showed a significant increase in PI at T0 (*n* = 24; *p* = 0.026). *shank3* KO zebrafish showed no significant change (*n* = 24; *p* = 0.490). Significance was calculated by one-tailed paired comparisons. ns, not significant; \**p* < 0.05.

To illustrate how position data give rise to the PI measurement, representative trajectories from wild-type and *shank3* KO fish are shown in Figure 4. In the two baseline reference windows, fish of both genotypes showed mixed occupancy or a weak spatial bias across the chamber. During T0, the wild-type trajectories shifted more distinctly away from the checkerboard-associated region and toward the dark region, consistent with the increase in PI (Figures 4A, 4B; Supplementary Videos 1, 2), whereas *shank3* KO fish continued to traverse both halves (Figures 4C, 4D; Supplementary Videos 3, 4). Because T0 followed training immediately and no stimulation was delivered during the probe, this shift reflects behavior in the absence of reinforcement rather than an ongoing reaction to shock.

**Figure 4.**
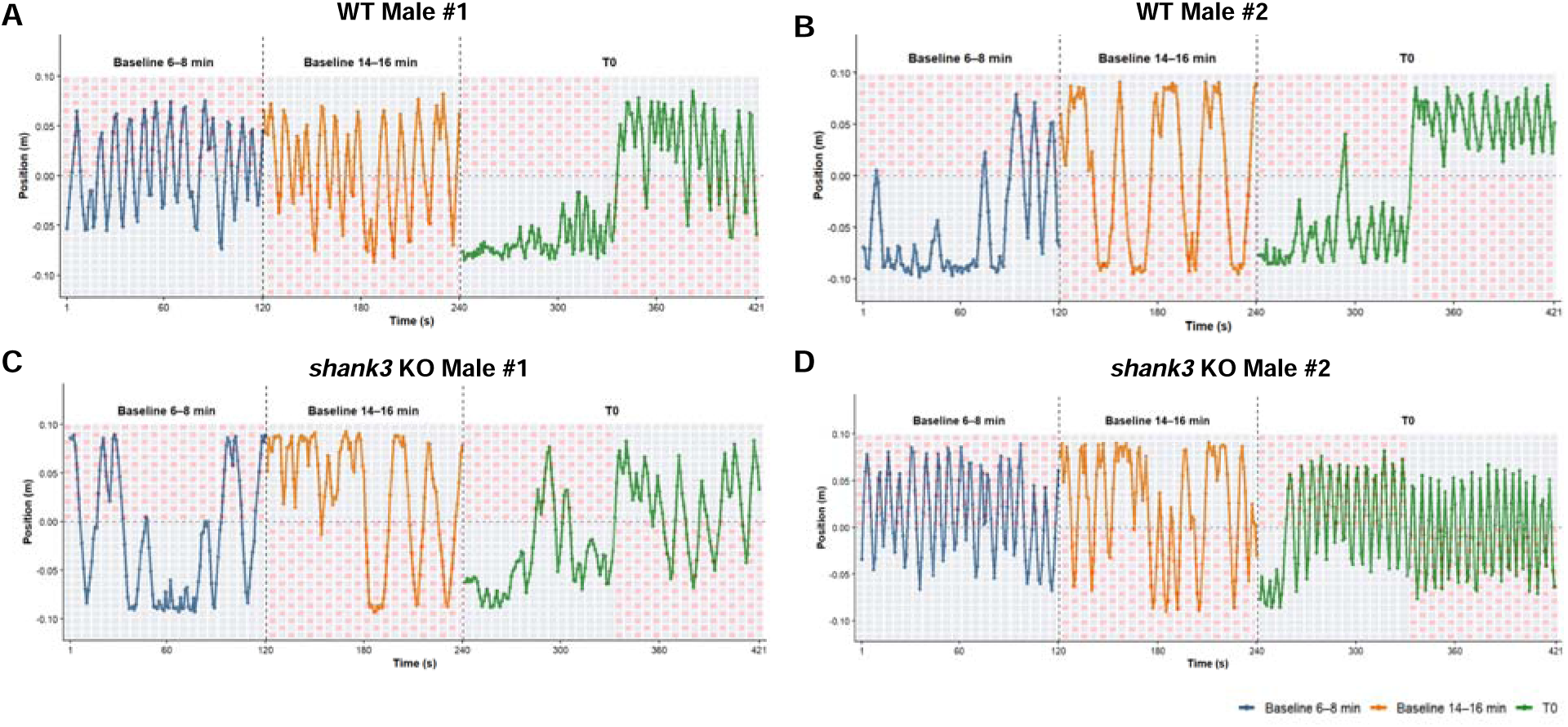
Representative positional trajectories illustrating the PI measurement. Position along the length of the chamber is plotted over time for four representative male fish: wild type (**A**, **B**) and *shank3* KO (**C**, **D**). Each panel concatenates the three windows used for representative trajectory analysis, separated by dashed vertical lines: the baseline 6–8 min window (blue, 0–120 s), the baseline 14–16 min window (orange, 120–240 s), and the first post-training probe, T0 (green, 240–421 s). Points indicate tracked positions extracted from video at one frame per second. Shaded background indicates which half of the chamber displayed the checkerboard pattern during each window. In wild-type fish, occupancy during T0 shifted toward the dark region relative to the two baseline windows, consistent with an increased performance index immediately after training. *shank3* KO fish continued to traverse both halves during T0, showing no comparable shift.

These results indicate that the zAcademy protocol produced an immediate dark-region preference in wild-type zebrafish that was not detectable in the *shank3* KO line under matched conditions.

### The learning response in wild-type zebrafish is sex-dependent

To determine whether the immediate post-training response differed by sex, wild-type zebrafish were analyzed separately as male and female subgroups (Figure 5). While male wild-type zebrafish showed no significant change in PI from baseline to T0 (Figure 5A), female wild-type zebrafish showed a significant increase in PI at T0 by two-tailed Student’s *t*-test (Figure 5B).

**Figure 5.**
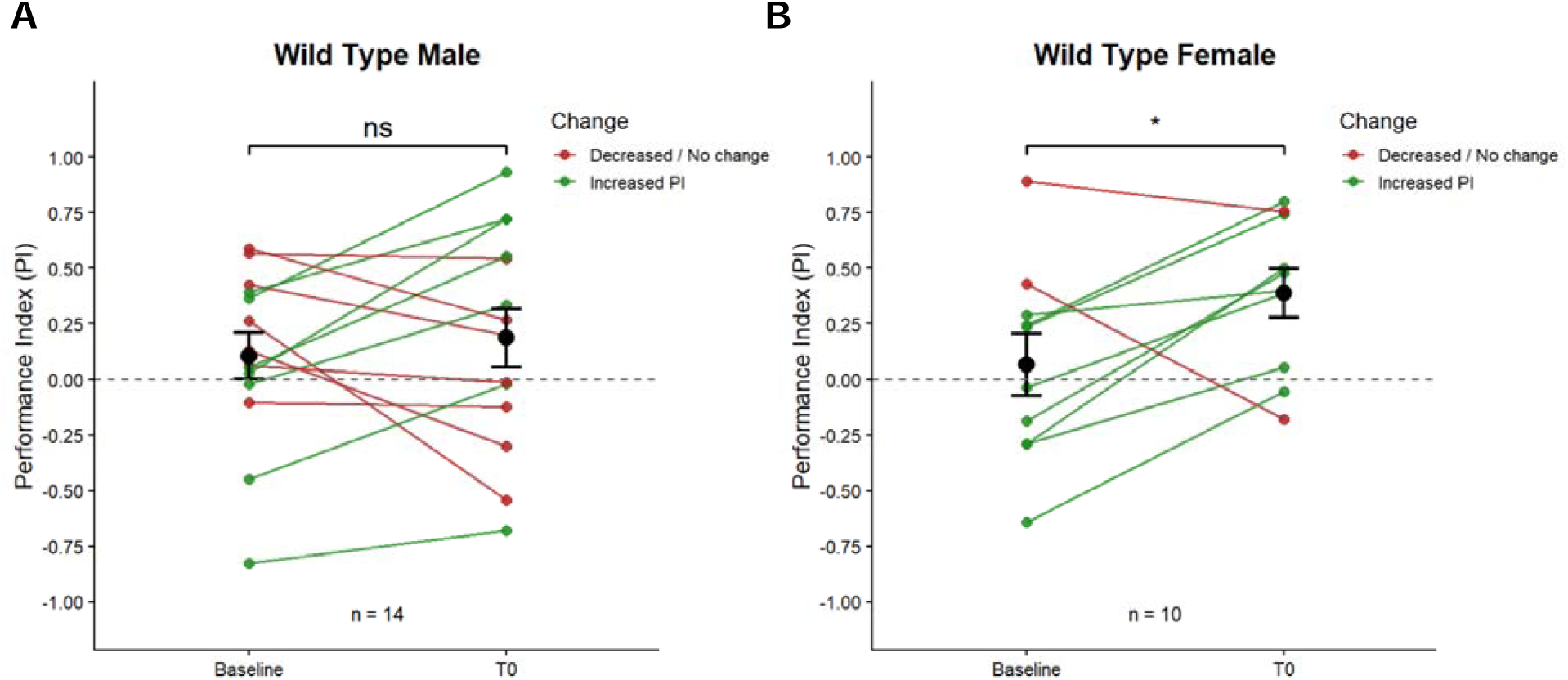
Sex-specific immediate change in performance index in wild-type zebrafish. Performance index was compared between baseline and T0 in male and female wild-type zebrafish. Individual paired changes are shown for each fish, with green lines indicating increased PI and red lines indicating decreased or unchanged PI. Black points and error bars represent mean ± SEM. Male wild-type zebrafish showed no significant change from baseline to T0 (*n* = 14; *p* = 0.498). Female wild-type zebrafish showed a significant increase in PI at T0 (*n* = 10; *p* = 0.0392). Significance was calculated by two-tailed paired comparisons between baseline and T0. ns, not significant; \**p* < 0.05.

The female subgroup therefore accounts for most of the immediate baseline-to-T0 response observed in the pooled wild-type cohort. This pattern indicates that pooling sexes in this paradigm dilutes a response that is present in one sex.

### Memory across the post-training period

To evaluate whether the post-training response persisted beyond T0, PI was analyzed across the full post-training period from T1 to T5 (Figure 6). In the pooled wild-type group, PI was significantly increased at T0 and T5 relative to baseline but not at T1–T4, indicating that a group-level dark-region preference was detectable immediately after training and again at the final probe, with intermediate probes not differing significantly from baseline (Figure 6A).

**Figure 6.**
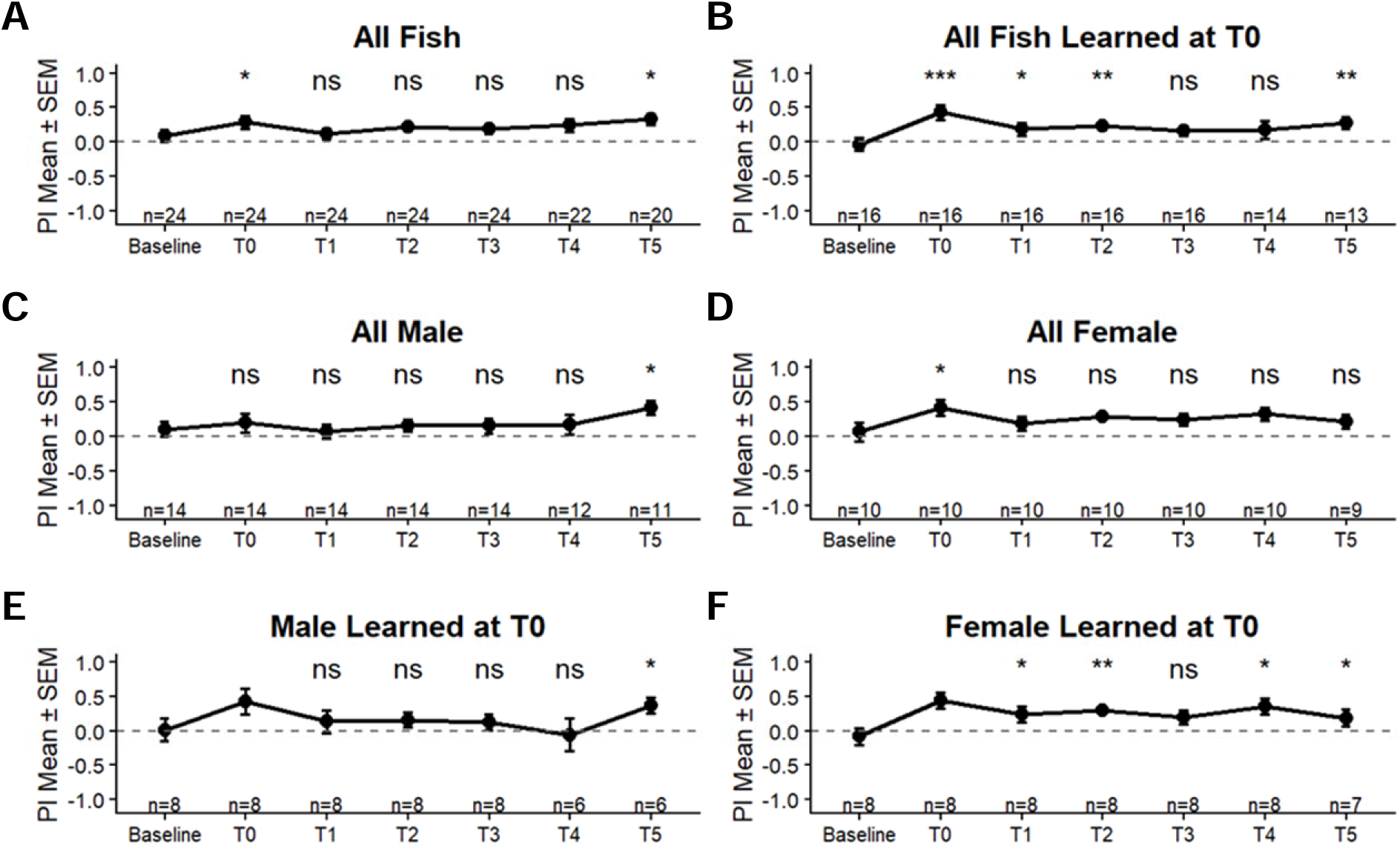
Learning and memory across the post-training period in wild-type zebrafish. (**A**–**F**) Performance index (PI) at baseline and across the six post-training probes, T0–T5. Positive PI values indicate greater occupancy of the dark region, whereas negative values indicate greater occupancy of the checkerboard-illuminated region. Data are mean ± SEM for all wild-type fish (**A**), fish classified as learned at T0 (**B**), all males (**C**), all females (**D**), males learned at T0 (**E**), and females learned at T0 (**F**). Fish were classified as learned at T0 when PI increased from baseline to the first probe. Significance labels indicate paired comparisons against baseline; sample sizes for each time point are shown above the x-axis and decline at later probes because some recordings were excluded when a fish jumped between chambers or when video acquisition was interrupted. ns, not significant; \**p* < 0.05, \*\**p* < 0.01, \*\*\**p* < 0.001.

Because not all fish showed an immediate increase in PI after training, fish were additionally classified as “learned at T0” when their PI increased from baseline to the first probe. This classification is descriptive and based on the direction of change rather than on a per-animal significance test. Because the subgroup is defined by its T0 response, T0 is not an independent test for these fish, and subsequent analyses of this subgroup concern retention (memory) from T1 to T5 rather than acquisition at T0 (learning). In the learned-at-T0 subgroup, PI remained significantly higher than baseline at T1, T2, and T5, while T3 and T4 did not reach significance (Figure 6B).

Sex-resolved analysis showed distinct temporal patterns. Among all males, PI was significantly increased only at T5 (Figure 6C); among all females, only at T0 (Figure 6D). When analysis was restricted to fish classified as learned at T0, males showed a significant increase at T5 (Figure 6E), whereas females showed significant increases at T1, T2, T4, and T5 (Figure 6F). Within this dataset the learned female subgroup therefore displayed the more consistent retention profile across the analysis period (Figures 6E, 6F). Because both learned subgroups were defined by their T0 response and comprised 8 fish each, these comparisons should be read as descriptive of retention among initial responders rather than as independent tests of a sex difference.

Sample sizes decreased at later time points because some recordings were excluded when a fish jumped between chambers or when video acquisition was interrupted by loss of computer power (Figures 6A).

### Individual-level retention

Group means can obscure how many individual animals contribute to an effect at each probe, so retention was additionally visualized at the level of individual fish (Figure 7). Of the 24 wild-type fish tested, 16 showed an increase in PI from baseline to T0 and were classified as learned at T0, while 8 did not show an immediate response. Among the 16 fish that responded at T0, 11 still showed increased PI at T1, 9 at T2, 8 at T3, 7 at T4, and 5 at T5.

**Figure 7.**
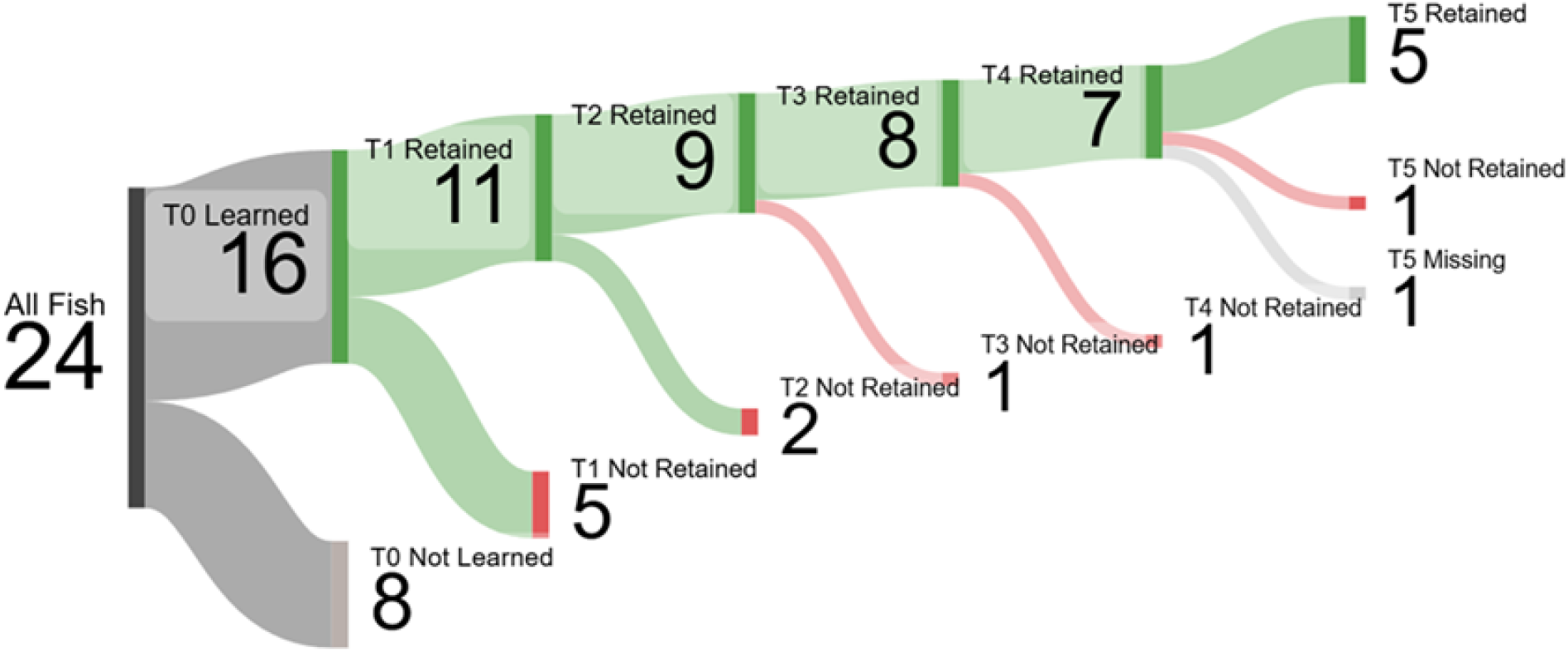
Individual-level retention of the conditioned response in wild-type zebrafish. Flow diagram tracking how the full cohort progressed from T0 learning status through retention outcomes at T1–T5. Of 24 wild-type fish, 16 showed increased PI at T0 and were classified as learned at T0, while 8 did not. Among learned-at-T0 fish, green paths indicate animals that maintained increased PI at each successive probe (11 at T1, 9 at T2, 8 at T3, 7 at T4, and 5 at T5), red paths indicate animals that no longer showed increased PI at that probe, and light grey indicates missing data. Because a fish leaves the retained path at its first failure and is not followed thereafter, these counts are cumulative and are lower than the per-timepoint sample sizes in Figure 6B, which include every learned-at-T0 fish still recorded regardless of its intermediate history.

The number of fish maintaining a dark-region preference therefore declined progressively across the post-training period, with the largest single drop occurring between T0 and T1, although a subset continued to show increased PI at the final probe. Because a fish leaves this path at its first failure and is not followed thereafter, these counts are cumulative and are lower than the per-timepoint sample sizes in Figure 6B, which include every learned-at-T0 fish still recorded regardless of its intermediate history. This flow-based view complements the group means by making explicit that the T5 group-level effect rests on a minority of the original cohort, and that retention was heterogeneous across individuals rather than uniformly present or absent.

## DISCUSSION

We developed zAcademy, an open programmable conditioning arena for adult zebrafish, and used it to measure spatial preference before and after visual–shock pairing within a single three-hour session. Wild-type zebrafish showed an increase in performance index from baseline to the first unreinforced probe, indicating an immediate shift toward the dark region after conditioning, whereas *shank3* KO zebrafish showed no detectable change under the same protocol. Within the wild-type cohort, the immediate response was carried primarily by females, and among fish that responded at T0 the female subgroup sustained elevated PI at more subsequent probes.

The design goal for zAcademy was that another laboratory should be able to rebuild the system from published files and then change the parts of it that matter for their question. Two features follow from that goal. First, stimulus intensity is set by an external bench supply gated by a relay, so shock amplitude is adjusted at the supply rather than by modifying the circuit. This removes the constraint present in fixed-output op-amp designs, where changing intensity requires substituting resistor values^23^, and the relay keeps the microcontroller electrically isolated from the stimulation pathway. Second, the visual stimulus is generated by pixel-addressable LED matrices under firmware control, so pattern geometry, spatial subdivision, and timing are software parameters. Combined with the release of the arena STL files, firmware, recording scripts, and analysis code, this makes the platform straightforward to reproduce and to repurpose for paradigms other than the one reported here. The system also runs two chambers in parallel and is built from inexpensive, widely available components, which lowers the barrier for smaller laboratories and for teaching settings. zAcademy is not the first open zebrafish behavioral apparatus, but open designs remain rare for automated conditioning platforms that combine programmed visual stimulation, programmed electrical stimulation, and long-duration recording.

Prior automated conditioning work in zebrafish has largely been confined to wild-type animals^23,24^, leaving open whether such assays can resolve genotype differences. Existing behavioral characterization of zebrafish *shank3* models has focused on locomotion, social interaction, and repetitive behavior^32,33^, domains in which *shank3b*^−/−^ fish show clear phenotypes. Our data extend this characterization to an associative visual–aversive paradigm: under a protocol sufficient to shift the behavior of wild-type fish, the *shank3* KO line showed no measurable change.

This negative result should be interpreted conservatively. A failure to express conditioned avoidance is consistent with impaired association formation, but it is equally consistent with altered sensory processing of the checkerboard cue, altered nociceptive or stress responses to the stimulation, reduced baseline locomotion restricting the expression of a spatial preference, or a different behavioral strategy under threat. The reduced locomotor activity already reported in *shank3b*^−/−^ zebrafish^33^ makes the locomotor explanation particularly worth excluding. Distinguishing these possibilities requires locomotor-matched analysis, an independent test of visual discrimination for the specific pattern used, and a graded shock–response curve, all of which the programmable stimulation of zAcademy is intended to support. The finding is best read as a demonstration that the platform detects a behavioral difference between genotypes, and as motivation for the mechanistic experiments that would explain it.

Sex is frequently unreported in zebrafish conditioning studies, and where it has been examined the results conflict^26–28^. Our data align in direction with Yin et al.^27^: females but not males shifted spatial preference immediately after training. We additionally observed a sex difference in the shape of the retention profile, with females among initial responders showing elevated PI at multiple early and late probes while males showed a single late increase at T5. Two features of the present design bear on why an effect appeared here. Acquisition and five subsequent retention probes were measured in the same animals within one uninterrupted session, so the retention time course is not reconstructed across separately handled cohorts. The comparison studies did not do this: Fontana et al. assessed retention at a single post-training point^26^, Yin et al. across separate daily sessions^27^, and Corcoran et al. on discrete conditioning days^28^. And because the probes are unreinforced and repeated, a response that is present early and one that emerges late are distinguishable, which a single post-training test cannot do. It is worth noting that female zebrafish show higher baseline anxiety-like behavior^26^, a finding replicated across twenty laboratories in a recent multi-site study of the novel tank test^34^; a sex difference in dark preference under threat is therefore not, on its own, evidence of a sex difference in associative learning.

Several limitations should be considered when interpreting these results. First, the wild-type acquisition effect rests on a one-tailed paired *t*-test (*p* = 0.026) whose two-tailed counterpart does not reach the conventional threshold (*p* = 0.053). The directional hypothesis was specified in advance, but the effect should be regarded as provisional pending replication in a larger cohort. Second, sex subgroup sizes were unbalanced (14 males, 10 females) and were not planned a priori. A formal test of the interaction in a balanced, adequately powered cohort is required before a sex difference can be considered established. Third, PI is a position-based metric and does not measure neural memory formation. Occupancy of the dark region may be influenced by learning, anxiety, locomotor state, sensory sensitivity, or other factors. Finally, sample sizes varied across later time points because some recordings were excluded when fish moved between chambers or when video acquisition was interrupted.

zAcademy detected a conditioning-induced behavioral change in adult wild-type zebrafish, resolved a difference between wild-type and *shank3* KO fish, and detected a sex difference in both acquisition and the shape of the retention profile that warrants further testing in a balanced cohort. The platform is released in full as open hardware and open software, and its stimulation and visual parameters are set in software rather than in hardware. Together these features make zAcademy a practical starting point for laboratories that need a conditioning assay they can both reproduce and modify.

## MATERIALS AND METHODS

### Zebrafish husbandry and experimental animals

Zebrafish were maintained under standard laboratory conditions at 26–27 °C on a 14 h light/10 h dark cycle. Fish were collected from the facility on the morning of each experiment, after routine morning feeding when possible, to reduce variability associated with handling stress and prolonged fasting. During transfer and behavioral testing, animals were monitored for abnormal swimming, immobility, or other signs of stress, and only fish showing normal baseline behavior were included. Two genetic backgrounds were tested: wild-type AB zebrafish and a *shank3* KO line. The *shank3* KO line was generated in-house by the Geng Lab and maintained on the AB background. All experiments were conducted under controlled environmental conditions to minimize external disturbance and improve reproducibility across trials.

### Behavioral apparatus

The dual-chamber behavioral arena was designed in SelfCAD, sliced in Bambu Studio, and fabricated on a Bambu H2D dual-nozzle 3D printer using black PLA, white PLA, and water-soluble Bambu PVA support material. After printing, the arena was submerged in water to dissolve the PVA support. Each chamber measured 192 mm (length) × 62 mm (width) × 120 mm (depth). Upper walls were printed in black PLA to reduce internal reflection; the chamber base was printed in white PLA to transmit light from the LED matrix positioned beneath the arena.

Visual stimuli were delivered by MAX7219-based 32 × 8 dot matrix LED modules, each comprising four integrated 8 × 8 LED panels, mounted beneath the white PLA chamber base. The matrix was addressed as two independently controlled regions (Area 1 and Area 2) corresponding to the two halves of each chamber.

Electrical stimulation was delivered through graphite rod electrodes (6 mm diameter, 90 mm length) inserted along the chamber walls. The two chambers shared a central row of negative electrodes along the divider, while each chamber had a separate row of positive electrodes along its outer wall. Electrodes were connected via alligator clips to the output terminals of a single-channel relay module, which switched an external DC bench power supply.

An Arduino Uno served as the central controller, coordinating LED pattern presentation, relay activation, and protocol timing. A TM1637 four-digit LED display mounted outside the enclosure indicated protocol time. Arduino pin assignments are given in Supplementary Material 1, Table.

Behavior was recorded by a 38 × 38 mm infrared-sensitive camera module with an automatic IR-CUT filter and integrated 850 nm infrared LEDs, mounted above the arena for a top-down view of both chambers. The infrared LEDs engaged automatically during dark periods and disengaged when the LED matrix was illuminated; they were not part of the programmed visual stimulus.

The arena was housed inside a light-isolated enclosure (62.3 cm height × 34.2 cm length × 34.2 cm width) constructed from MakerBeam framing and lined with acoustic treatment foam. Cabling passed through a single small aperture. Complete assembly instructions are provided in Supplementary Material 1.

### Calibration and experimental procedure

Before each session, each chamber was filled with system water to a depth of 3.0 cm, verified using a 3D-printed depth ruler (STL provided). Water depth was held constant across experiments because it affects electrode contact, stimulation consistency, and swimming behavior.

With no fish present, the stimulation circuit was verified using a dedicated Arduino calibration sketch. The external DC supply was set to 9.50 V with a 0.200 A current limit, and the voltage across each positive–negative electrode pair was measured with a digital multimeter in DC mode. Sessions proceeded only when all pairs measured approximately 9.4–9.5 V. An optional reference LED connected in series with a 250 Ω resistor, mounted outside the enclosure, provided a visual indication that the stimulation circuit was active during programmed shock periods; it was not visible to the fish.

The experimental sketch was then uploaded, LED function was confirmed during the startup illumination check, and one fish was introduced into each chamber from individual transfer containers within the 1-min loading window. The enclosure was closed and room lights turned off before recording began. Video recording started at the defined synchronization point at the beginning of the 180-min protocol. Trials in which a fish left its assigned chamber, or in which recording or power was interrupted, were documented and excluded from analysis; affected fish were not re-tested. The full operating procedure and checklist are given in Supplementary Material 2, and troubleshooting guidance in Supplementary Material 4.

### Behavioral protocol

The protocol lasted 180 min: a 24-min baseline period (8 min checkerboard on one half, 8 min on the opposite half, 8 min full darkness), a 3-min training period (12 cycles of 2 s full-chamber checkerboard with 9.5 V stimulation followed by 13 s darkness), and a 153-min analysis period containing six 3-min unreinforced probe tests at 30-min intervals (T0–T5), each consisting of 1.5 min checkerboard on one half followed by 1.5 min on the opposite half, separated by 27-min dark intervals.

### Video acquisition and position extraction

Behavior was recorded either as a single continuous 180-min video, subsequently segmented with LosslessCut, or as automatically captured analysis windows using OBS Studio driven by a Python script via obs-websocket (code provided). Analysis segments corresponded to baseline windows at 6–8 min and 14–16 min and to probe windows at 27–30, 57–60, 87–90, 117–120, 147–150, and 177–180 min.

Position was extracted using Tracker (Open Source Physics). For each clip, a consistent coordinate system was defined with the x-axis along the chamber length and the origin at a fixed chamber corner, and spatial scale was set with the calibration stick tool against a known arena dimension. Videos were recorded at 30 frames per second and Tracker advanced 30 frames per tracking step, yielding one analyzed frame per second (60 analyzed frames per minute). The fish was tracked as a point mass at the center of the body using the autotracker function, with manual correction where tracking failed. Position data were exported as text files using a naming convention encoding strain, trial, time segment, and chamber, and the Tracker project was saved to preserve the coordinate system, calibration, and sampling interval. The full workflow is given in Supplementary Material 3.

### Performance index and statistical analysis

For each analyzed frame, the fish was assigned a positional score of +1 if it occupied the dark region and −1 if it occupied the checkerboard-illuminated region. The performance index (PI) for a given window was the mean positional score across that window: PI = (1/*N*) Σ *s_i_*, where *s_i_* = +1 for the dark region and *s_i_* = −1 for the checkerboard-illuminated region, and *N* is the number of analyzed frames in the window. A PI of +1 indicates exclusive occupancy of the dark region, 0 indicates equal occupancy, and −1 indicates exclusive occupancy of the checkerboard region.

Baseline preference was computed from two reference windows, the final 2 min of each of the two 8-min baseline illumination phases (6–8 min and 14–16 min of the protocol). Combining a window in which the pattern occupied one half with a window in which it occupied the other controls for intrinsic side bias. At 60 analyzed frames per minute, the two windows together contained *N* = 240 frames. Each 3-min probe test contained *N* = 180 frames.

Statistical comparisons used paired *t*-tests comparing baseline PI with post-training PI within each fish. One-tailed tests were used for the directional hypothesis that conditioning would increase PI; two-tailed *p* values are also reported for baseline-to-T0 comparisons. Data are presented as mean ± SEM, and significance was defined as *p* < 0.05. Analyses were performed in R (RStudio).

## ACKNOWLEDGEMENTS

This work was supported by the Sheldon D. Murphy Endowed Chair in Toxicology and Environmental Health and the National Institute of Environmental Health Sciences (NIEHS) of the NIH under the award number R00ES031050. The content is solely the responsibility of the authors and does not necessarily represent the official views of the NIH.

## AUTHOR CONTRIBUTIONS

S.H., B.H., and Y.G. conceived and designed the study. S.H. conducted experiments and analyzed data. S.H. and Y.G. wrote the manuscript.

## DATA AND MATERIALS AVAILABILITY

All data needed to evaluate the conclusions in the paper are present in the paper and/or the Supplementary Materials.

