## Supplementary Material 1 for "zAcademy: an open-source programmable conditioning arena reveals sex- and genotype-dependent avoidance learning in adult zebrafish"

**Arena Assembly and Setup Guide**

**1. Overview**

This supplementary guide describes the construction and operation of the programmable zebrafish conditioning system. The system consists of a 3D-printed dual-chamber behavioral arena, an enclosed testing box, LED matrix modules for visual stimulus delivery, graphite rod electrodes for electrical stimulation, an Arduino-controlled relay circuit, an external DC power supply, a digital timer display, and an overhead infrared camera for behavioral recording. The guide is intended to support replication or modification of the platform by other laboratories.

**2. Required Components and Preparations**

| **Component** | | **Quantity** | **Purpose** |
| --- | --- | --- | --- |
| Behavioral arena | Black PLA (200g) | 1 | Holds zebrafish during conditioning and aligns the LED matrix and electrodes |
|  | White PLA (200g) | 1 |  |
|  | Water-soluble PVA (200g) | 1 |  |
| Enclosure box | MakerBeam starter kit (Including beams, brackets, nuts and bolts) | 1 | Isolates the behavioral setup from external light, sounds, and disturbance |
|  | Acoustic treatment foam (6 packs) | 1 |  |
| Arduino Uno | | 1 | Controls LED pattern timing, relay activation, and protocol progression |
| MAX7219 32 × 8 LED matrix modules | | 6 | Delivers visual stimuli |
| Single-channel relay module | | 1 | Controls electric stimuli |
| External DC power supply | | 1 | Provides electrical stimulation voltage |
| Graphite rod electrodes | | 14 | Deliver electrical stimulation across each chamber |
| Infrared camera module | | 1 | Records zebrafish behavior during the full experiment |
| TM1637 digital timer display | | 1 | Shows protocol time during experiments |
| Acoustic treatment foam | | As needed | Reduced the external noise to the system |
| Alligator clips and jumper wires | | As needed | Used for electrical connections among the components |
| Digital Multimeter | | 1 | Used for voltage calibration |
| Reference LED bulb indicator  (optional) | | 1 | Provides a visible external indicator when the stimulation circuit is active |
| 250 Ω resistor (optional) | | 1 | Limits current and protect the reference LED |

In addition to the components listed above, a computer and a 3D printer are required for system fabrication and operation. A Bambu H2D 3D printer was used to fabricate the dual-chamber behavioral arena from the provided STL file using multi-material printing. A computer was used to run Bambu Studio for slicing, upload the behavioral protocol to the Arduino Uno through the Arduino IDE, and store behavioral videos from the infrared camera.

**3. System Assembly and Setup**

**3.1. Behavioral Arena**

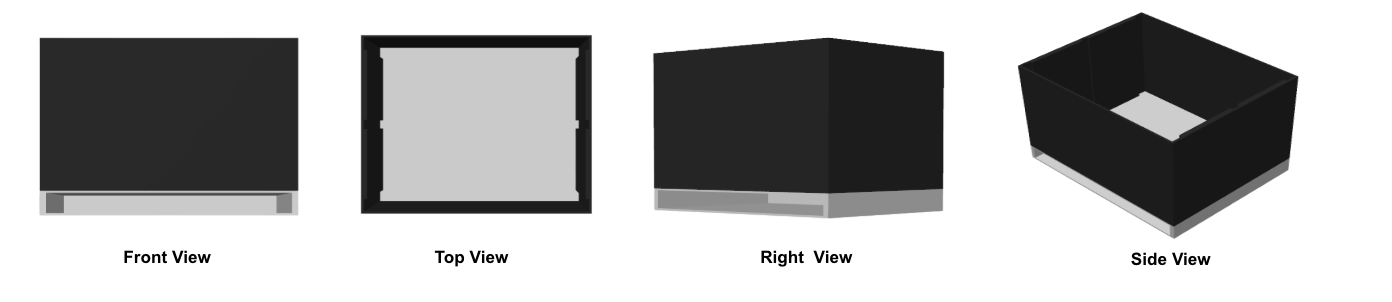

**Figure 1.a.** Behavioral arena design. The arena was fabricated using 3D printing. The 3D-printing model file for the behavioral arena is provided as an *.stl* file in the supplemental materials and should be used as the source file for arena fabrication.

The behavioral arena was designed in SelfCAD and then imported into Bambu Studio for slicing. The arena was fabricated using a Bambu H2D 3D printer, which contains two independent printing nozzles and allows multi-material printing within a single object. Three filaments were used: black PLA, white PLA, and Bambu PVA support material.

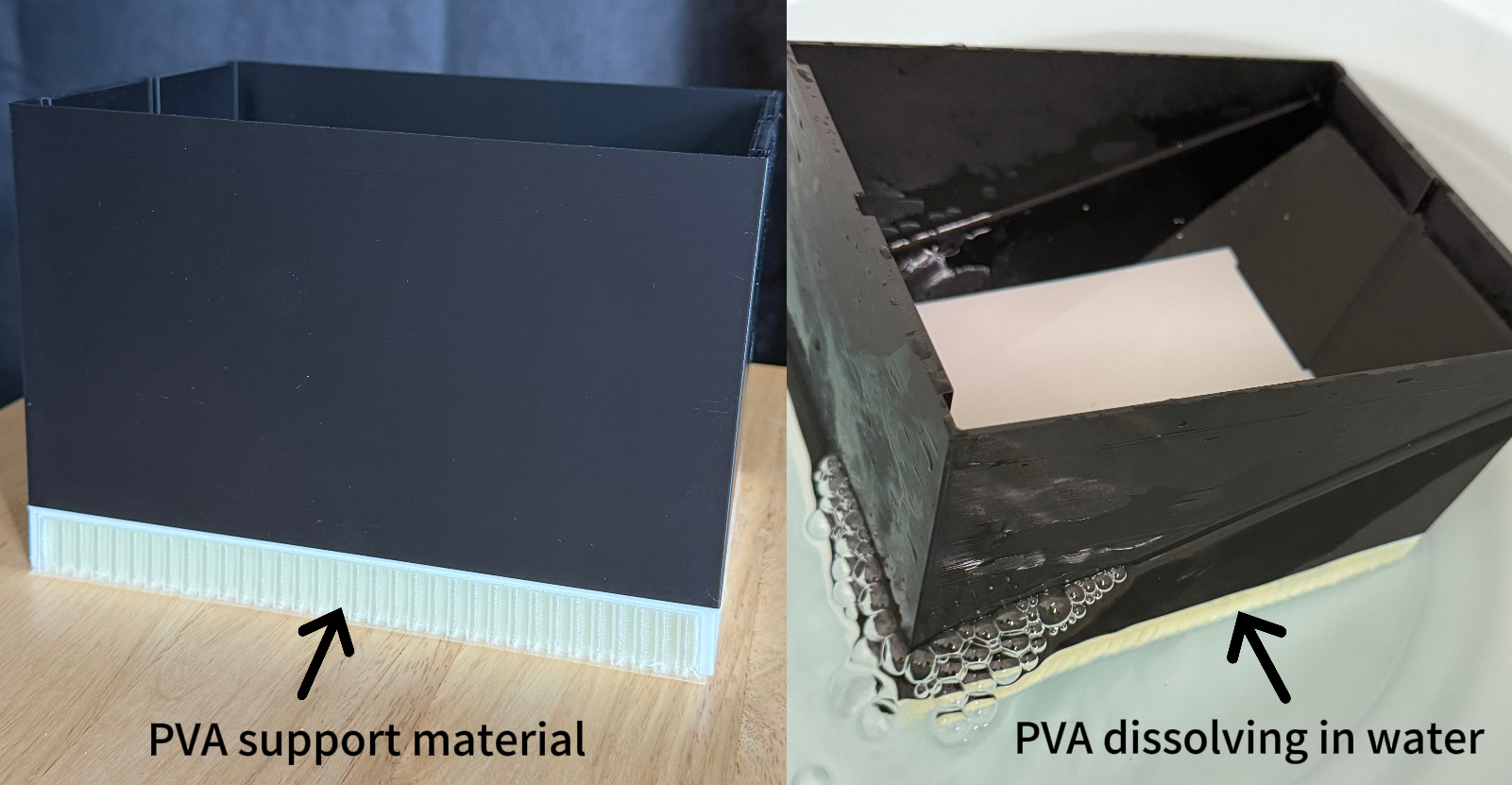

**Figure 1.b.** PVA support material (left) and PVA dissolving in water (right).

PVA was used to support the bottom opening during printing because it can be dissolved in water after fabrication, reducing the risk of damaging the printed structure during support removal. After printing, the arena was submerged in water to dissolve the PVA support material.

**3.2. Electrodes**

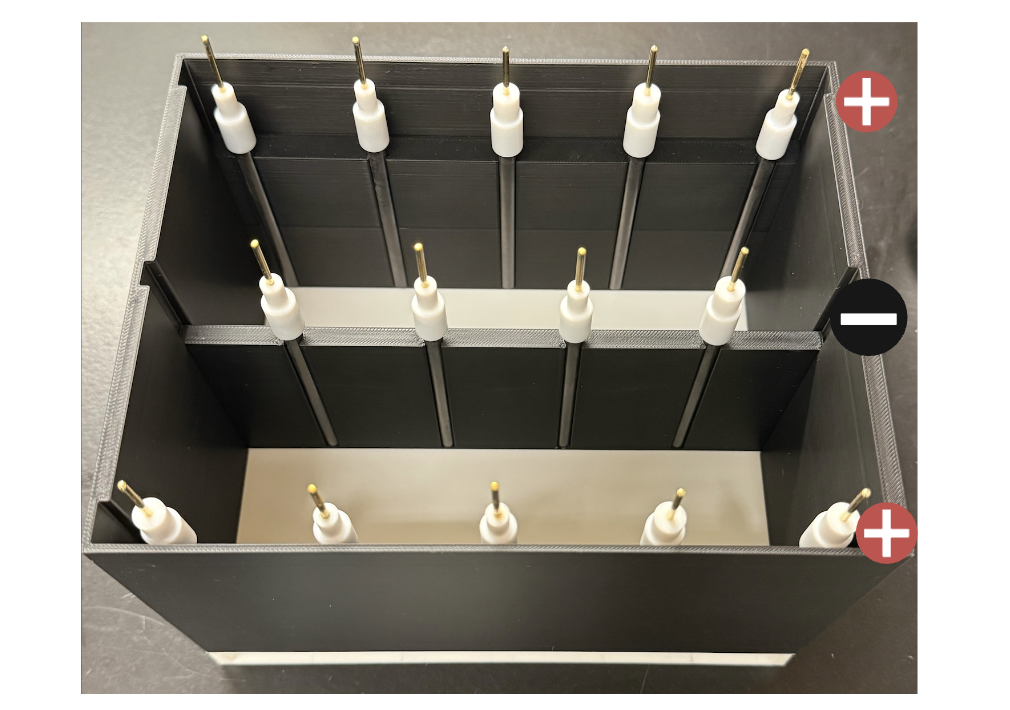

**Figure 2.** Electrode arrangement in the dual-chamber behavioral arena. Graphite rod electrodes were positioned along the side walls of the two-chamber arena. The central row of electrodes served as the shared negative electrode array for both chambers, while the outer rows served as positive electrodes. This configuration allowed electrical stimulation to be delivered across each chamber during conditioning trials.

Electrical stimulation was delivered using graphite rod electrodes integrated into the side walls of the dual chamber arena. Each rod measured 6 mm in diameter and 90 mm in length. As shown in Figure 2, the two chambers shared a central row of negative electrodes positioned along the divider between chambers, while each chamber had a separate row of positive electrodes along its outer wall. This arrangement allowed stimulation to be delivered across the width of each chamber while maintaining a compact electrode layout for the two chamber system.

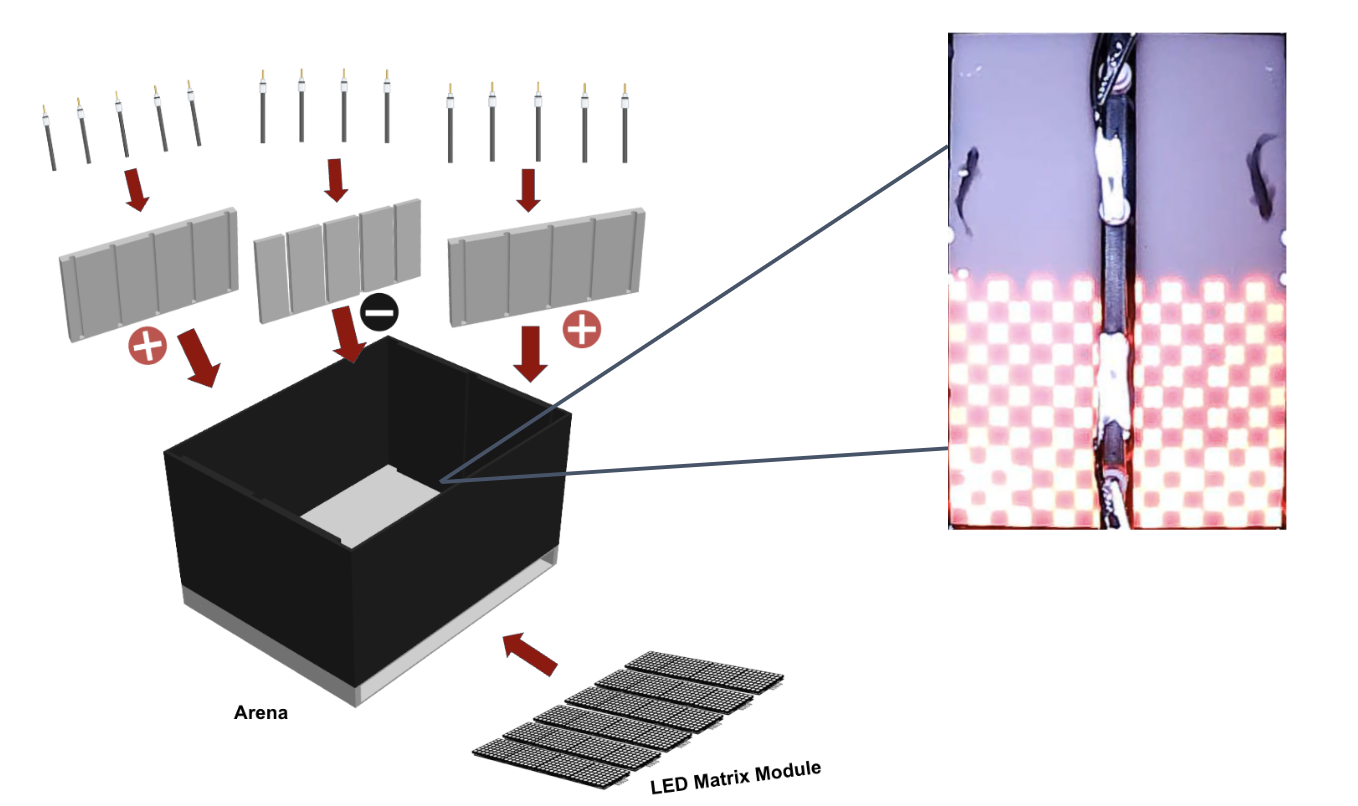

**Figure 3.** Arena assembly and component integration. Graphite rod electrodes were positioned along the chamber walls to deliver electrical stimulation, while the LED matrix module was mounted beneath the arena floor to provide programmable visual cues. The inset image at the right side demonstrates the checkerboard stimulus projected through the translucent base of the arena.

To increase experimental throughput, the arena incorporated two identical behavioral chambers within a single setup. Each chamber measured 192 mm in length, 62 mm in width, and 120 mm in depth. The upper walls of the chambers were printed in black PLA to reduce internal reflection, limit unintended light scattering, and maintain a dark visual environment during conditioning. In contrast, the chamber base was printed in white PLA to permit transmission of light from the LED matrix positioned beneath the arena. This material contrast allowed visual stimuli to be delivered from below while minimizing visual artifacts from the side walls.

A bottom opening was incorporated beneath each chamber to align the LED matrix module with the chamber floor while keeping the electronic display physically separated from the aquatic environment. This configuration allowed controlled visual cues to be presented through the base of the chamber without direct contact between water and electronic components. The arena also included opposing side positions for graphite electrodes, enabling electrical stimulation to be delivered across the chamber during conditioning trials. Together, the dual chamber structure, multi-material design, bottom LED interface, and electrode placement supported reproducible visual stimulation, safe electrical stimulation, and repeated behavioral testing under controlled environmental conditions.

**3.3. Arduino Board**

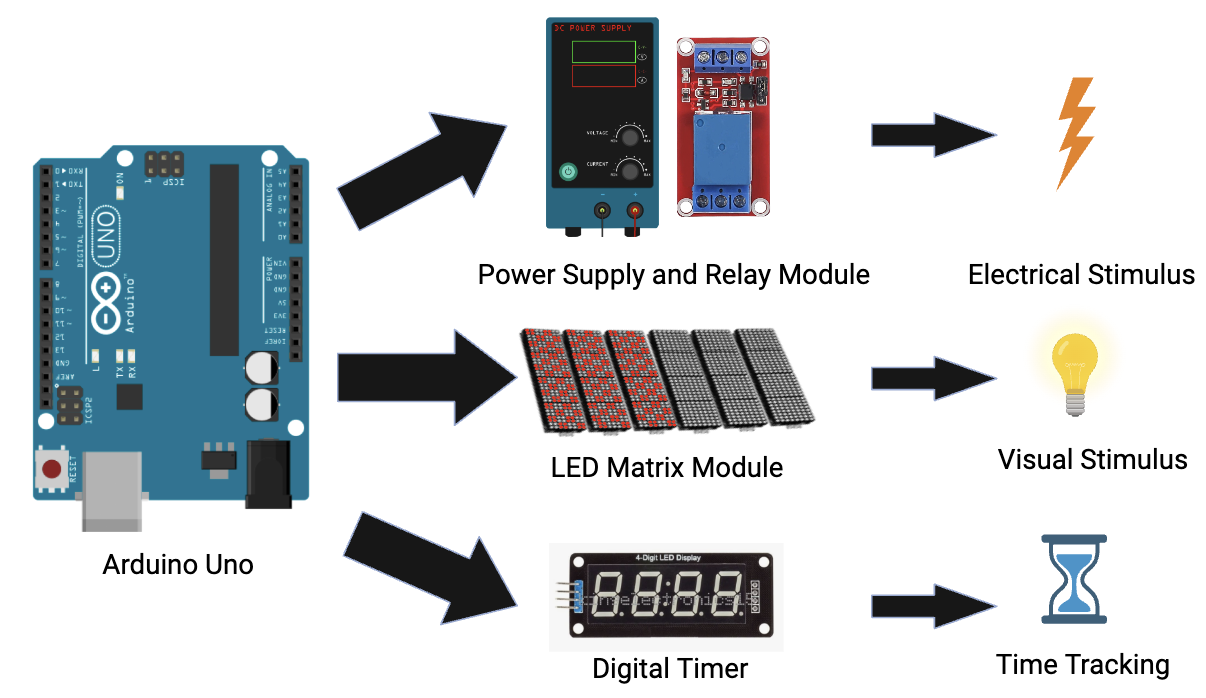

**Figure 4.** Arduino-based control architecture for coordinated stimulus delivery and timing. The Arduino Uno served as the central controller of the behavioral conditioning system, coordinating the relay-controlled electrical stimulation circuit, LED matrix visual stimulus modules, and digital timer display. The Arduino control code is provided in the supplemental materials as an *.ino* file and should be uploaded to the Arduino Uno using the Arduino IDE before running the system.

An Arduino Uno microcontroller served as the central control unit for the behavioral conditioning system. As shown in Figure 4, the Arduino coordinated three major output modules: the relay-controlled power supply for electrical stimulation, the MAX7219-based LED matrix for visual stimulus presentation, and the digital timer for protocol time tracking. Experimental protocols were programmed in the Arduino IDE. During behavioral trials, the Arduino controlled the presentation of checkerboard patterns on the LED matrix, triggered the relay module to deliver electric shocks during the training phase, and updated the digital timer to track protocol progression.

**3.4. Power Supply and Relay Module**

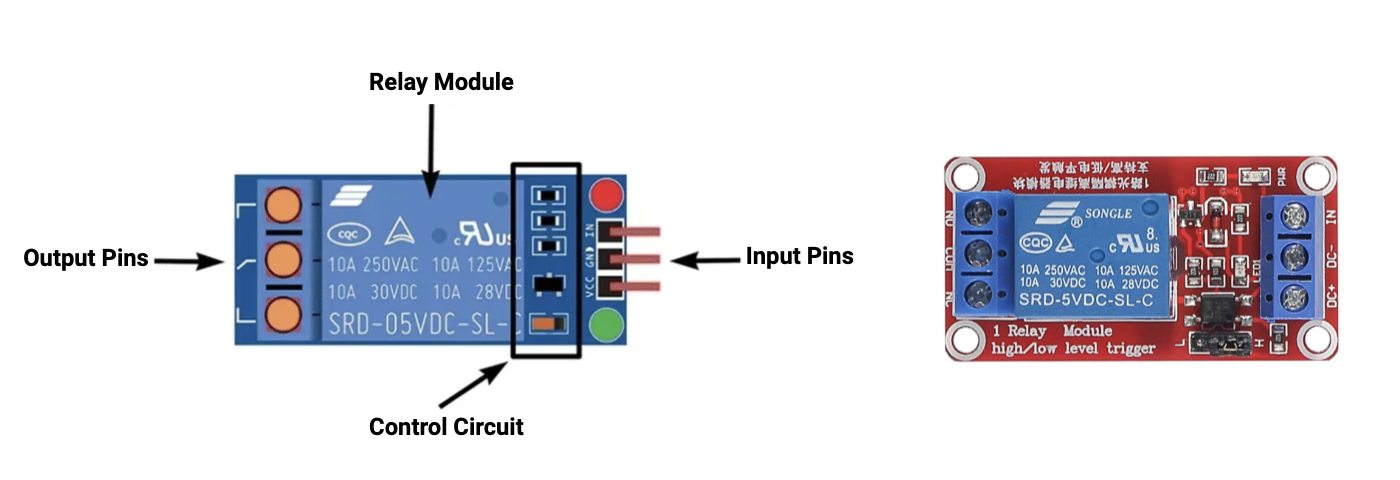

**Figure 5.** Relay module interface for Arduino-controlled electrical stimulation. The relay module separates the low voltage Arduino control signal from the external stimulation circuit. Arduino input pins activate the relay control circuit, while the output terminals switch the external power supply connection to the electrodes during programmed shock periods.

Electrical stimulation was delivered using an external DC bench power supply controlled through a single channel relay module. During the training phase, the Arduino activated the relay according to the programmed protocol. When triggered, the relay closed the external circuit and allowed the power supply to deliver electrical stimulation through graphite electrodes positioned on opposite sides of the behavioral chamber. When the relay was inactive, the circuit remained open and no stimulation was delivered. The external power supply allowed the stimulation voltage to be adjusted during calibration. In the behavioral experiments reported here, the stimulation voltage was set to 9.5 V.

**3.5. LED Matrix Module**

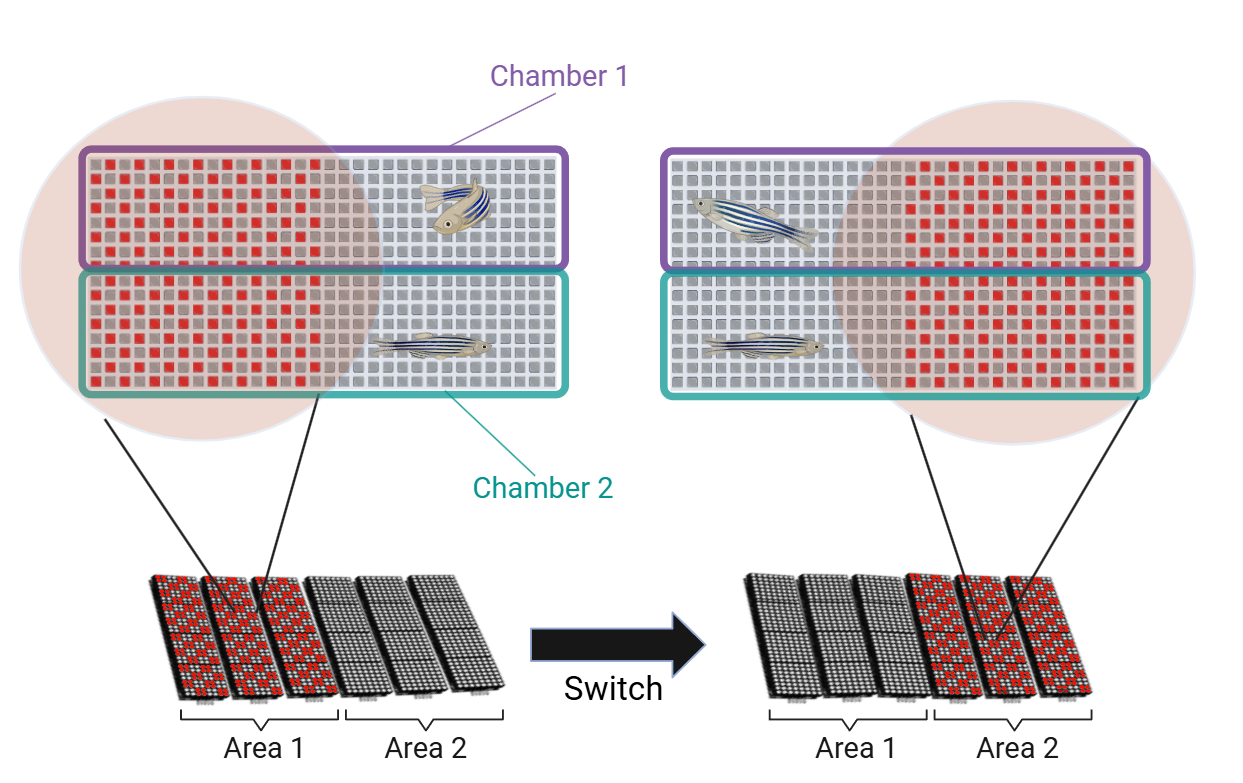

**Figure 6.** Spatial organization and switching logic of the LED visual stimulus. The LED matrix was divided into two independently controlled regions, designated Area 1 and Area 2. During baseline and post-training analysis periods, one region displayed the checkerboard pattern while the opposite region remained dark, after which the illuminated and dark regions switched according to the behavioral protocol.

Visual stimuli were delivered using four MAX7219-based 32 × 8 dot matrix LED modules, each consisting of four integrated 8 × 8 LED panels. The modules were selected for their compact size and ability to generate programmable spatial patterns with pixel-level control. The LED display was programmed to present checkerboard patterns according to the experimental phase. During baseline and post-training analysis periods, the LED matrix was divided into two independently controlled regions, referred to as Area 1 and Area 2 (Figure 6). One area displayed the checkerboard pattern while the other remained dark, and the illuminated and dark regions switched according to the experimental protocol. During the training period, the full chamber displayed the checkerboard pattern while electric shocks were delivered independent of fish position, allowing the visual pattern to be paired with the aversive stimulus.

**3.6. Digital Timer**

A digital countdown timer was integrated into the system using a 4-digit red LED segment display module driven by the TM1637 chip, compatible with the Arduino UNO. This module was used to visually indicate the remaining time in each behavioral training session. The timer was mounted externally on the enclosure, allowing researchers to quickly assess the current stage of the trial and the time remaining without relying on a computer display or interrupting the experimental setup. This addition was particularly useful in environments where multiple trials are being conducted, as it provided immediate contextual information to experimenters without requiring direct interaction with the software interface.

**3.7. Infrared Night Camera**

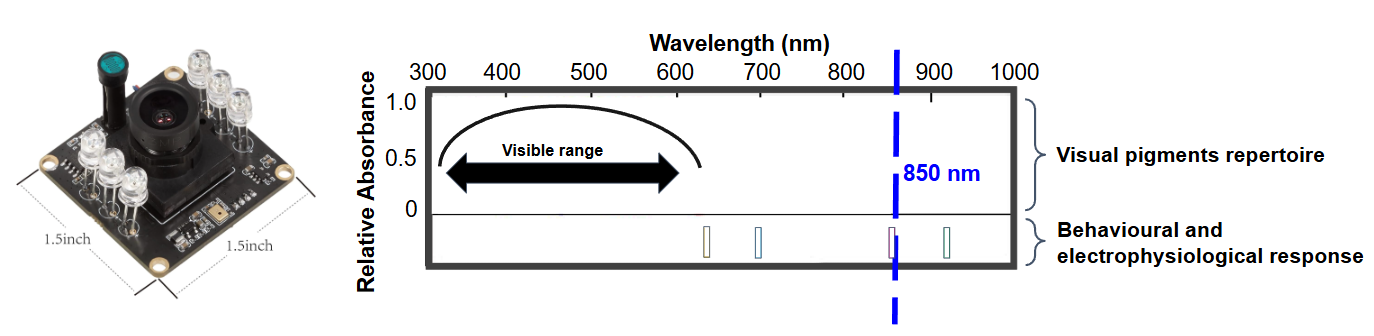

**Figure 7.** Infrared camera module. The infrared camera module contained 850 nm LEDs surrounding the lens and was mounted above the arena for top-down behavioral recording during dark periods.

Behavioral recordings were acquired using a compact infrared-sensitive camera mounted above the dual-chamber arena to capture a top-down view of zebrafish movement throughout the three-hour protocol. The camera measured 38 mm × 38 mm and included an automatic IR-CUT filter with integrated 850 nm infrared LEDs positioned around the camera lens, as shown in Figure 7. These infrared LEDs were separate from the LED matrix used for visual stimulus presentation and were used only to support video acquisition during dark periods.

Under visible light conditions, the camera recorded in standard imaging mode. When the arena became dark, the camera automatically activated its infrared night vision mode, allowing the 850 nm LEDs around the lens to illuminate the chamber for recording. When the LED matrix was illuminated, the infrared mode turned off automatically. Thus, the infrared LEDs were used only to support image acquisition during dark periods and were not part of the programmed visual stimulus delivered from below the arena.

**3.8. Control and Interface Computer**

A desktop computer served as the central platform for interfacing with the behavioral system, providing both power and control to the Arduino. The computer was responsible for uploading experimental code to the Arduino microcontroller, monitoring system behavior, and storing data generated during trials. It also handled the live video feed from the overhead camera, allowing researchers to observe experiments in real time.

**3.9. Arduino Pins Connection and Wiring**

| Component | Pin Name | Arduino Pin | Function |
| --- | --- | --- | --- |
| Breadboard | Positive rail (+) | 5V | Provides 5V power to the components through the breadboard |
|  | Ground rail (-) | GND | Provides the ground reference for the circuit |
| LED Matrix Panel  (Area 1*) | DIN | 11 | Area 1 visual stimulus control |
|  | CS | 10 |  |
|  | CLK | 13 |  |
| LED Matrix Panel  (Area 2*) | DIN | 9 | Area 2 visual stimulus control |
|  | CS | 6 |  |
|  | CLK | 7 |  |
| Relay Module | VoltageCircuit | 2 | Programmed shock delivery |
| Timer Display | CLK | A5 | Protocol time display |
|  | DIO | A4 |  |

**Table 1.** Arduino Pin Connections for the System

*Note.* Area 1 and Area 2 correspond to the two independently controlled LED matrix regions shown in Figure 6.

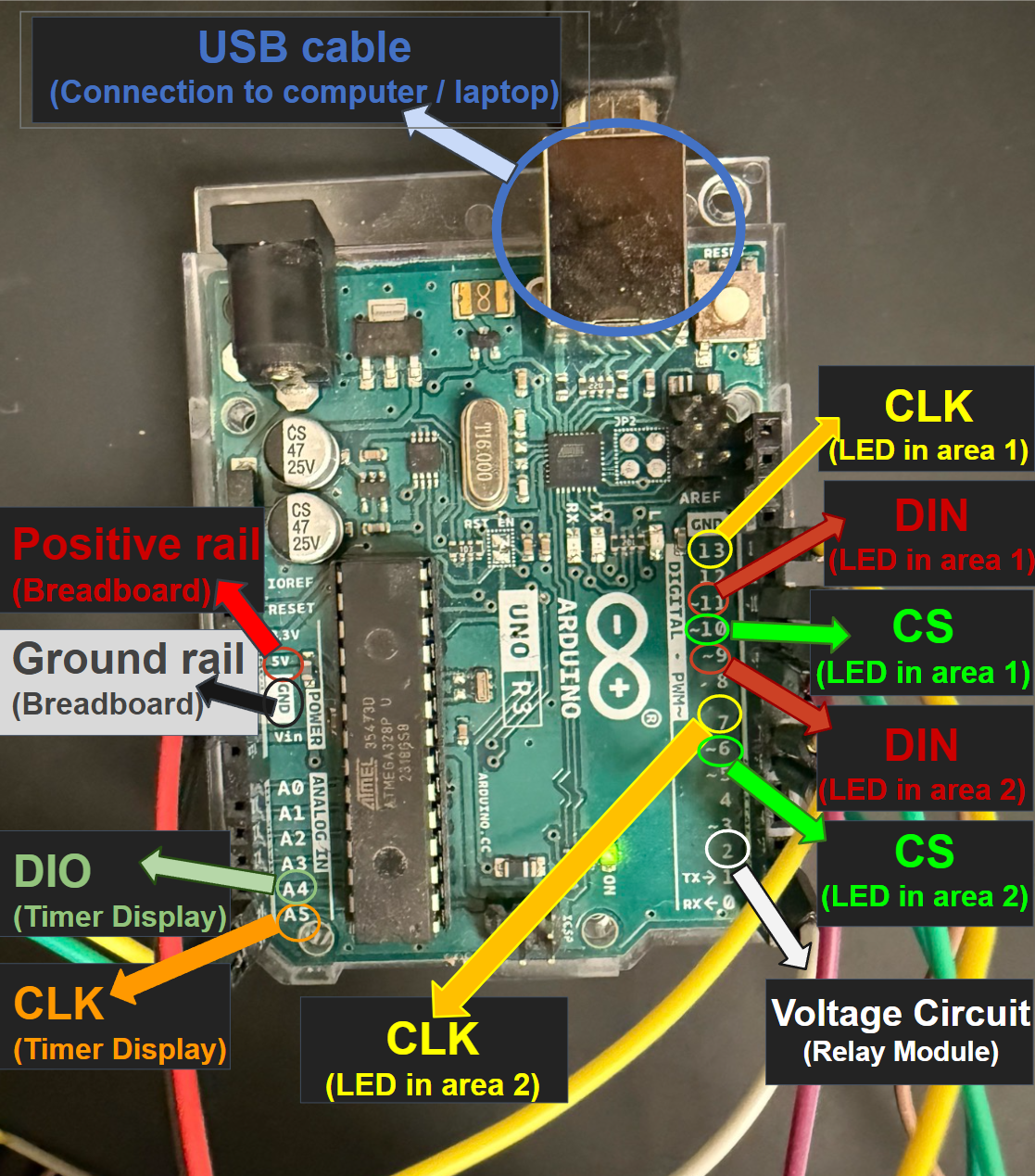

**Figure 8**. Arduino Wiring and Pin Assignment Overview.

| Component | Connection to PC | Power Source | Function |
| --- | --- | --- | --- |
| Arduino Uno | USB cable | PC USB port | Receives the uploaded Arduino IDE program and controls LED stimulus timing, relay activation, and timer display |
| Infrared camera module | USB cable | PC USB port | Provides live top-down video monitoring and records zebrafish behavior during the full experiment |

**Table 2.** PC-Connected Components for System Control and Behavioral Recording

*Note.* The computer should have at least three available connections for system operation: one USB port for the Arduino Uno, one USB port for the infrared camera module, and one connection for the mouse. If the computer has fewer available USB ports, a powered USB hub can be used. Alternatively, a Bluetooth wireless mouse can be used to avoid occupying an additional USB port.

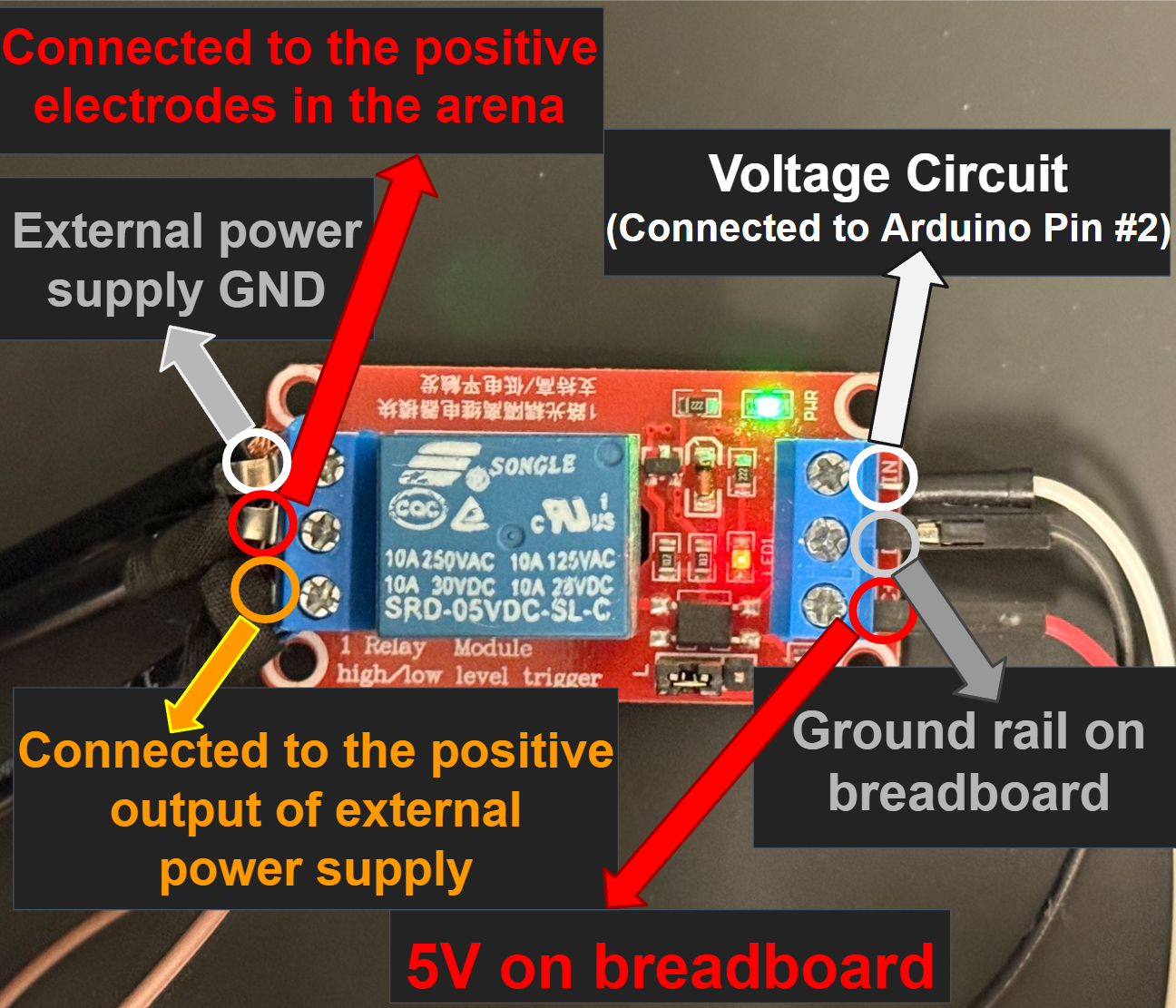

**Figure 9.** Relay Module Wiring Connections.

| Relay Module Side | Terminal / Pin | Connected to | Function |
| --- | --- | --- | --- |
| Input side | DC+ (VCC) | Arduino 5V | Powers the relay control circuit |
| Input side | DC- (GND) | Arduino GND | Provides common ground with Arduino |
| Input side | IN | Arduino digital pin 2 | Connects the power supply to the relay-switched stimulation pathway |
| Output side | NO | External power supply GND | Provides common ground with power supply |
| Output side | COM | Positive electrode pathway | Delivers stimulation voltage to the positive electrodes in the arena |
| Output side | NC | Positive output of external DC power supply | Connects the power supply to the relay-switched stimulation pathway |

**Table3.** Relay Module Connections for Programmed Electrical Stimulation

*Note.* The external DC power supply was set to 9.5 V for the behavioral experiments. The terminal names should be checked against the specific relay module label before wiring, because NO, NC, and COM positions may vary across relay boards.

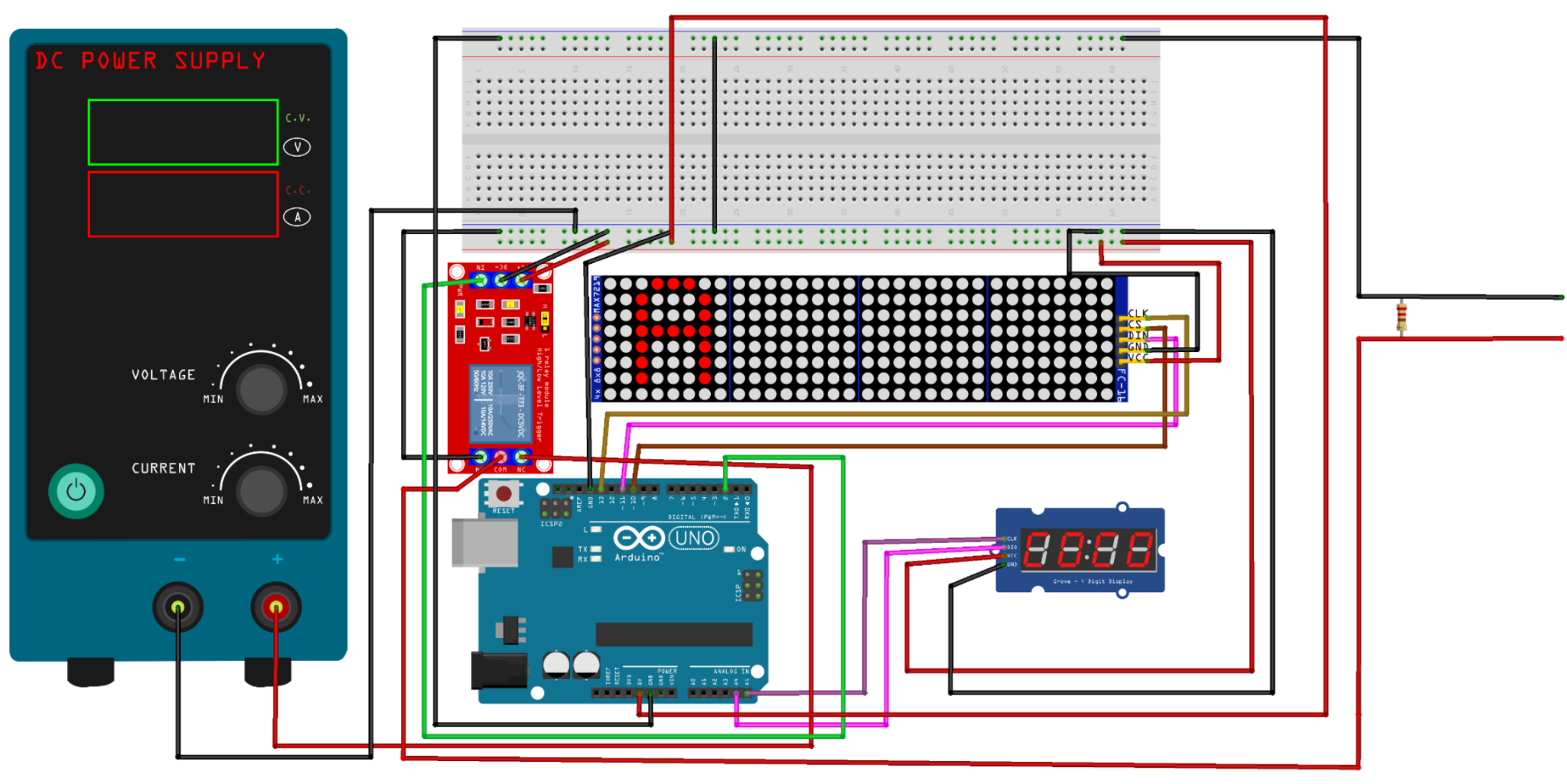

**Figure 10.** Wiring diagram of the zebrafish conditioning system. The diagram shows the electrical connections among the Arduino Uno, LED matrix panels, relay module, timer display, external DC power supply, and electrodes used for programmable visual and electrical stimulation.

*Note.* For clarity, this wiring diagram shows a simplified version of the LED matrix connection. Only one LED matrix module from Area 1 is shown, although Area 1 consists of three LED matrix modules connected in parallel. The LED modules for Area 2 are not shown, but they follow the same connection logic using separate Arduino pins. Area 2 should be connected according to the pin assignments listed in Table 1.

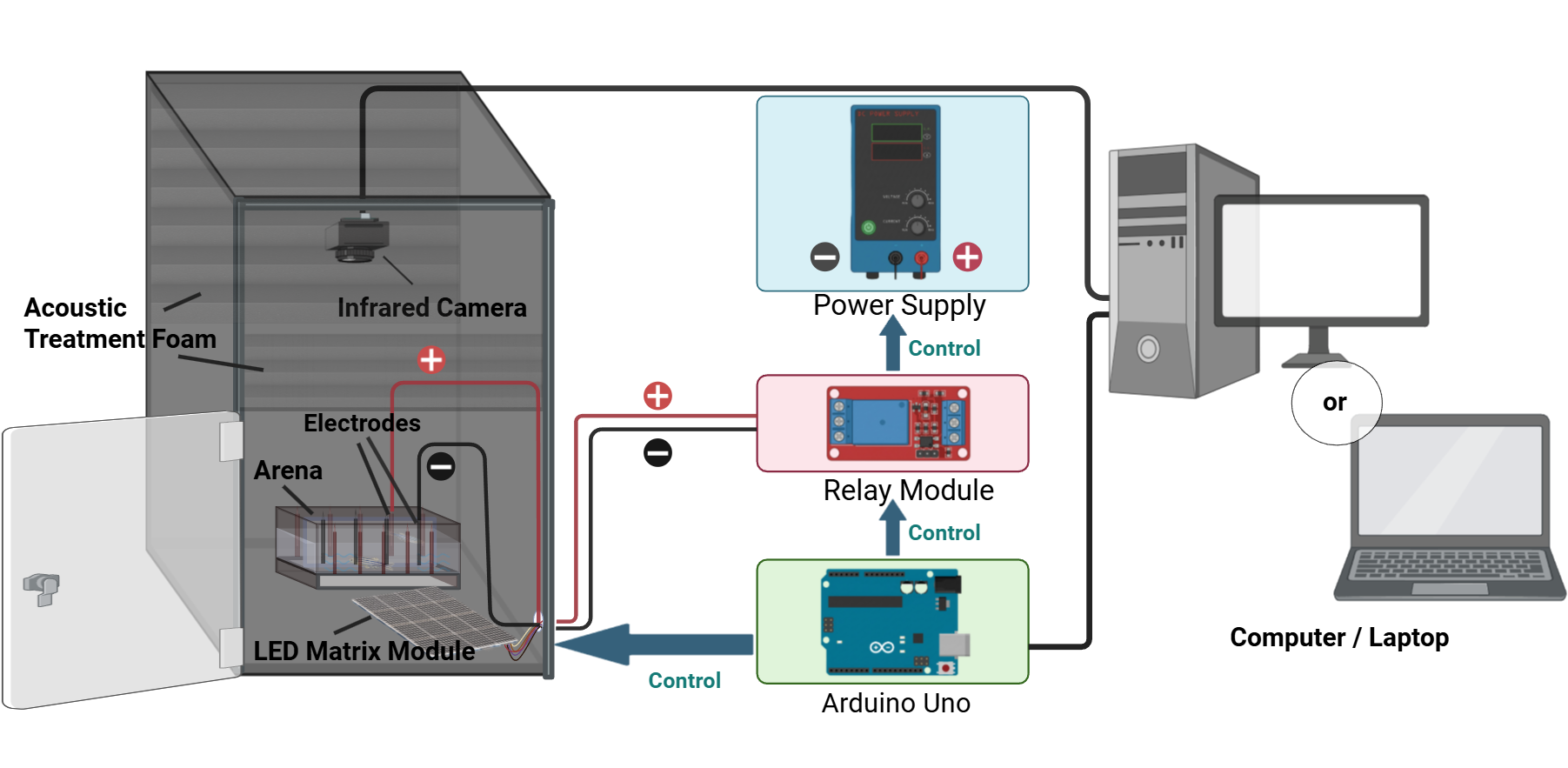

**Figure 11.** Final assembly of the zebrafish conditioning platform. The complete setup integrates a dual-chamber behavioral arena, bottom-mounted LED matrix module, graphite rod electrodes, overhead infrared camera, Arduino Uno, relay module, external power supply, and computer interface.

The final system integrated visual stimulus delivery, electrical stimulation, and behavioral recording within an enclosed experimental setup (Figure 11). The LED matrix was positioned beneath the chamber floor to deliver visual stimuli from below, while graphite rod electrodes were inserted along the chamber walls and connected through the relay-controlled stimulation circuit to the external power supply. An infrared camera was mounted above the arena to record zebrafish movement throughout the full behavioral protocol. The Arduino Uno served as the central controller, coordinating LED pattern presentation, relay activation, and protocol timing, while the connected computer supported code upload, system operation, and video acquisition.

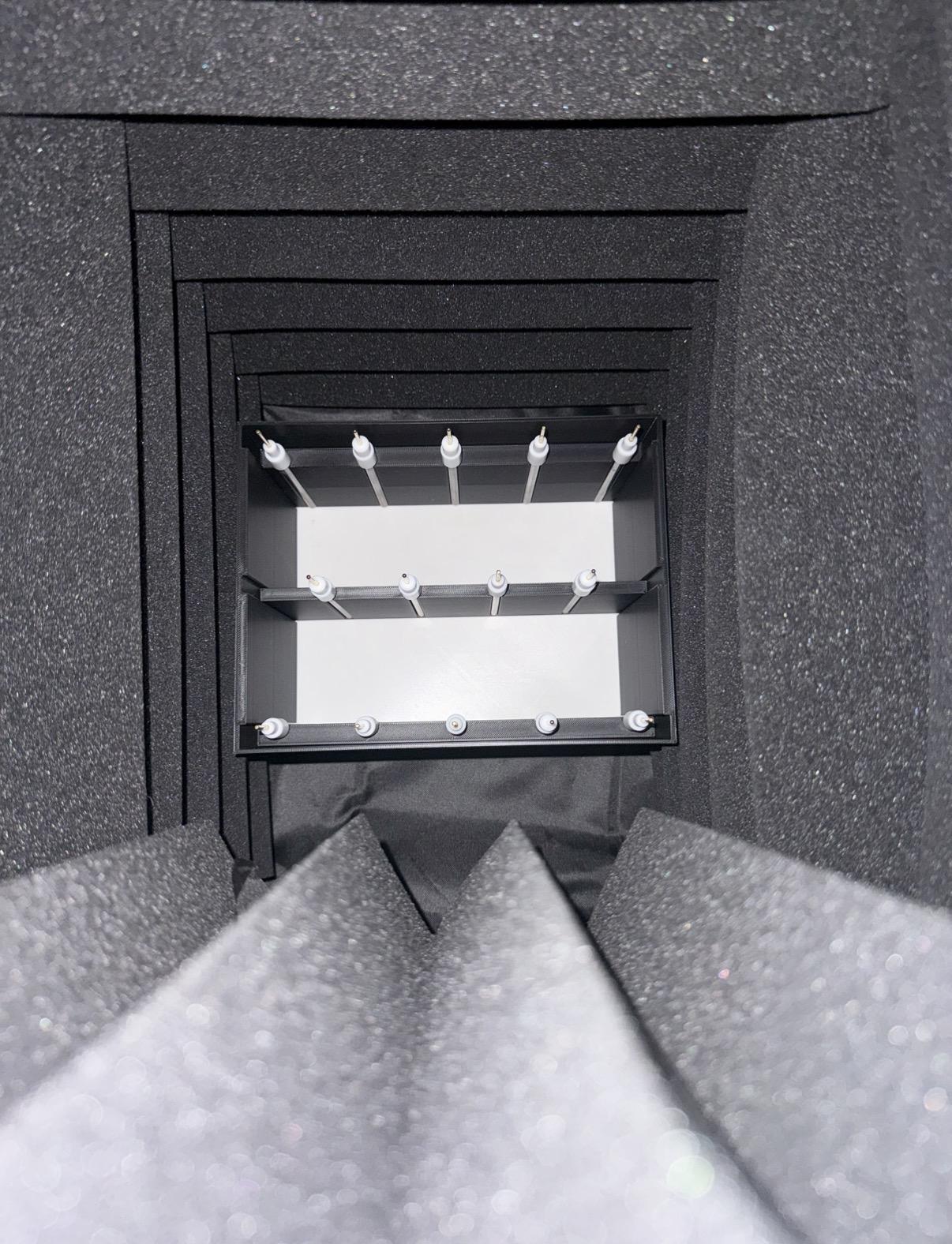

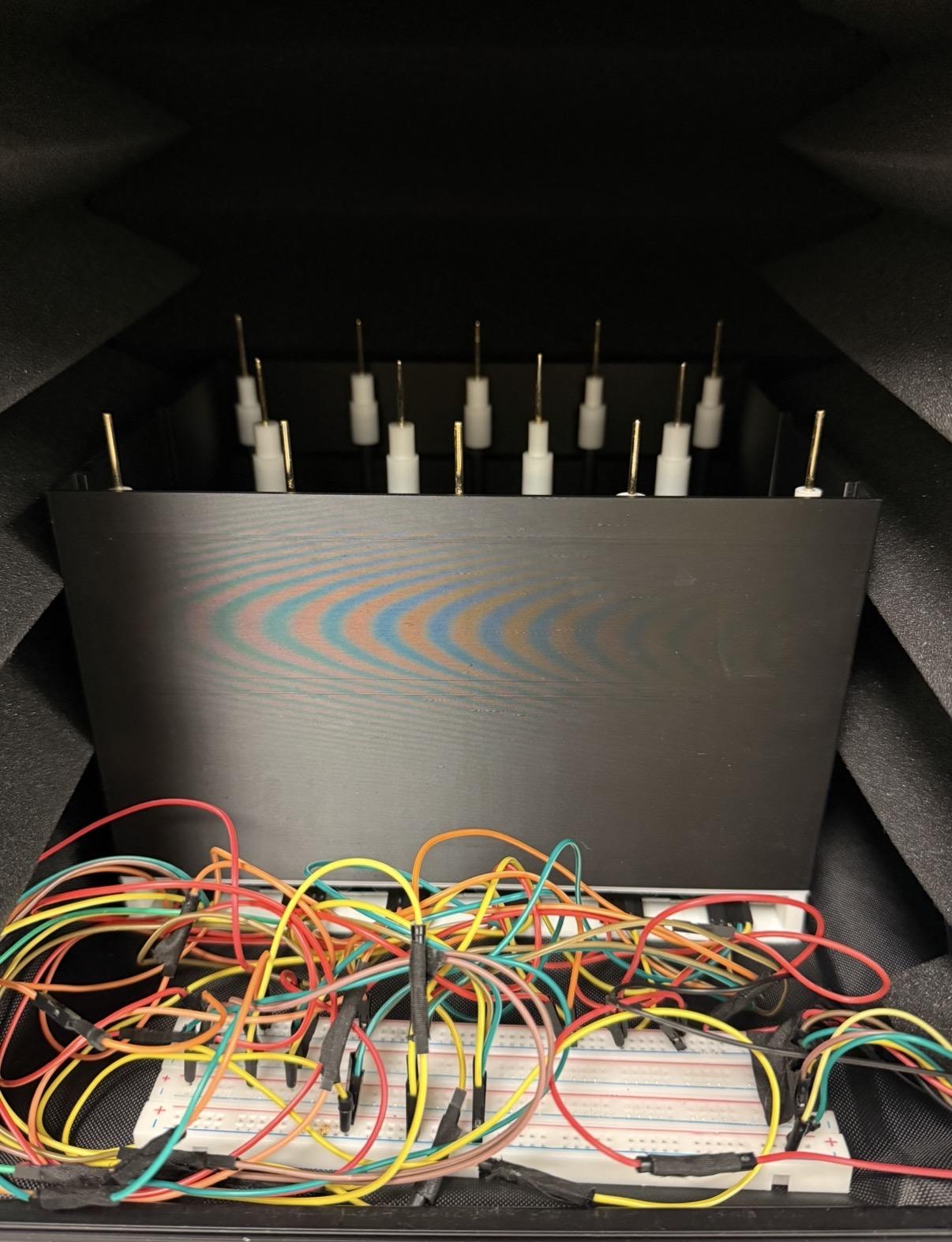

**Figure 12.** Interior view of the enclosure box with acoustic foam lining. Top-down view of the dual-chamber behavioral arena placed inside the enclosure box (left). Acoustic treatment foam was attached to the inner walls to reduce external sound, light leakage, and environmental disturbance during behavioral experiments, while leaving the arena visible for overhead camera recording.

As shown in Figure 12, the dual-chamber arena was placed inside a light isolated enclosure, with acoustic treatment foam attached to the inner walls to reduce external sound and environmental disturbance during behavioral testing.The foam should be positioned so that it does not block the camera view, contact the arena water, or interfere with the wiring.

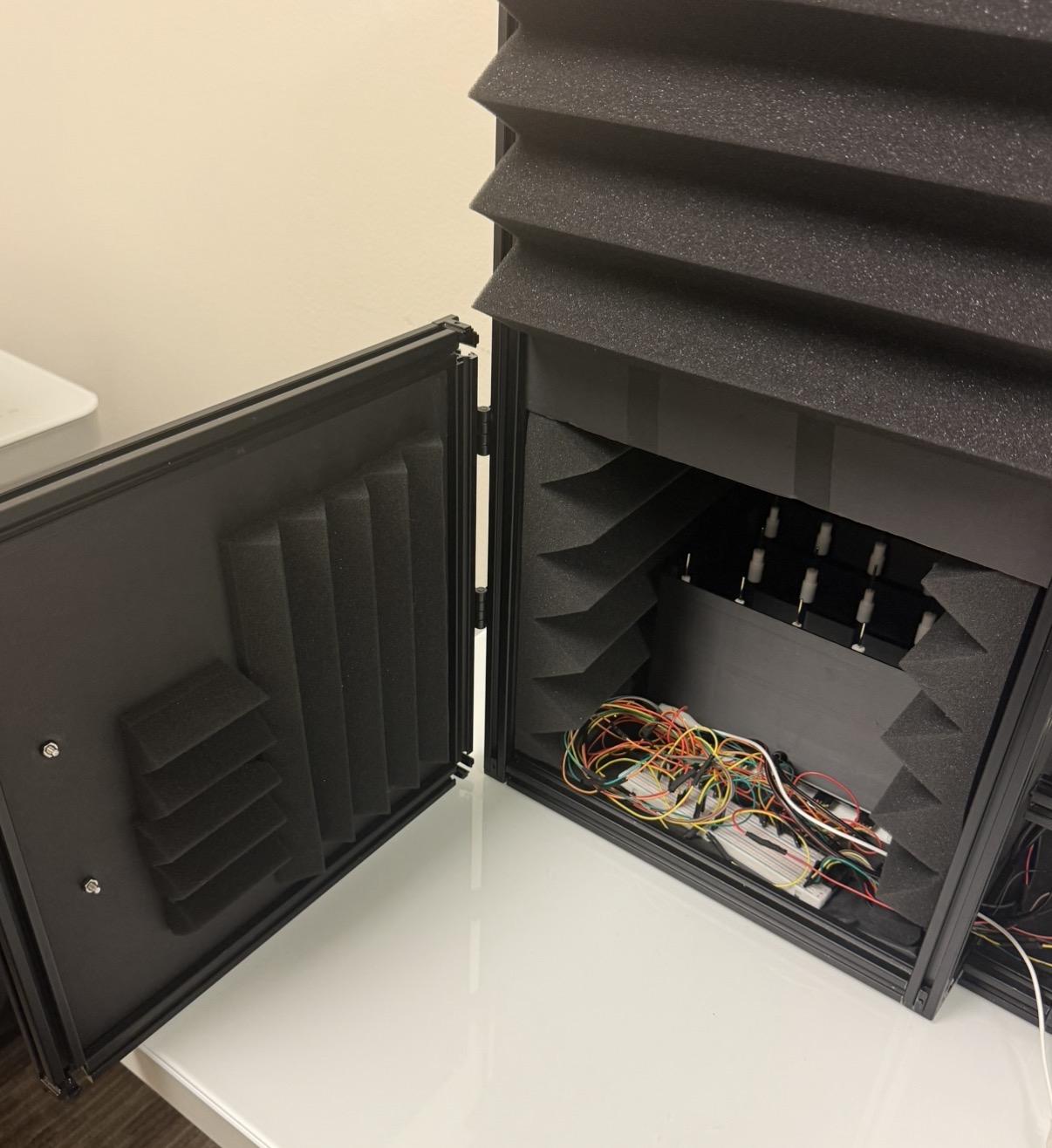

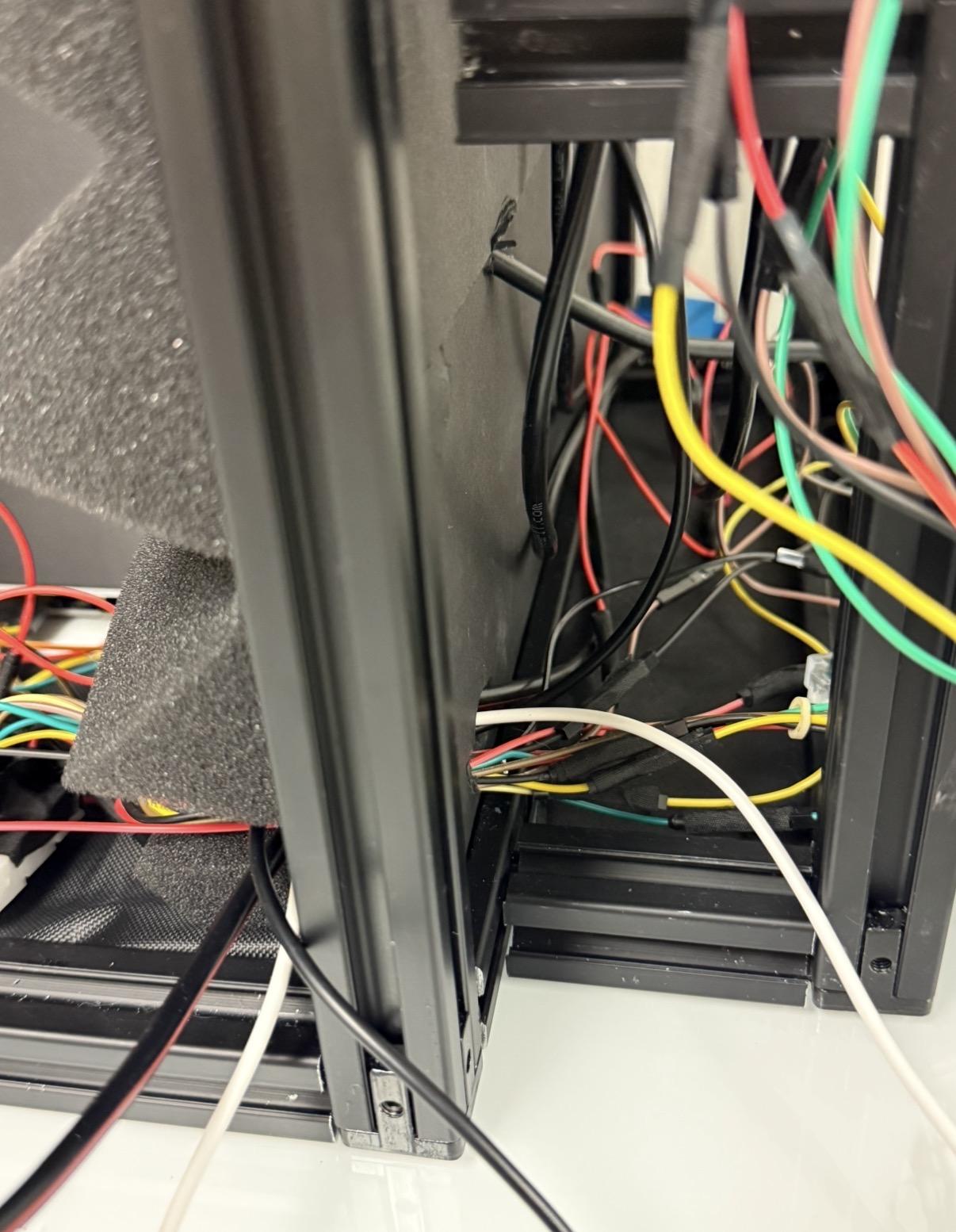

**Figure 13.** Enclosure box assembly and cable routing. Arena positioned inside the enclosure (left). Cable passing through a hole used to route wires (right).

The enclosure used for this system measured 62.3 cm in height, 34.2 cm in length, and 34.2 cm in width. A small hole should be drilled through one wall of the enclosure to allow wires and cables to pass through from the inside of the box to the Arduino Uno and external power supply placed outside the enclosure. The hole should be large enough for the required wires to pass through without pulling or bending sharply, but small enough to minimize external light entering the enclosure.

**Reference LED Bulb (Optional)**

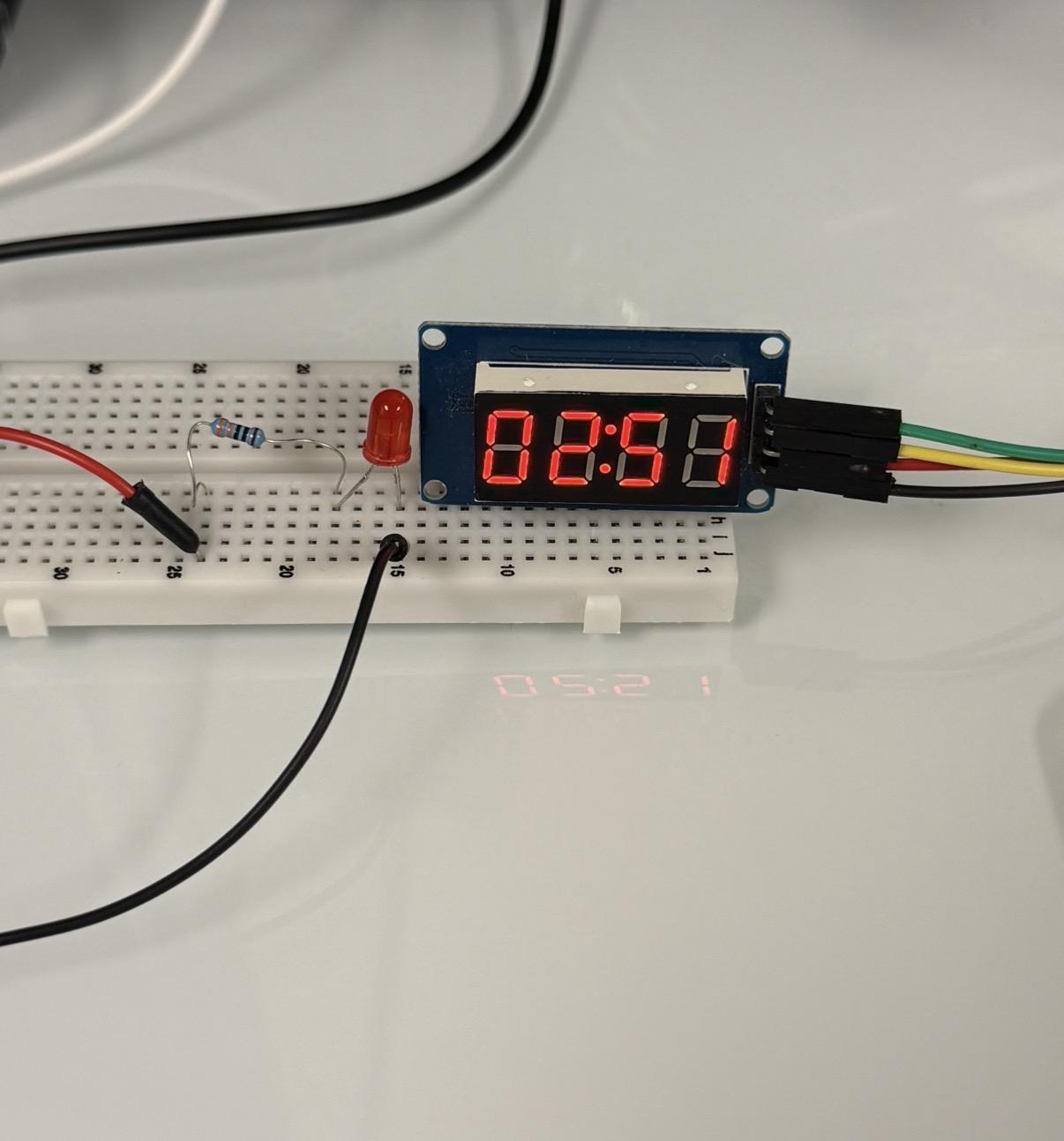

**Figure 14.** Optional Reference LED Indicator for Shock Monitoring. The reference LED bulb was placed on the breadboard outside the enclosure and connected in series with a 250 Ω resistor. When the stimulation circuit was active, the LED illuminated to provide a visible external indicator of shock delivery. The timer display is shown next to the indicator for protocol monitoring.

An optional reference LED indicator was added outside the enclosure to help researchers visually confirm when the stimulation circuit was active. Because electrical stimulation in the water is not directly visible, the LED provides a simple external marker during programmed shock periods. The LED was connected in series with a 250 Ω resistor to limit current and prevent the LED from burning out. This indicator is used only for experimenter monitoring and is not part of the visual stimulus presented to the fish.

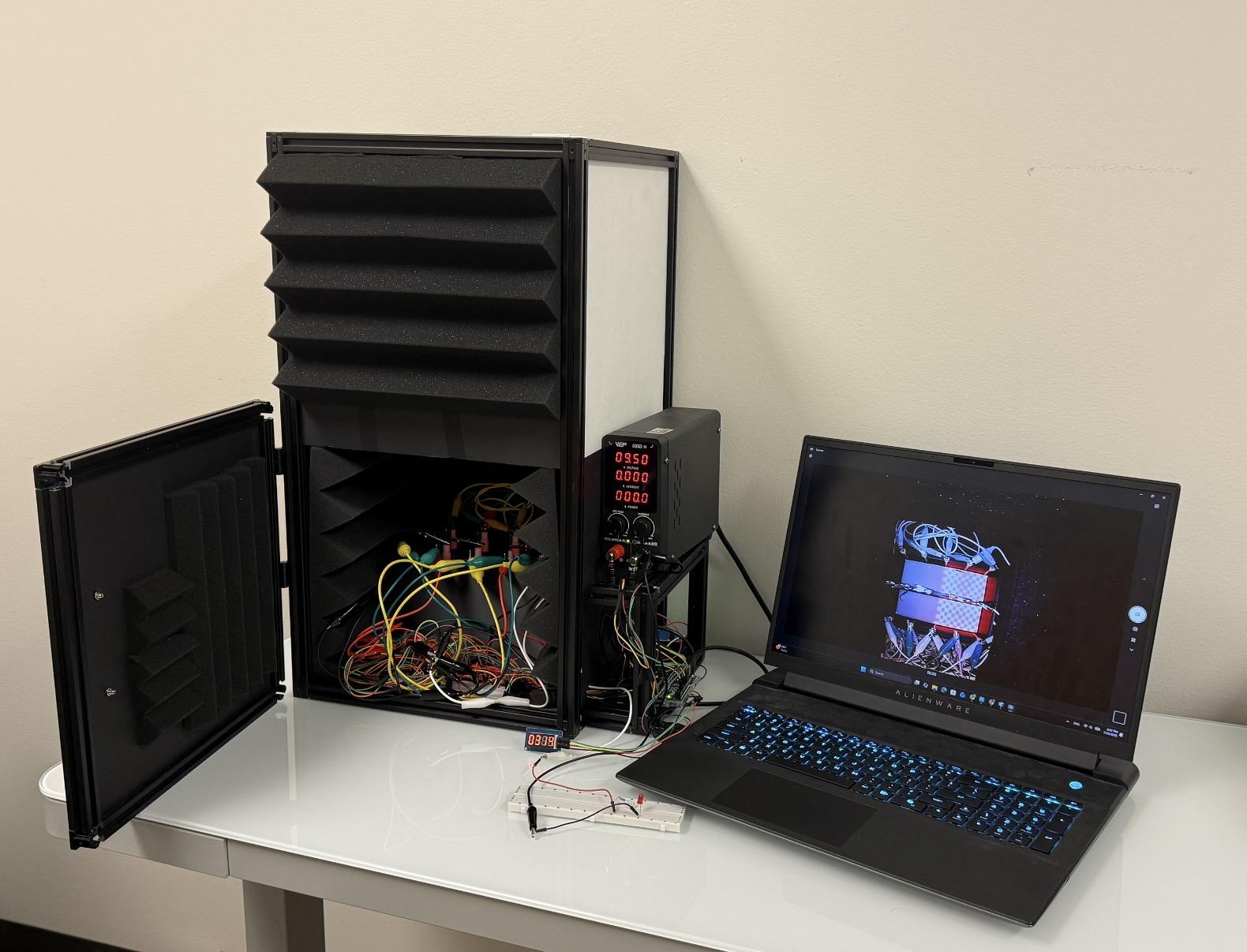

**Figure 15.** Fully assembled zebrafish conditioning system.

After assembly, the full system should resemble the setup shown in Figure 15. The behavioral arena is housed inside the foam-lined enclosure, while the external power supply, Arduino-controlled circuit, timer/reference indicator, and computer remain outside for safe access and monitoring. The computer is used to upload Arduino code and record the top-down camera view during the experiment. Before beginning a trial, researchers should confirm that the wiring is stable, the power supply is set correctly, the camera view is clear, and the enclosure can be fully closed without disturbing the arena or cables.

This supplementary material focuses on the physical setup and assembly of the zebrafish conditioning system, including the enclosure box, 3D-printed arena, electrode arrangement, wiring connections, Arduino-controlled circuit, camera placement, and external monitoring components. It is intended to help researchers reproduce the hardware configuration before running behavioral experiments. Detailed instructions for conducting the actual experiment, including preparation, calibration, fish loading, video recording, and the full step-by-step conditioning protocol, are provided separately in **Supplementary Material 2**: Experimental Protocol.
