## Supplementary Material 2 for "zAcademy: an open-source programmable conditioning arena reveals sex- and genotype-dependent avoidance learning in adult zebrafish"

**Supplemental Material 2.**

**Experimental Protocol**

**1. Overview**

This supplementary protocol provides a practical guide for conducting zebrafish conditioning experiments using the programmable dual-chamber arena. The guide is intended for researchers who are operating the system directly and need a clear workflow for preparing the setup, verifying system function, introducing fish, recording behavior, and completing the experimental run. The experiment is designed to measure zebrafish spatial behavior before training, pair a checkerboard visual stimulus with electrical stimulation during conditioning, and record post-training behavior for later analysis. Video recordings from the experiment are used to extract fish position and calculate behavioral performance metrics.

Because the system combines water, electrical stimulation, programmable LEDs, and live animals, careful preparation and calibration are required before each trial. Researchers should confirm that the arena, electrode connections, LED panels, power supply, Arduino code, and video recording system are functioning properly before fish are introduced. This guide therefore emphasizes both experimental consistency and safe operation of the conditioning platform.

**2. Preparation**

Before running the behavioral experiment, researchers should first complete the system assembly described in Supplementary Material 1: Arena Assembly and Setup Guide which explains how to construct the enclosure, fabricate the 3D-printed arena, install the LED matrix modules and electrodes, and connect the wiring. The present guide, Supplementary Material 2, should be used after the system has been assembled and focuses on the step-by-step experimental workflow for conducting zebrafish conditioning trials.

In addition, researchers should also confirm that the computer or laptop recording software can support continuous 3-hour video recording. Some built-in camera recording applications may have recording-length limits, storage limits, or automatic shutdown settings. If continuous full-length recording is not supported, researchers should use an alternative recording solution. Two possible options are: installing video recording software that supports continuous 3-hour recording, or using software that can automatically capture and save the selected analysis windows during the experiment. The full recording is preferred because it preserves the complete behavioral session. However, if only selected time windows are recorded, researchers must ensure that all baseline and post-training analysis windows are captured correctly. Detailed instructions for video segmentation, position extraction, and data export are provided in ‘Supplementary Material 3: Tracker-Based Zebrafish Position Extraction Workflow’.

Before calibration or animal testing, confirm that:

| **Item** | **Check** |
| --- | --- |
| Arena position | Arena is placed securely inside the enclosure |
| Water level | Both chambers are filled to the same standardized level  (at least 3cm) |
| Electrode placement | Graphite electrodes are inserted and submerged consistently |
| Wiring | Alligator clips and wires are stable and routed away from water-contact areas |
| LED matrix modules | LED panels are aligned beneath the chamber floor |
| Camera | Infrared camera captures a clear top-down view of both chamber |
| Arduino | Arduino Uno is connected to the computer and is placed outside of the enclosure box |
| Power supply | **External power supply remains off** during setup and wiring |
| Safety | No water is present near exposed electronic components |
| Fish | **No fish are introduced** before calibration and system verification |
| Gloves | Disposable gloves are worn during setup, calibration, fish handling, and cleanup to protect researchers and reduce contamination |

**Water level:**

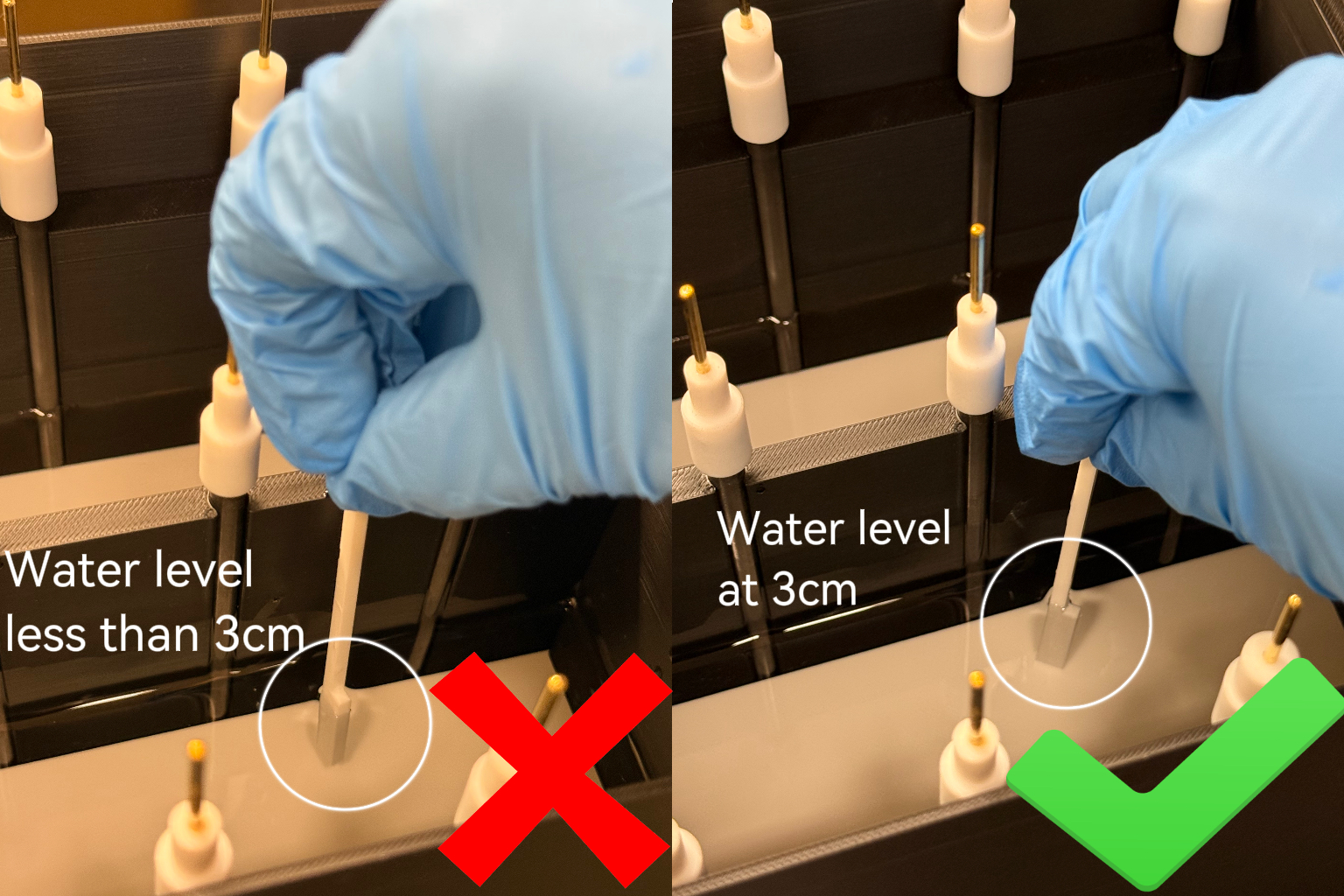

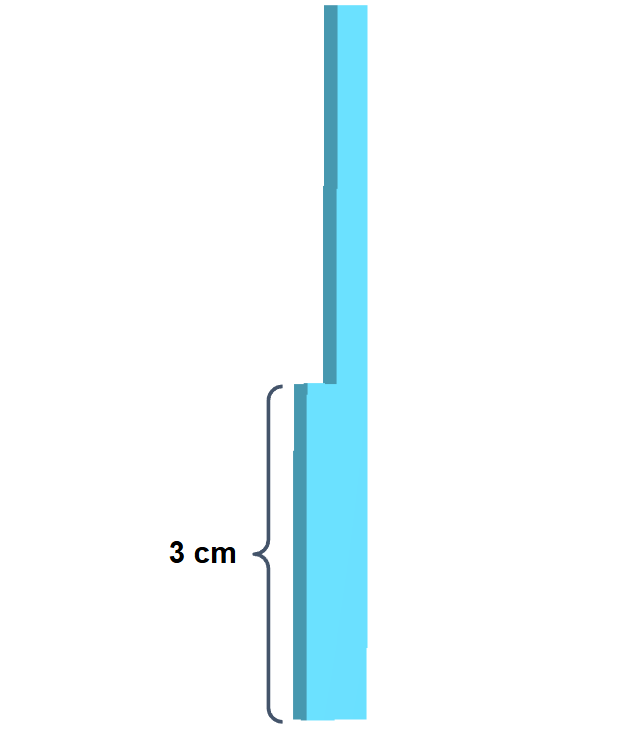

**Figure 1.** Water-level calibration using a 3D-printed depth ruler. The water depth in each chamber should be checked before calibration and behavioral testing. A 3D-printed ruler with a 3 cm reference mark is placed vertically into the chamber, with the base touching the chamber floor. The water level should reach or slightly exceed the 3 cm mark. The 3D-printing model file for the ruler is provided as an *.stl* file in the supplemental materials.

Each chamber should be filled with system water to a target depth of 3.0 cm before calibration and behavioral testing. Because exact water filling can be difficult in practice, researchers should use the provided 3D-printed water-level ruler to verify the depth. Place the ruler vertically into the chamber with the base touching the chamber floor. The water level should reach or slightly exceed the 3 cm reference mark. The water level should not be below 3.0 cm, because insufficient water depth may reduce electrode contact and affect stimulation consistency. Water levels that are too high should be avoided because they increase the chance that fish jump out of the chamber. **The same water level range should be maintained across experiments** because water depth can affect electrode contact, electrical stimulation consistency, and zebrafish swimming behavior. If a fish jumps out of its chamber or crosses into another chamber during the experiment, stop both the Arduino-controlled protocol and video recording immediately. The trial should be documented and excluded from downstream behavioral analysis.

**Electrode and Relay Wiring:**

Before uploading the experimental protocol code or introducing fish, connect the electrode stimulation circuit and verify that all wiring is secure. The external power supply should remain **off** while the relay module, electrode, and alligator clips are being connected. All exposed clips and wires should be positioned outside the water-contact area to prevent unintended short circuits or accidental contact with the aquatic environment.

The stimulation circuit connects the external DC power supply, relay module, and graphite rod electrodes. In this setup, the NO terminal of the relay is connected to the external power supply ground pathway, the COM terminal is connected to the positive electrode pathway, and the NC terminal is connected to the positive output of the external DC power supply (Figure 2.). The positive electrode pathway delivers stimulation to the outer positive electrode rows, while the shared central electrode row serves as the negative electrode pathway for both chambers.

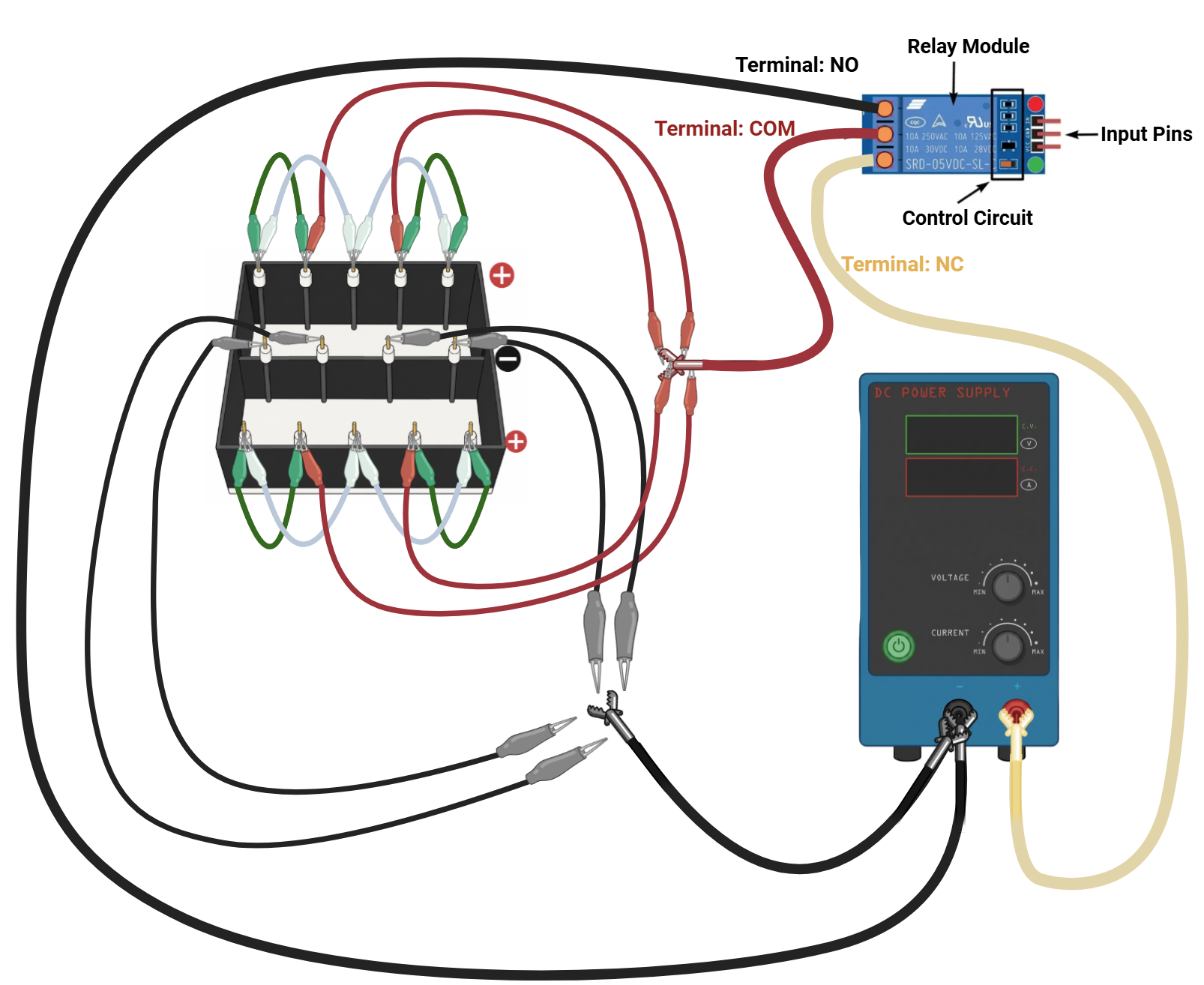

**Figure 2.** Wiring configuration for electrode stimulation in the arena. This schematic illustrates the wiring arrangement used to connect the relay module, external DC power supply, and graphite rod electrodes in the behavioral arena.

After wiring, check that:

a. Alligator clip stability and separation: each alligator clip should be firmly attached to its corresponding electrode rod. Clips should not touch one another, and wires should be arranged so that they do not pull on the electrodes, interfere with the arena, or enter the water. Any loose, unstable, or unclear connection should be corrected before calibration.
b. Camera visibility: alligator clips and wires should not obstruct the top-down camera view of either chamber. After wiring, confirm through the recording software that both chambers remain **fully visible** and that fish movement can be captured clearly throughout the experiment.

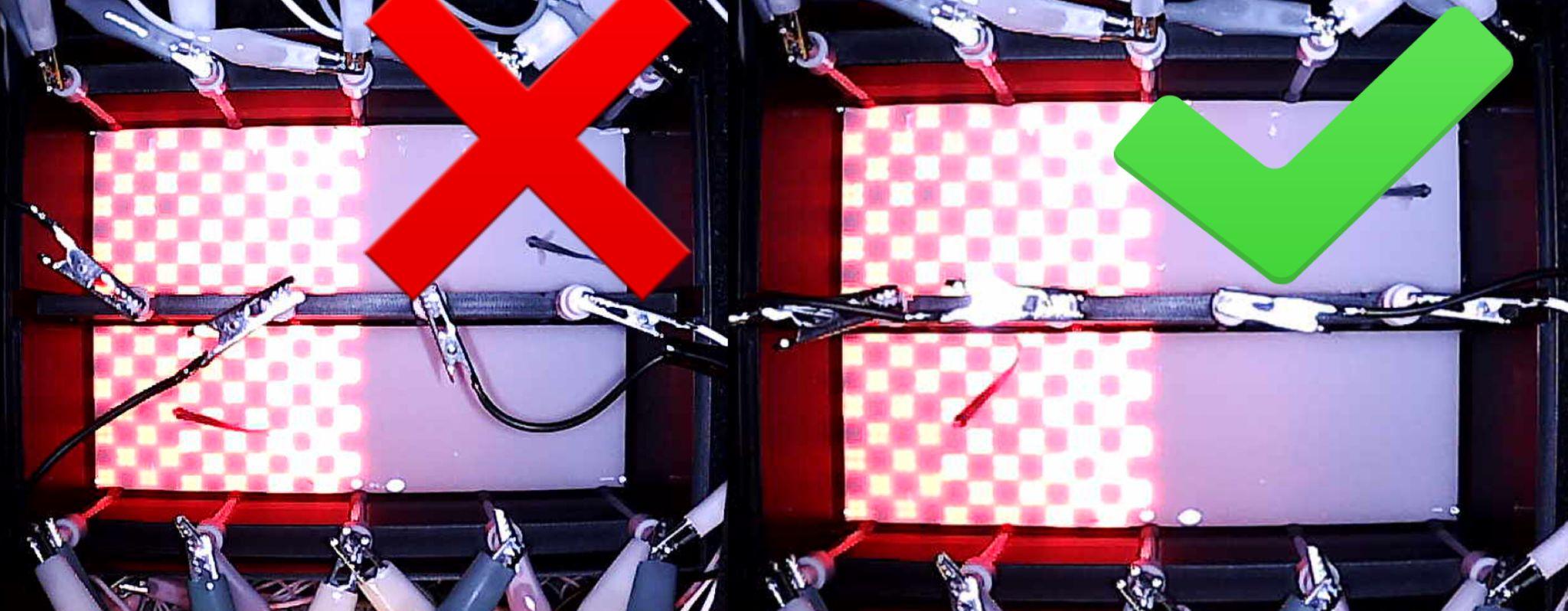

**Figure 3**. Camera Visibility Check. **Left**: incorrect wiring arrangement, where alligator clips and wires obstruct the top-down camera view. **Right**: correct arrangement, where both chambers remain visible and fish movement can be recorded clearly.

*This image is provided only to illustrate poor versus proper camera visibility;* ***no fish should be introduced during the wiring and visibility-check stage****. Researchers should check camera visibility frequently before and during setup to ensure that the recording view remains unobstructed.*

c. Electrical isolation from the Arduino: The negative electrodes pathway should not be connected to the Arduino board, including Arduino GND or any Arduino input/output pin. Connecting the stimulation pathway directly to the Arduino may allow excessive current to pass through the microcontroller and damage the board.
d. Final setup: after filling the chambers, inspect the arena and surrounding setup for leaks, water droplets, or loose connections. The exterior of the arena and the area around the LED matrix should remain dry. Confirm that the electrode connections are stable, the LED panels are aligned with the arena floor, the Arduino is connected to the computer, and the infrared camera is recognized by the recording software. **No fish should be introduced at this stage**. The system should first pass electrical calibration and visual stimulus verification before any behavioral trial begins.

**Voltage Calibration Before Experiment:**

This section should focus on verifying that the electrical stimulation circuit is working correctly before fish are introduced. The calibration step uses a separate Arduino calibration code, not the experimental training code. The purpose is to confirm that voltage delivery across each positive–negative electrode pair is consistent and within the expected range before running the behavioral experiment.

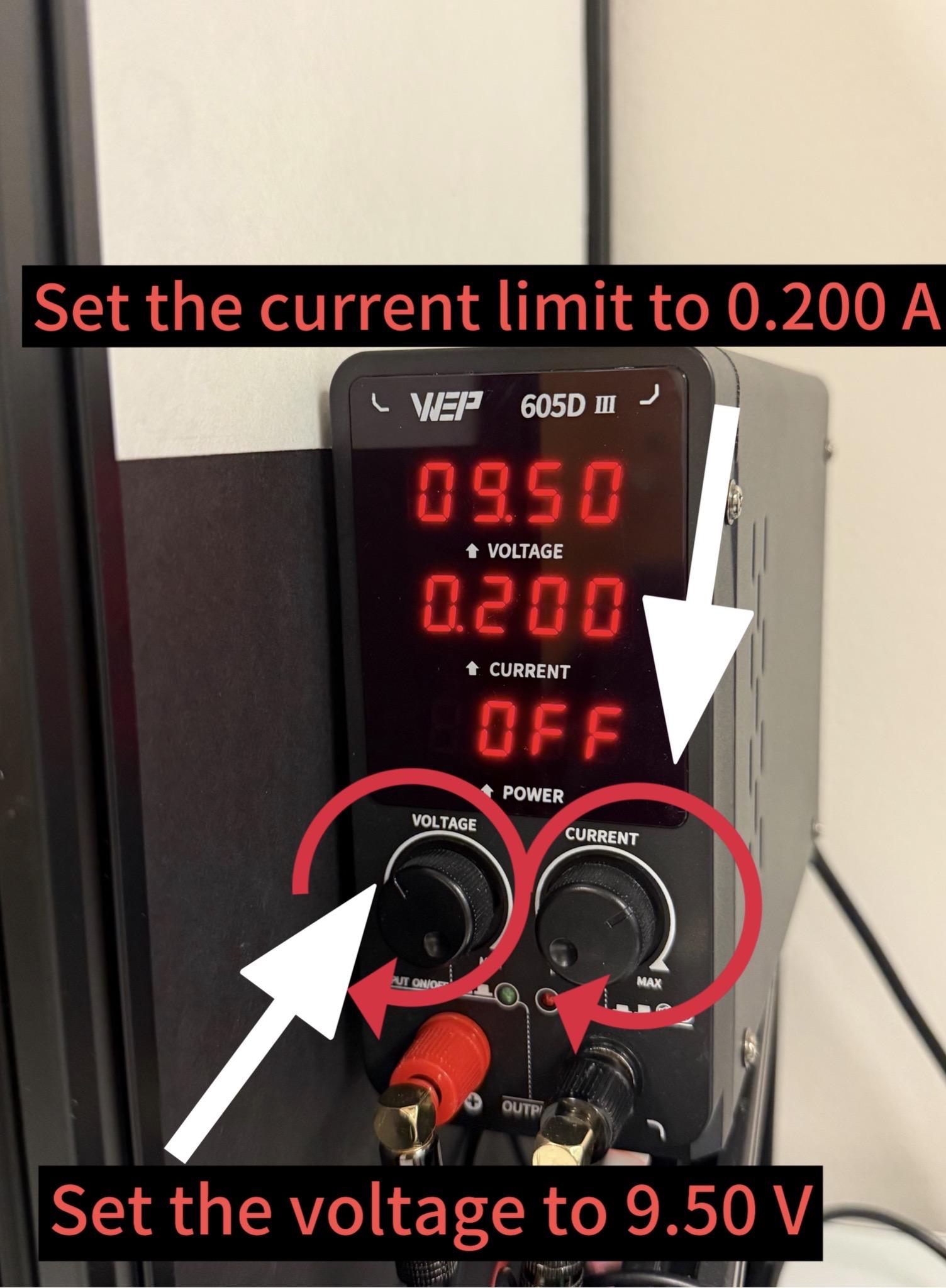

**Figure 4.** External Power Supply Settings for Voltage Calibration. The external DC power supply was set to 9.50 V with a current limit of 0.200 A. Researchers should use the voltage and current adjustment knobs to fine-tune these values before enabling output power.

During voltage calibration, the external DC power supply should be turned on and set to 9.5 V, with the current limit set to 0.200 A. The voltage setting controls the stimulation voltage delivered through the relay-switched electrode circuit, while the current setting acts as an upper safety limit. Researchers should upload the calibration code to the Arduino Uno.

To upload the calibration code to Arduino:

a. Download the .ino file named ‘ElectrodeVoltageManualTest’ on your PC or laptop

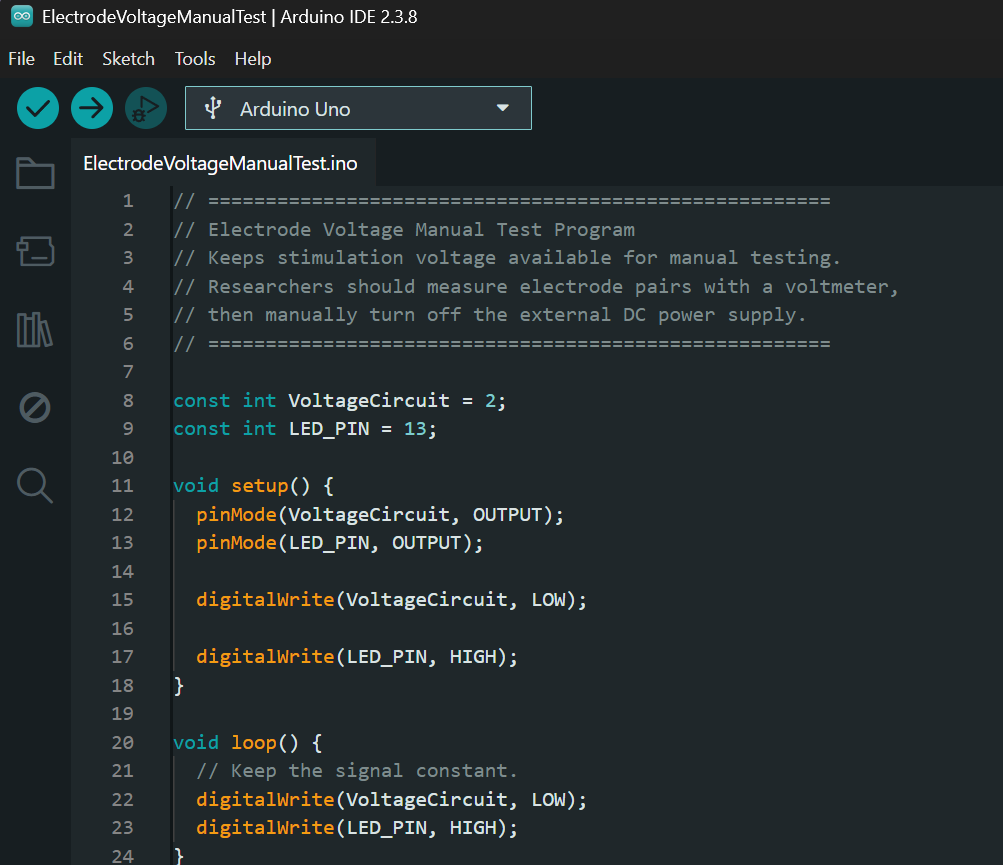

*Find the ino. file in the directory and open it in Arduino IDE*

b. Select the the Arduino board you want to use

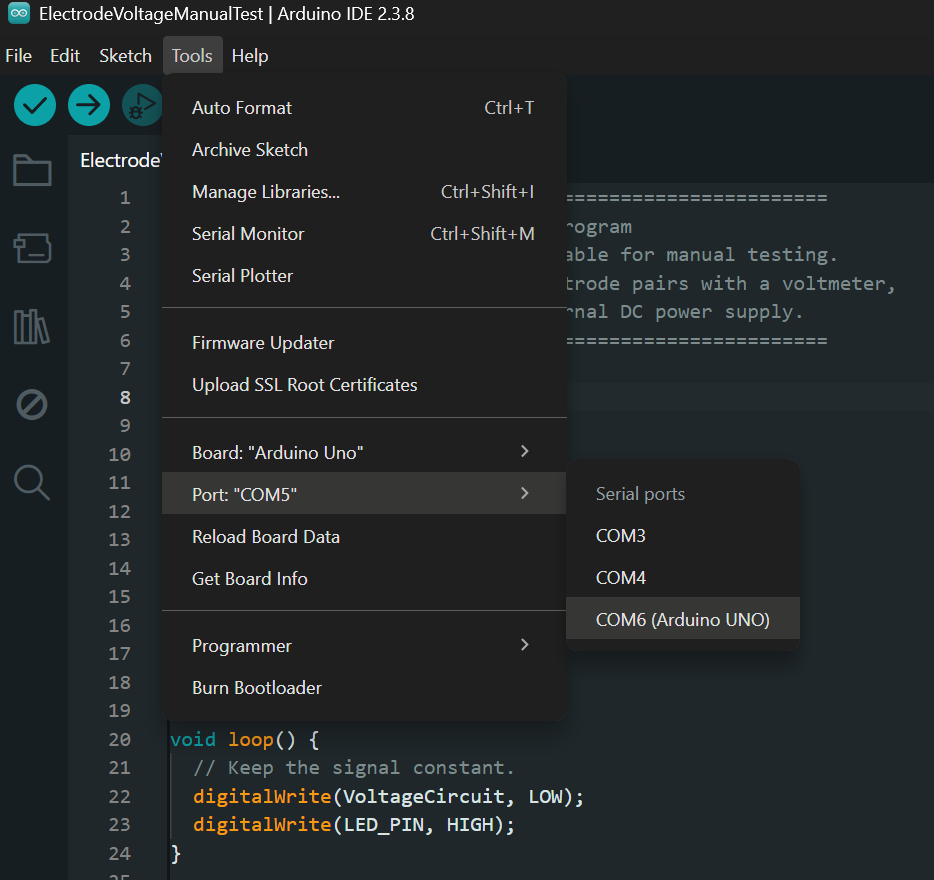

*Select ‘Tools’ -> then select ‘Port’ -> then choose the Arduino board you are using*

c. Upload the voltage calibration codes to the Arduino board

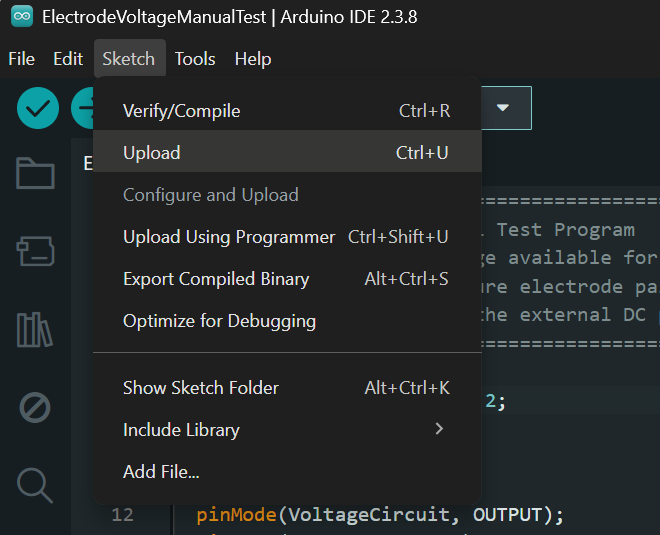

*Click ‘Sketch’ -> then ‘Upload’ the codes the Arduino Uno board*

d. Done compiling

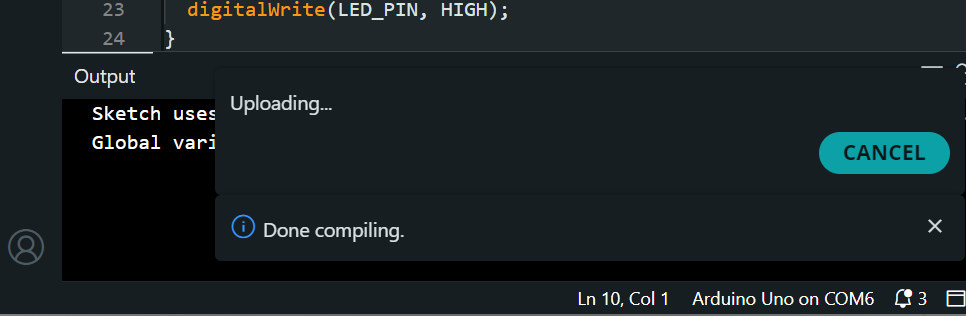

After uploading the voltage calibration code to the Arduino, turn on the external power supply output by pressing the voltage control on the power supply. Using a voltmeter in DC voltage mode, measure the voltage across each positive and negative electrodes pair. The measured voltage should be approximately 9.4–9.5 V, as shown in Figure 6. If the voltage measured from any electrode pair falls outside this range, do not proceed with the experiment until the wiring, relay connections, alligator clips, electrode contact, and power supply settings have been checked and corrected. **No fish should be introduced during this calibration step.** After the voltage calibration, **turn off the external power supply output.**

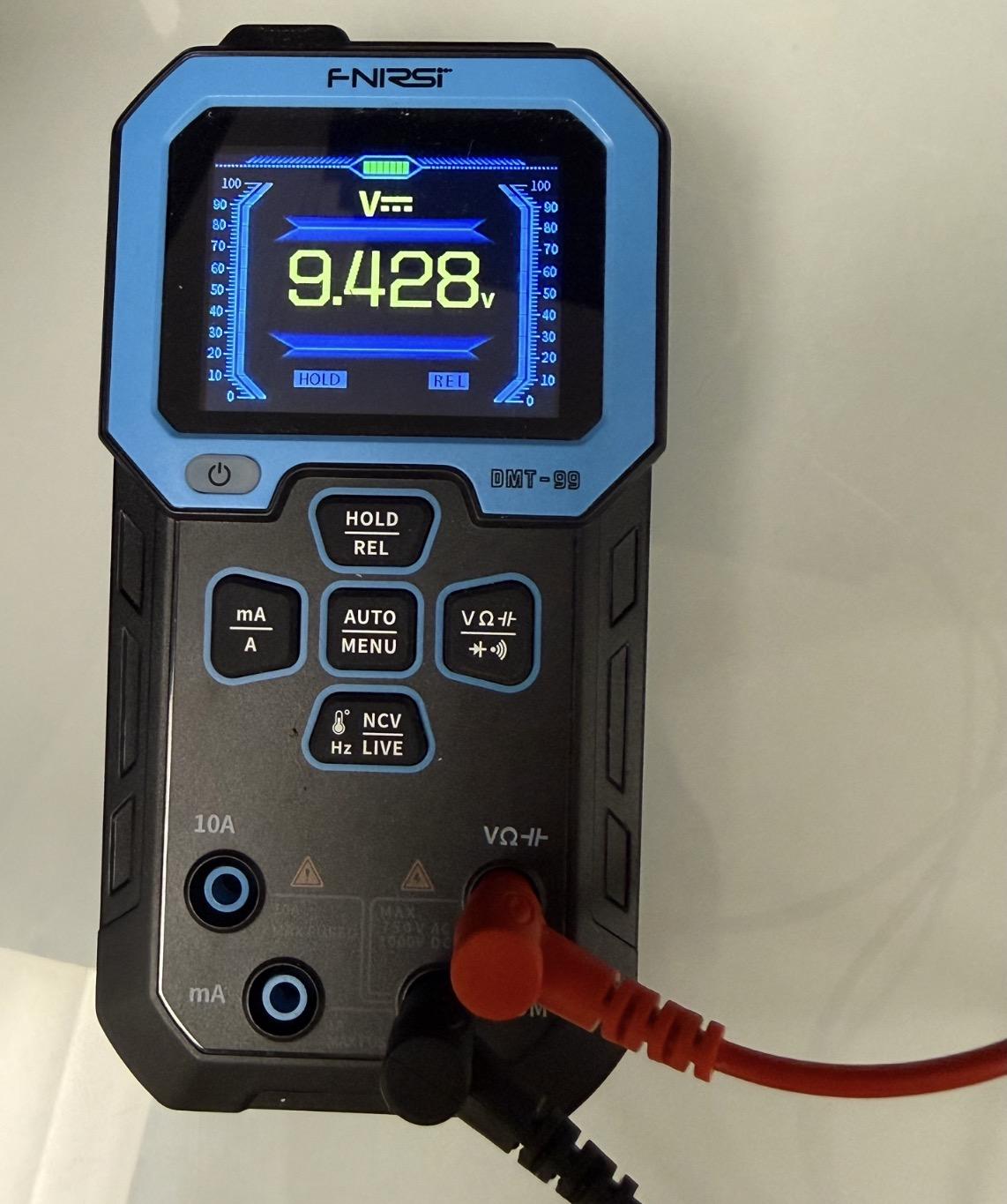

**Figure 6.** Voltage Measurement During Electrode Calibration.

Before starting to do the experiments, double check that (1) the water level is at the desired level, (2) all electrode and relay wiring is secure, (3) camera visibility is unobstructed, and (4) voltage calibration has passed. After the preparation steps are complete, the system is ready for the behavioral experiment.

**3. Starting the Experiment**

**Step1. Upload the experimental code to the Arduino**

Upload the experimental Arduino code file, *BehavioralExperimentCode.ino*, to the Arduino Uno using the Arduino IDE. This code is different from the voltage calibration code and is used to run the full behavioral experiment.

**Step2. Turn on the external power supply**

Turn on the external DC power supply and confirm that it is still set to 9.50 V with the current limit set to 0.200 A.

**Step 3. Verify LED function and voltage stimulation**

After the experimental code is uploaded, the LED panels should briefly illuminate at the beginning of the program. Use this startup illumination as a visual check to confirm that all LED modules and pixels are functioning. If any LED module or pixel fails to light during this check, it will likely not work during the experiment. In that case, **re-upload** the experimental code and inspect the LED wiring before continuing.

Researchers may also use the optional reference LED indicator to confirm that the stimulation circuit is active during programmed shock periods. The reference LED should be connected in series with a resistor, with its positive side connected to the positive electrode pathway and its negative side connected to the external power supply negative/GND pathway.

*If needed, restart the Arduino program using the white reset button on the Arduino board. Once the LED startup check and stimulation-circuit check are completed successfully, proceed to fish loading.*

**Step 4. Introduce fish into the arena, close the enclosure, and turn off room lights**

The display should begin counting down from 03:01. Researchers have **only 1 minute** to introduce the fish before the recording period begins.

To make fish loading faster and more consistent, prepare the fish in separate transfer containers before this step, with one fish per container. Gently introduce one fish into each chamber of the arena. Avoid splashing water onto external wiring or disturbing the electrode positions.

After both fish are loaded, close the door of the enclosure box and turn off the room lights. The enclosure should remain closed during the experiment to reduce external light, sound, and environmental disturbance. Confirm that the fish remain in their assigned chambers before proceeding to video recording.

**Step 5. Start video recording**

Open the video recording software before fish loading or immediately after fish are placed into the arena. You should monitor the time display and start recording **immediately at 03:00.** This timing ensures that the video recording begins at the start of the experimental protocol and captures the full 180 minutes behavioral session for downstream analysis.

**Step 6.** **Monitor the experiment**

During the experiment, avoid opening the enclosure or moving the setup unless animal safety requires intervention. Periodically check the recording software to confirm that both chambers remain visible and that each fish remains in its assigned chamber. If a fish jumps out of its chamber, crosses into the other chamber, or if video recording or power is interrupted, stop the Arduino protocol and video recording immediately. Document the issue clearly, including the time and reason for interruption, and exclude the affected trial from downstream analysis.

**Step 7. End the experiment and save files**

At the end of the 180-min protocol, stop the video recording and turn off the external power supply. Save the full recording using a consistent file name that includes the fish ID, strain, sex, chamber position, date of birth, and any relevant experimental notes. Record any abnormalities observed during the experiment, such as fish movement between chambers, tracking obstruction, power interruption, or unusual behavior.

**Step 8. Clean the system**

After the experiment is finished and the external power supply has been turned off, disconnect the wiring from the arena and electrodes. Remove the fish according to the laboratory’s animal-handling procedures. Rinse the arena and graphite electrodes with clean water to remove debris or residue. Wipe the outside of the arena and electrode rods gently, then place them in a well-ventilated area to dry completely before the next experiment.

*Do not rinse or wet the Arduino, relay module, LED matrix modules, power supply, timer display, camera, or any other electronic components. Only the water-contact components, such as the arena and electrodes, should be cleaned directly. Make sure all components are fully dry before reconnecting the system.*

Detailed instructions for video segmentation, Tracker-based position extraction, and data export are provided in **Supplementary Material 3: Tracker-Based Zebrafish Position Extraction Workflow.**

**Simplified Experimental Checklist**

This checklist summarizes the major preparation and experimental steps for running one zebrafish conditioning experiment. Researchers may print this page and check each item during setup and operation.

*This checklist is not a replacement for the full protocol; users should read the complete Supplementary Material 2 before running the experiment.*

**Preparation:**

| Step | Required Action | Check |
| --- | --- | --- |
| Gloves | Wear disposable gloves during setup, calibration, fish handling, and cleanup. |  |
| System assembly | Confirm that the arena, enclosure, LED modules, electrodes, Arduino, relay, power supply, timer, and camera are assembled correctly. |  |
| Power supply | Keep the external power supply output **OFF** during water filling and wiring. |  |
| Water level | Fill each chamber to approximately 3.0 cm. Water level should be ≥3.0 cm. |  |
| Electrode wiring | Confirm that alligator clips are firmly attached to the electrode rods and that clips are not touching each other. |  |
| Camera visibility | Confirm that alligator clips and wires do not block the top-down camera view of either chamber. |  |
| Voltage calibration | (1) Set the external power supply to 9.5 V and current limit to 0.200 A  (2) Upload the voltage calibration code to the Arduino.  (3) Turn on power supply output and measure voltage across each positive–negative electrode pair.  *Expected reading: approximately 9.4–9.5 V.* |  |
| Fish status | Confirm that no fish are in the arena during preparation, wiring, calibration, or LED verification. |  |

**Starting the experiment:**

| Procedures | Check |
| --- | --- |
| Step 1. Upload the experimental code to the Arduino |  |
| Step 2. Turn on the external power supply |  |
| Step 3. Verify LED function and voltage stimulation  *If needed, restart the Arduino program using the white reset button on the Arduino board.* |  |
| Step 4. Introduce fish into the arena, close the enclosure, and turn off room lights  *You only have 1 minute to complete this step* |  |
| Step 5. Start video recording (start right at 3:00) |  |
| Step 6. Monitor the experiment |  |
| Step 7. End the experiment and save files |  |
| Step 8. Clean the system |  |
