## Supplementary Material 3 for "zAcademy: an open-source programmable conditioning arena reveals sex- and genotype-dependent avoidance learning in adult zebrafish"

**Tracker-Based Zebrafish Position Extraction Workflow**

**1. Overview**:

This guide describes how to use ‘Tracker’ to extract zebrafish position data from behavioral videos recorded during the three hours conditioning protocol. The exported position data are later used in RStudio to calculate the performance index (PI) and quantify spatial preference across baseline and post training analysis periods.

**2. Preparation**:

**Video Recording Options**

**Option 1. Full three-hour recording**

The simplest approach is to record the full 180-min behavioral experiment as one continuous video file. After the experiment, the full video can be divided into shorter analysis segments corresponding to the baseline and post-training analysis windows.

Software:

To download OBS Studio (optional): <https://obsproject.com/>

To download LosslessCut (required): <https://losslesscut.app/>

*OBS Studio is optional when the computer’s built-in camera application can reliably record the full three-hour experiment. If the built-in application cannot support continuous recording, use OBS Studio or another recording program capable of capturing the required video duration.*

Before importing videos into Tracker, cut the full three hours behavioral recording into shorter clips corresponding to the time windows selected for analysis. This step makes tracking more manageable and reduces the risk of software lag or loading errors caused by processing a long video file. Video clipping was performed using ‘*LosslessCut*’ which can quickly cut long videos into short clips without re-encoding, so the video quality stays the same and the export is fast. For this study, clips should be generated for the two baseline reference windows and each post training memory test.

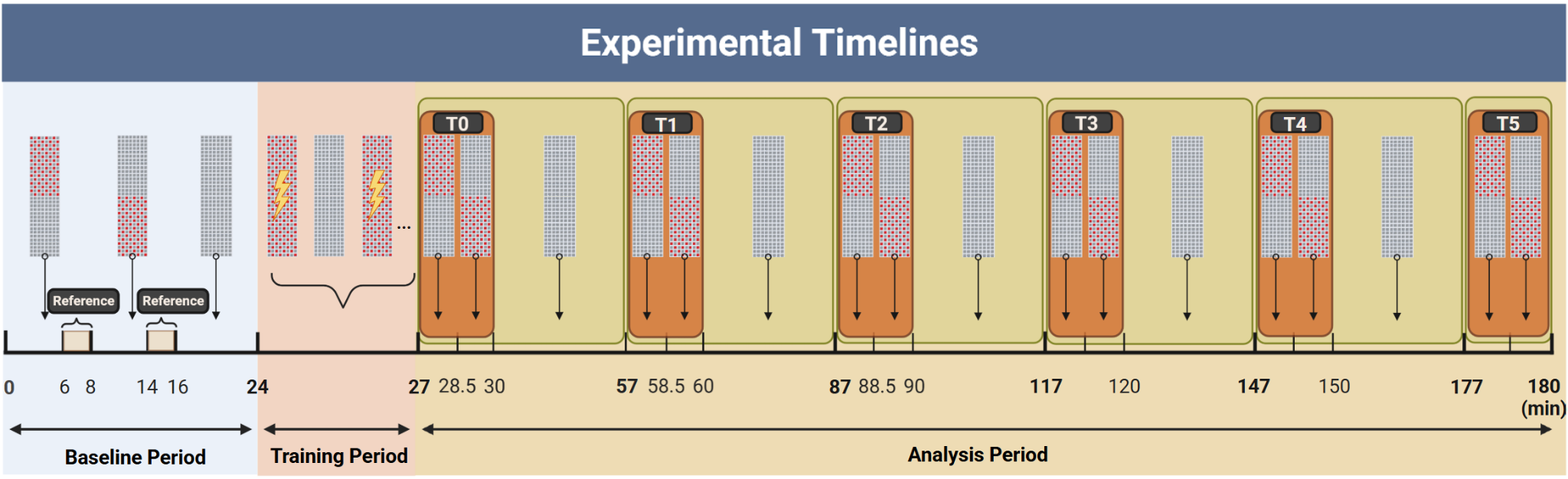

**Figure 1.** Experimental timeline of the three-hour zebrafish conditioning protocol.

| **Video segment** | **Time window in full recording** |
| --- | --- |
| Baseline 1 | 6-8 min |
| Baseline 2 | 14-16 min |
| T0 | 27-30 min |
| T1 | 57-60 min |
| T2 | 87-90 min |
| T3 | 117-120 min |
| T4 | 147-150 min |
| T5 | 177-180 min |

**Table 1.** Selected video time windows for analysis.

After clipping, import each short video clip into Tracker **separately**. Use consistent file names so that each exported coordinate file can be matched to the correct fish, strain, and time window. For example:

WTTest1T0.mp4

WT：The strain

Test1: The Test number

T0: The video segment

*This method (Option 1) is easy to understand and provides a complete backup of the behavioral session. However, the file size can be large, and some built-in camera recording applications may not support uninterrupted three-hour recording.*

**Option 2. Automated recording using OBS Studio, obs-websocket, and Python**

The second approach uses OBS Studio for video recording and a Python script to control when recording starts and stops. In this workflow, OBS receives the camera feed, while Python communicates with OBS through obs-websocket and automatically records only the selected time windows. This method is useful when the computer cannot easily store a full three-hour recording.

Software:

To download Python (required): [https://www.python.org/downloads/](https://www.python.org/downloads/?utm_source=chatgpt.com)

To download Visual studio codes (required): <https://code.visualstudio.com/>

To download OBS Studio (required): [https://obsproject.com/](https://obsproject.com/?utm_source=chatgpt.com)

To download the Python code (required): [Github code](https://github.com/IlluminationLi/ZebrafishAutomatedRecord.git)

Procedures:

**Step 1.** Connect the Camera

Connect the camera to the computer and open OBS Studio.

Confirm that:

a. The camera image is visible.
b. The correct camera source is selected.
c. The video quality and framing are correct.

**Step 2.** Check OBS WebSocket Connection

In OBS: Select ‘Tools’ -> ‘WebSocket Server Settings’

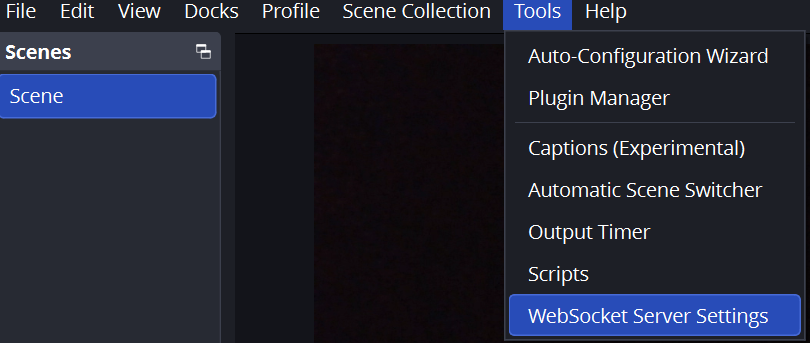

Then, Select ‘Enable WebSocket server’ and ‘Server Port’ 4455

Set your own ‘Server Password’ (for example: zebrafish)

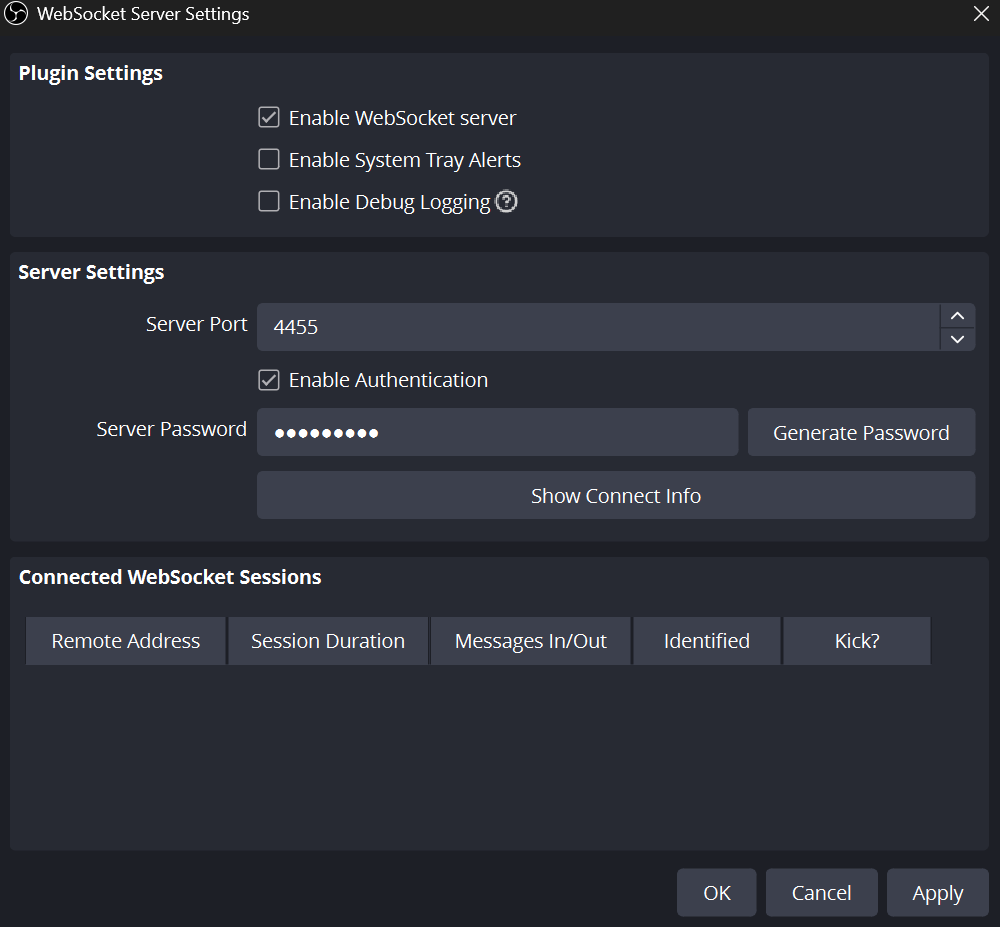

*Do not change these settings and remember the password you just set*

Go to config,py to change the PASSWORD to the one you just set

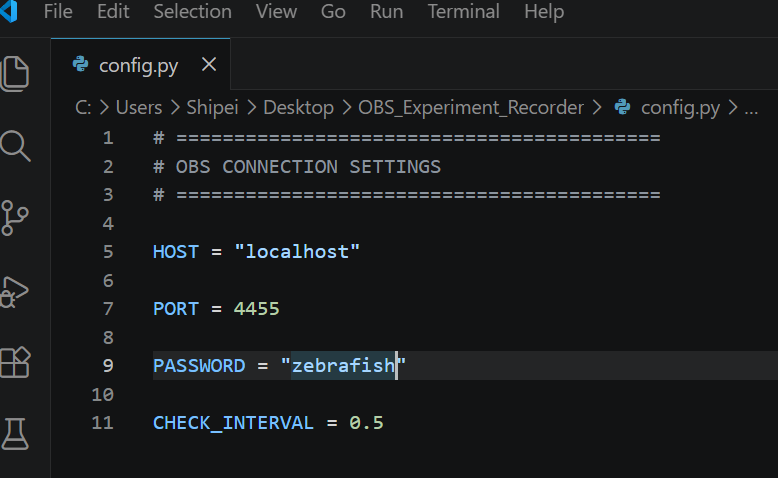

**Step 3.** Open the Recording Program

1) Download the Python code from the github link (see above).
2) Then, open the folder, OBS_Experiment_Recorder, using Visual Studio Code
3) In the terminal window, type the directory of the downloaded file:

*cd "C:\Users\UserName\Desktop\OBS_Experiment_Recorder"*

4) Then, type:

*dir*

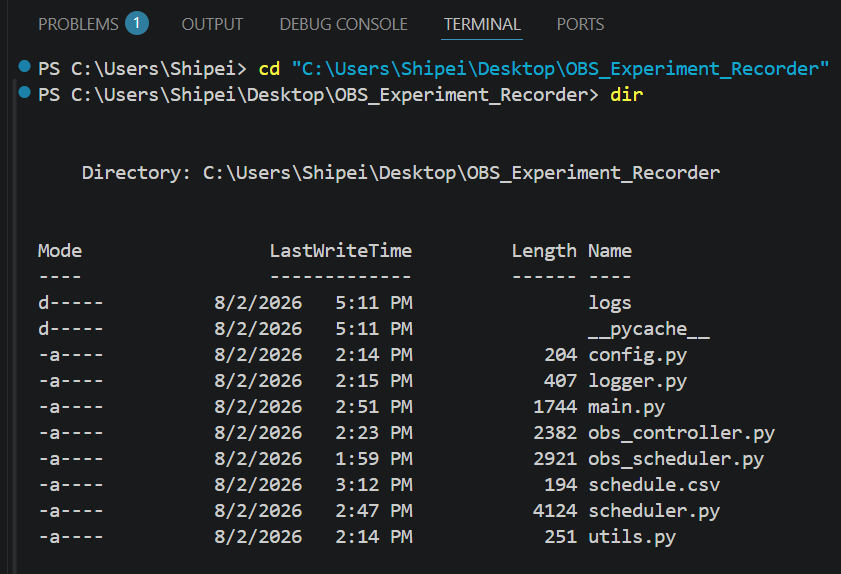

*You should see these files listed in this directory*

5) Download the obsws-python package (if you haven’t).

In the terminal, type:

python -m pip install obsws-python

6) Next, in the terminal, type:

*python main.py*

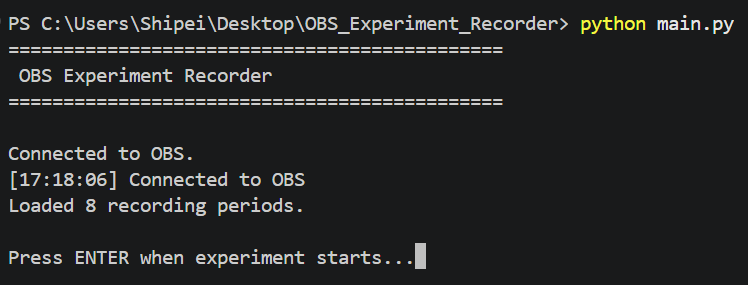

7) Press ENTER on your keyboard when you start experiments.

Begin the behavioral experiment according to Supplementary Material 2. The Python recording script should be started at the same defined time point used for synchronization when the timer display reaches 03:00 and the behavioral recording period begins.

**Important:** Do NOT press Enter until the actual experiment begins. When the experiment starts and you press enter, the program will start the experiment timer and start recording.

After this point:

- Do not close the terminal
- Do not close OBS
- Do not shut down the computer

The system will **automatically** control recording

**Step 4.** Find recorded videos after the experiment

In OBS, select ‘File’ -> ‘Show recording’’

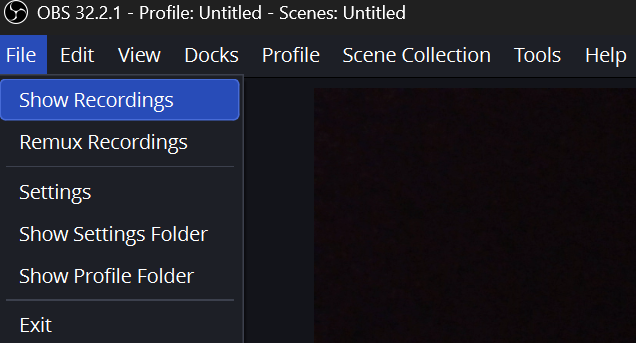

**Step 5.** Name your videos

Use consistent file names so that each exported coordinate file can be matched to the correct fish, strain, and time window.

For example:

WTTest1T0.mp4

WT：The strain

Test1: The test number

T0: The video segment

Automated recording reduces file size and removes the need to manually cut a three-hour video, but it also means that only selected analysis windows are saved. If only automated segment recording is used, researchers should verify that all required segments are saved before ending the experimental session. The automated recording workflow only controls video capture. It does not control the Arduino behavioral protocol, LED stimulus presentation, relay activation, or electrical stimulation. These functions are controlled separately by the Arduino program described in Supplementary Material 1.

**3. Tracker Analysis**:

To download Tracker: <https://opensourcephysics.github.io/tracker-website/>

**1. Video clipping before Tracker import**

Cut the full three hours behavioral recording into shorter clips corresponding to the time windows selected for analysis. Each clipped file should be named according to fish strain, test number, and time window before being imported into Tracker for position extraction.

**2. Open Tracker and import video**

Open Tracker and import the behavioral video file. Confirm that the video loads correctly and that the full arena is visible throughout the recording.

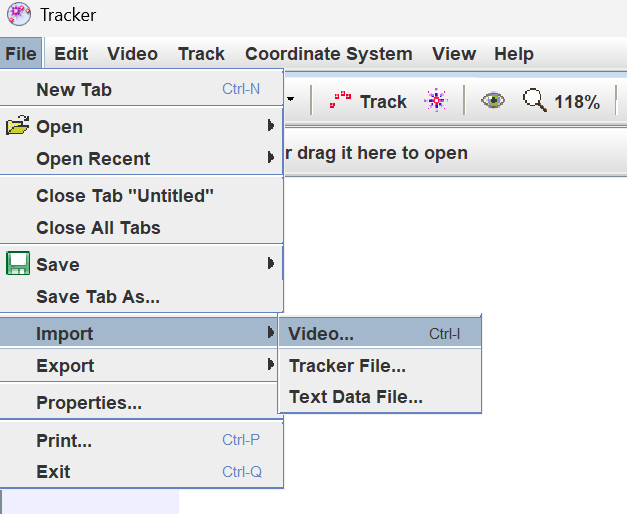

**3. Set up your coordinate system**

Before tracking, set a consistent coordinate system for the arena. The x-axis should run along the length of the chamber, and the y-axis should run across the chamber width. The origin should be placed consistently across videos, preferably at one corner of the chamber. Keeping the same coordinate orientation across trials is important because the exported coordinates will be used to classify whether the fish is in the illuminated or dark region.

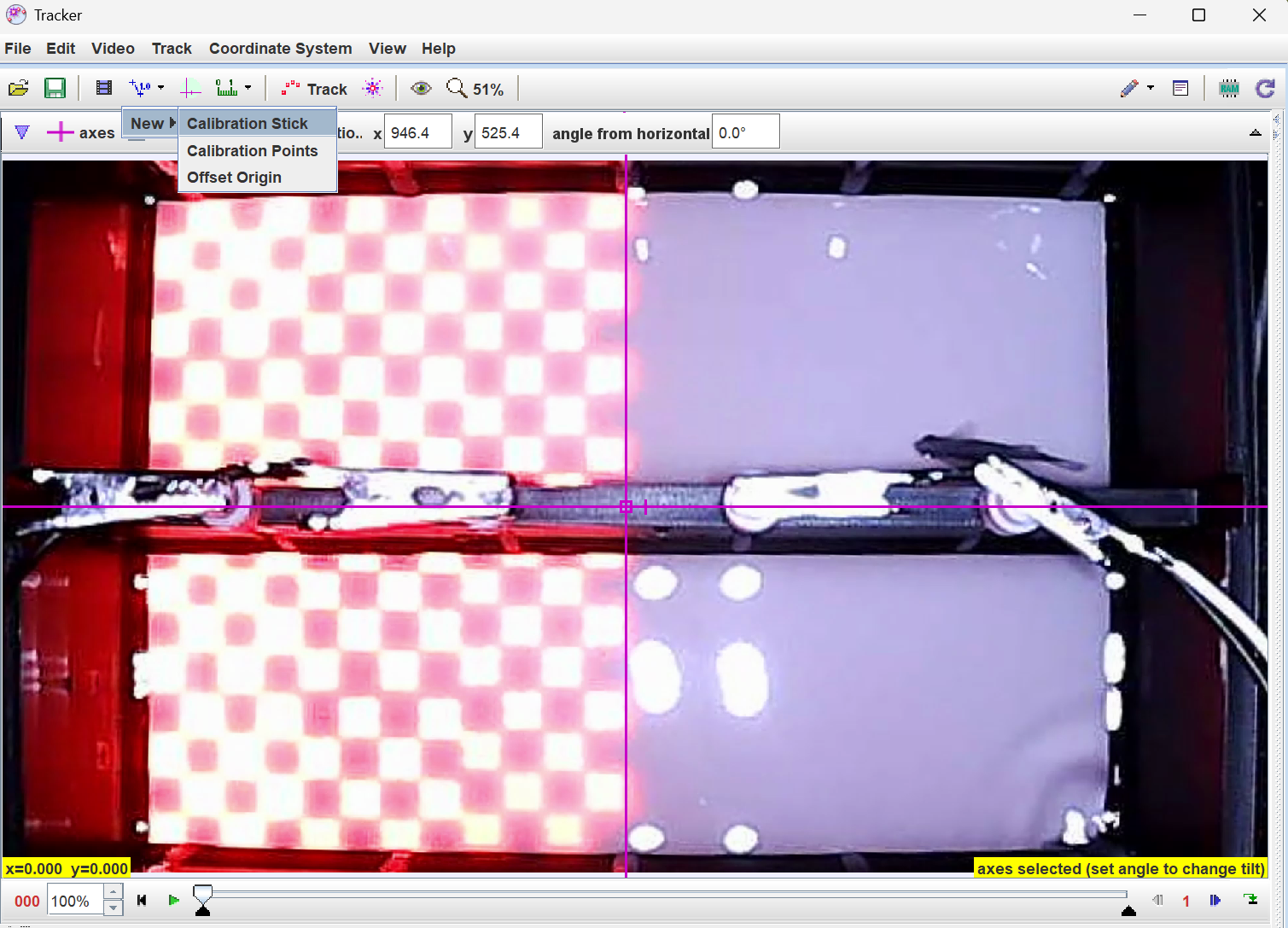

**4. Calibration stick**

Use the Calibration Stick tool to define the spatial scale of the video. Place the calibration stick along a known distance in the arena, such as the chamber length, and enter the corresponding real world measurement. This step converts the position data from pixels into physical units and ensures that movement measurements are comparable across videos. The same calibration procedure should be applied consistently to all video segments before tracking the fish position.

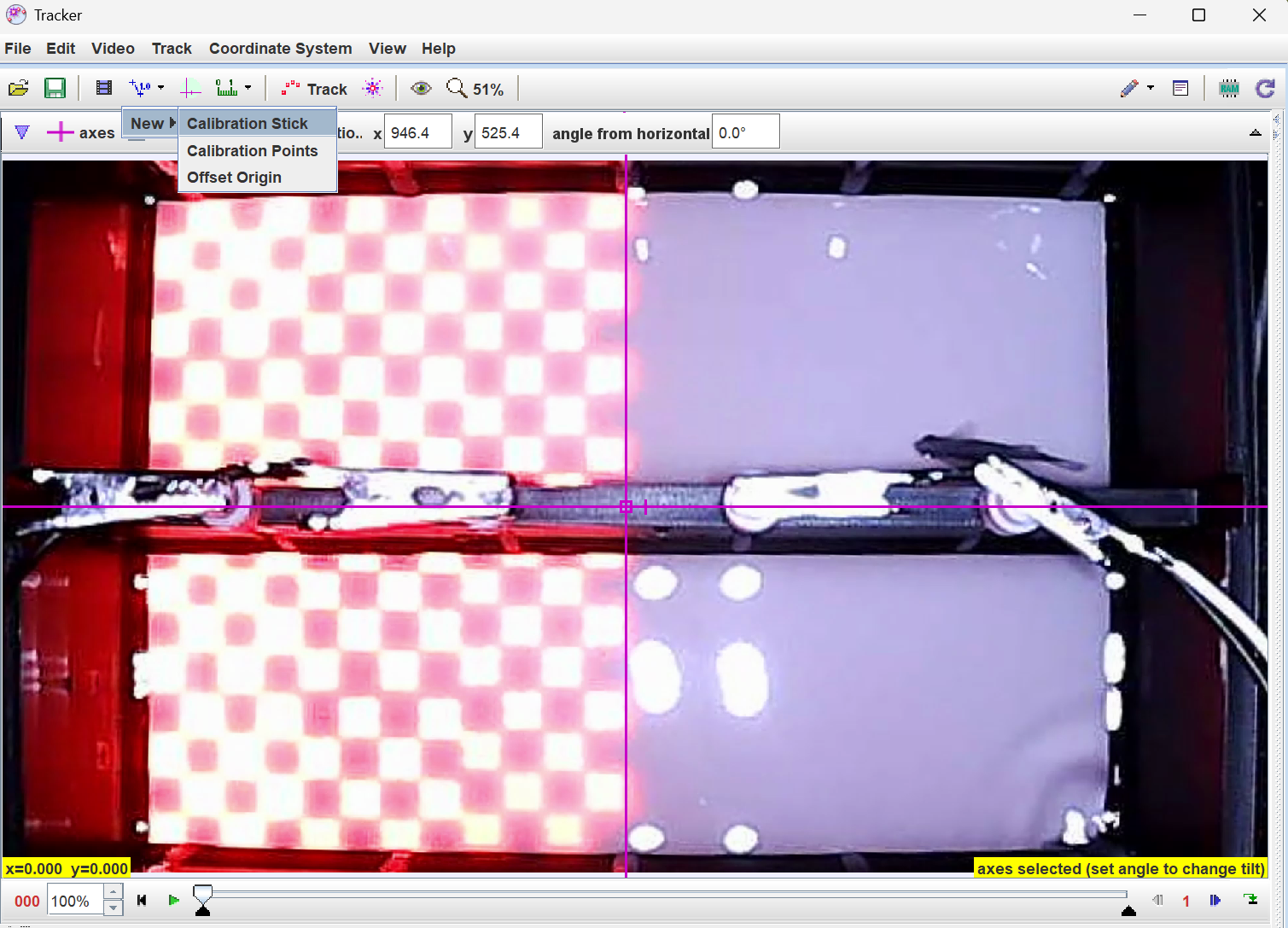

**5. Define the frame sampling interval**

The frame sampling interval should also be set before tracking. In this study, the original videos were recorded at **30** frames per second, and Tracker was set to advance 30 frames per tracking step. As a result, one frame was analyzed per second, corresponding to 60 analyzed frames per minute. This sampling rate was used for downstream PI calculation, where two 2-min baseline windows produced 240 analyzed frames and each 3-min memory test produced 180 analyzed frames.

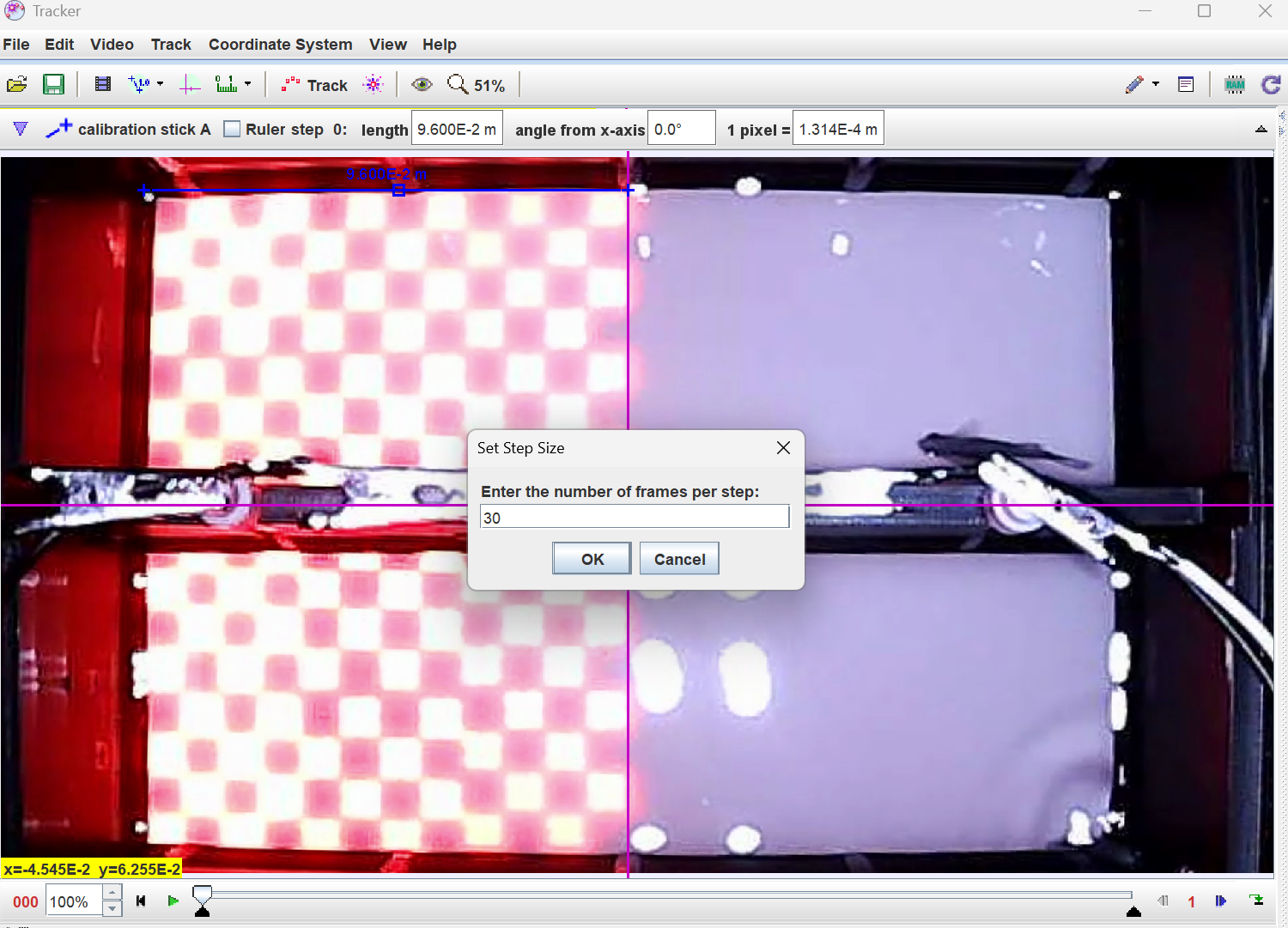

**6. Define the objectives**

Create a point mass to represent the zebrafish. Use the center of the fish body as the tracked point. If automatic tracking works reliably, use the autotracker function to follow the fish across frames. If tracking fails due to reflections, low contrast, overlap with chamber edges, or sudden movement, manually correct the position in selected intervals.

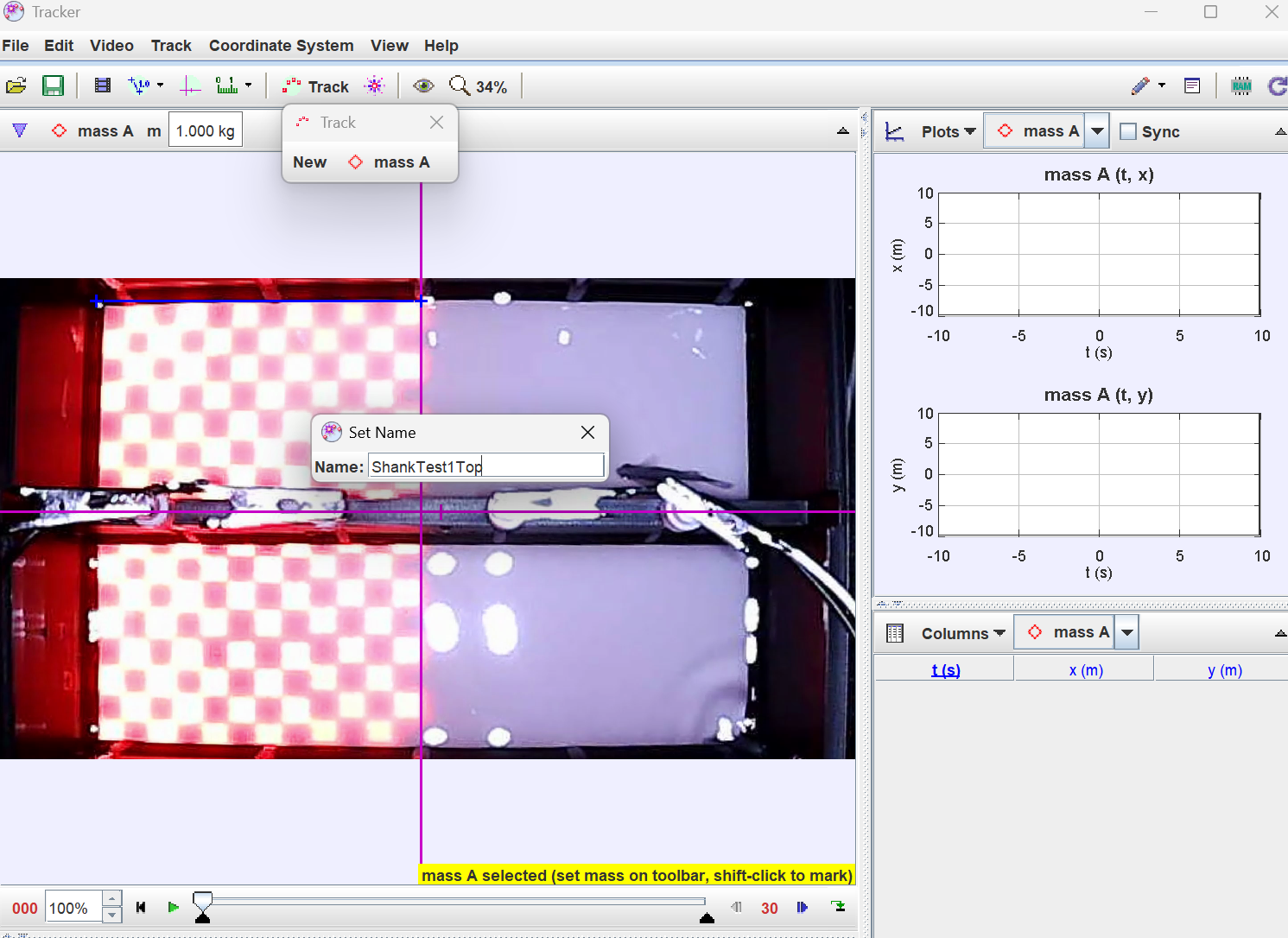

**7. Export the data**

After tracking is complete, export the position data as a .txt text file. Each exported file should be named consistently so that the strain, test number, time segment, and chamber location can be identified directly from the file name. A recommended naming format is:

StrainTestNumberTimeSegmentChamber.txt

*For example:*

*WTTest1T0Top.txt*

*In this example, WT represents the wild-type strain, Test1 represents the first experimental trial, T0 represents the analyzed time segment, and Top indicates that the fish was located in the upper chamber of the dual-chamber arena. The same naming structure should be used for all exported files to ensure that each position dataset can be matched correctly during downstream analysis in RStudio.*

**8. Save the *.trk* file**

After exporting the *.txt* file, save the Tracker project as a .trk file using the same naming structure. This file preserves the tracking setup, including the coordinate system, calibration scale, frame sampling interval, and tracked position data. Saving the *.trk* file allows the analysis to be reviewed, corrected, or repeated later if needed.

**9. Compute PI scores in RStudio**

Convert the exported *.txt* file into a *.csv* file while preserving the original column structure and file name information. The converted *.csv* file should then be imported into RStudio for PI score calculation. In RStudio, the position data are assigned with scores according to whether the fish is located in the dark region or the illuminated region over different time windows, and the frame-by-frame scores are averaged to compute PI.
