## Supplementary Material 4 for "zAcademy: an open-source programmable conditioning arena reveals sex- and genotype-dependent avoidance learning in adult zebrafish"

**Troubleshooting Guide for the Zebrafish Conditioning System**

**1. Overview**

This supplementary material provides troubleshooting guidance for common problems that may occur when assembling, calibrating, operating, or analyzing experiments using the zebrafish conditioning system. This guide should be used together with Supplementary Material 1: Zebrafish Conditioning Arena Build Guide, Supplementary Material 2: Experimental Protocol, and Supplementary Material 3: Tracker-Based Zebrafish Position Extraction Workflow.

**Important:** for safety, the external power supply should be turned off before adjusting wiring, alligator clips, electrodes, Arduino connections, LED modules, or relay terminals. If a problem occurs during an active behavioral experiment, researchers should stop the Arduino protocol and video recording, document the issue, and exclude the affected trial from downstream analysis when necessary.

**2. Common Problems and Solutions**

**Problem 1:** Different 3D printers to print the arena

**Solution:**

Other 3D printers can be used as long as it can accurately print the provided *.stl* arena file. The system described here was printed using a Bambu H2D printer with black PLA, white PLA, and water-soluble Bambu PVA support material.

**Problem 2:** Arduino code does not upload

**Solution:**

(1) Confirm that Arduino Uno is selected as the board in the Arduino IDE and that the correct COM port is selected. If the upload still fails, reconnect the USB cable, try a different USB port, and restart the Arduino IDE if needed. Make sure the USB cable supports data transfer, not only charging.
(2) After clicking upload, allow the IDE enough time to compile and transfer the code. Do not disconnect the Arduino until the IDE reports “Done uploading.” Sometimes it will take a few minutes for the arduino board to read and process the codes after uploading, just wait a few minutes before it is ready to work.

**Problem 3:** Some LED pixels or modules do not light

**Solution:**

(1) Re-upload the experimental code, check Table 1 pin assignments **(in Supplementary Material 1**), and confirm that LED Area 1 and Area 2 are connected to the correct Arduino pins. Then, check jumper wires, and confirm that 5V and GND are properly distributed through the breadboard
(2) If only part of the LED display fails to light, the problem is often caused by a loose connector or unstable wiring. Disconnect and reconnect the terminal/header connection on the LED module, then restart the Arduino using the reset button.

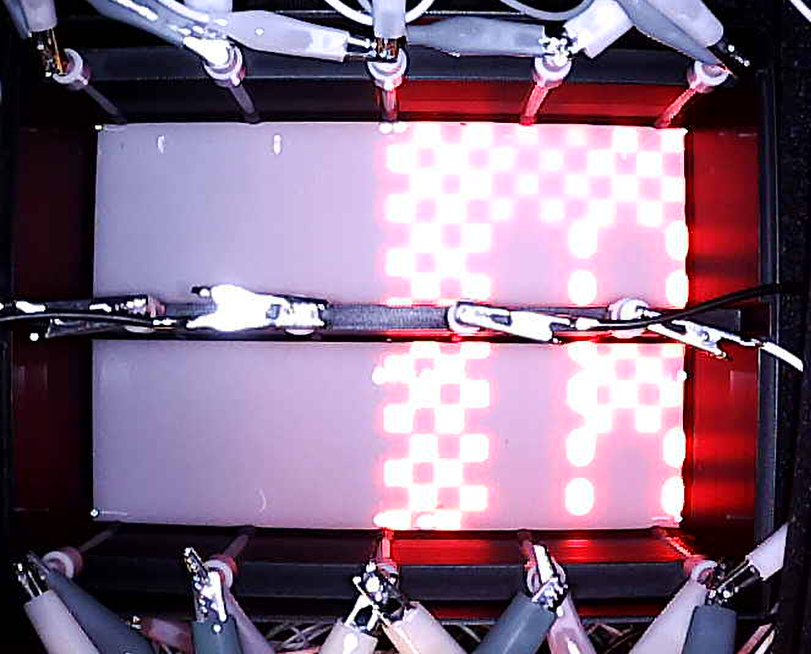

(3) If the LED pattern appears split across the wrong modules, as shown in the example image below, the LED modules may be arranged in the wrong order. In this case, correct the module order by switching the two middle LED modules so that the checkerboard pattern displays properly across the intended area.

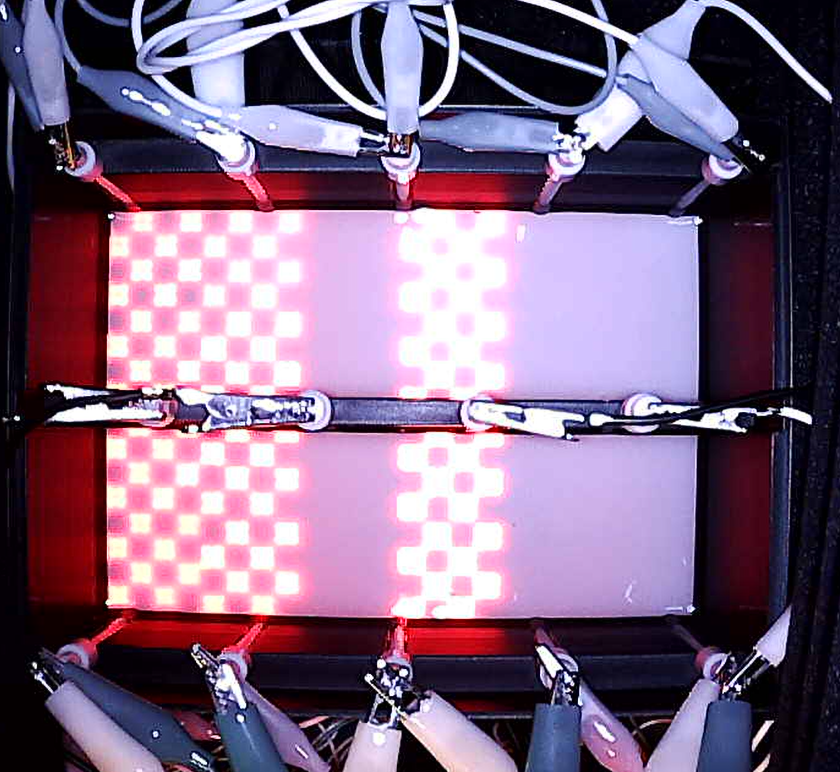

**Problem 4:** Relay does not activate

**Solution:**

(1) First, check the relay pin assignments listed in **Supplementary Material 1** and confirm that the relay input is connected to the correct Arduino pin. Verify that the relay module receives proper 5V and GND from the Arduino, and test whether the relay responds during the voltage calibration step. When the relay is activated, researchers should usually hear a small clicking sound or observe the relay indicator light changing state.
(2) If the relay does not switch even though the wiring and code are correct, the relay may be mechanically stuck. This can occur when the internal switch does not move properly between states, causing the relay to remain stuck on one side. In this case, gently tapping the blue relay housing may release the internal switch, but this should only be treated as a short-term troubleshooting step. If the relay continues to stick, switch inconsistently, or fail during calibration, replace it with a new relay module before running the experiment.

**Important.** Do not proceed with animal testing unless the relay switches reliably during calibration. The stimulation circuit should deliver voltage only during programmed shock periods and should remain inactive during non-shock periods

**Problem 5:** Reference LED does not light during shock check.

**Solution:**

Check LED polarity and confirm the 250 Ω resistor is in series.

**Problem 6:** Recording stops before 3 hours

**Solution**

(1) Use recording software that supports long recordings, connect the computer to power, and disable sleep mode.
(2) If the computer does not have enough storage space for a full 3-hour video, use the automated recording method described in **Supplementary Material 3**, which uses OBS Studio, obs-websocket, and Python to automatically record and save only the selected analysis windows.

**Problem 7:** Fish appears stressed before training

**Solution:**

Do not proceed with the trial if a fish shows signs of stress before training. Signs may include abnormal swimming, prolonged immobility, loss of balance, excessive freezing, erratic movement, or failure to explore the chamber during the baseline period. Observe the fish briefly under stable conditions and confirm that the water level, temperature, lighting, and handling conditions are appropriate. Only fish showing healthy appearance and normal baseline behavior should be used for the experiment. If the fish continues to appear stressed or abnormal, remove it according to the laboratory’s animal-handling procedures, document the observation, and replace it with another fish rather than continuing the trial.

**Problem 8:** Fish show little or no response during the training period

**Solution:**

(1) If the fish show little or no visible response during the shock-pairing period, first confirm that the stimulation system passed voltage calibration before the experiment. The measured voltage across each positive–negative electrode pair should be approximately 9.4–9.5 V, with the external power supply set to 9.50 V and the current limit set to 0.200 A. If calibration was not passed, do not interpret the behavioral result and do not proceed until the wiring and stimulation circuit have been checked.
(2) Next, check the physical and experimental conditions. The water level should be within the required range, with a target depth of 3.0 cm and below 3.5 cm. Water that is too shallow may affect electrode contact. Also confirm that the water temperature is within the normal experimental range and that the fish were handled consistently before testing. Feeding state, prolonged fasting, handling stress, or anxiety-like behavior may affect how strongly fish respond during training.
(3) The electrical circuit should also be inspected. Confirm that the relay activates during the programmed shock periods, the alligator clips are firmly attached to the correct electrode rods, and the positive and negative electrode pathways are not loose or accidentally disconnected. If the optional reference LED is used, it should illuminate during programmed shock periods, indicating that the stimulation circuit is active. If the reference LED does not light, check the power supply output, relay wiring, electrode connections, and resistor placement.

**Important**. If the stimulation circuit is working properly but the fish still show weak responses, document the observation rather than increasing the voltage arbitrarily. Any change to shock voltage, current limit, or training intensity should be treated as a protocol modification and should follow the approved experimental and animal-care guidelines.

**Problem 9:** Fish jumps out of the chamber or crosses into the other chamber

**Solution:**

Stop the Arduino protocol and video recording immediately. Remove or recover the fish according to the laboratory’s animal-handling procedures, and document the event, including the time point, chamber, fish ID, and possible cause. The affected trial should be excluded from downstream behavioral analysis. **Do not test the same fish again** in a new trial. Because the long-term behavioral effects of the training protocol have not been fully characterized, reusing the same fish may introduce learning, stress, or memory-related carryover effects.

**Problem 10:** Cannot connect to OBS

**Solution:**

(1) Possible causes: OBS is not open or WebSocket server is disabled. Open OBS.
(2) Check ‘WebSocket Server Settings’ in ‘Tools’, and confirm that ‘Enable WebSocket Server’ is selected and the ‘Server Port’ is 4455.

**Problem 11:** Camera does not show video feed

**Solution:**

(1) Reconnect the camera USB cable and confirm that the camera is recognized by the computer. In the recording software, select the correct camera source and restart the software if needed. If the camera still does not appear, try a different USB port, check whether another program is already using the camera, and restart the computer if necessary.
(2) An infrared-sensitive camera is required for recording during dark periods. If the camera feed appears too dark even when the infrared mode is active, additional infrared illumination may be needed. Any added infrared light source should be positioned so that it improves visibility without blocking the top-down view, creating strong reflections, or interfering with the arena. Visible room light should not be used during the experiments.
